# Microbial mechanistic requirements for eliciting a topical and intranasal immune response

**DOI:** 10.64898/2026.07.15.738787

**Authors:** Katherine D. Bauman, Pranav V. Lalgudi, Djenet Bousbaine, Anchaleena James, Miki Chiang, Tam T.D. Nguyen, Joyce M. Swenson, Eunice Tsang, Hyojin Kim, Felicitas Ruiz, Naveen Jasti, Zachary Jones, David B. Li, Annie T. Nguyen, Anne Trinh, Megha Lingamsetty, Aishan Zhao, Masahiko Terasaki, Neil P. King, Michael A. Fischbach

**Affiliations:** Department of Bioengineering, Stanford University, Stanford, CA, USA; Department of Genetics, Stanford University School of Medicine, Stanford, CA, USA; ChEM-H Institute, Stanford University, Stanford, CA, USA; Microbiome Therapies Initiative, Stanford University, Stanford, CA, USA; Department of Biochemistry, Stanford University, Stanford, CA, USA; Department of Biochemistry, University of Washington, Seattle, WA, USA; Institute for Protein Design, University of Washington, Seattle, WA, USA; Department of Molecular Engineering, University of Washington, Seattle, WA, USA; Chan Zuckerberg Biohub, Stanford, CA, USA

**Author notes:** These authors contributed equally (Katherine D. Bauman, Pranav V. Lalgudi). These authors contributed equally (Anchaleena James, Miki Chiang, Tam T.D. Nguyen).

## Abstract

The skin colonist *Staphylococcus epidermidis* elicits a potent antibody response that can be redirected against an antigen of interest, but this process relies on genetic engineering.^1^ Here, by adapting bioorthogonal chemistry methods to conjugate antigens to the cell surface, we make the process of generating a commensal vaccine rapid and efficient. A wide variety of bacteria displaying tetanus toxin fragment C (TTFC) elicit an antibody response when applied to mice topically, indicating that the inductive process is not limited to colonists. Colonization occurs at two different sites, the skin and nostril; by colonizing each with *S. epidermidis-*TTFC, we show that skin colonization yields a moderate IgG response, while nostril colonization elicits a highly potent systemic IgG response and an exuberant IgA response in the nostrils, lungs, and intestine. Two lines of evidence are consistent with the nasal-associated lymphoid tissue (NALT) as the inductive site for nostril colonization: imaging suggesting robust bacterial translocation, and the induction of commensal-specific B cells following colonization. On the skin, TTFC must be conjugated to live *S. epidermidis* to elicit an antibody response; in the nostrils, live *S. epidermidis*-TTFC and *S. epidermidis* mixed with TTFC are equally potent. Commensal vaccination yields a robust response in pet shop mice, and chemical conjugation facilitates antibody responses to a broad array of antigens, including a whole viral capsid and a rotavirus immunogen. Collectively, these findings provide compelling evidence of a translational path for commensal vaccines.

## INTRODUCTION

In recent work, we reported that the skin colonist *Staphylococcus epidermidis*, like commensals at other barrier sites,^2–7^ elicits an antibody response under conditions of physiologic colonization,^8^ and that this response can be redirected against non-native antigens genetically expressed on the cell surface.^1^ However, this work raises three important questions about the mechanistic features of this immune response: (*i*) Do other skin colonists elicit a similar response when engineered? (*ii*) What are the mechanistic requirements on the microbial side for eliciting a response? (*iii*) How broadly will this approach work for antigens beyond TTFC?

A key finding in our previous study was that a strain of *S. epidermidis* engineered to express SpyCatcher on its surface and then attached to a recombinant, SpyTag-bearing antigen elicited a robust antibody response upon colonization, demonstrating that continuous antigen production is not required for commensal-induced immunity. Here, guided by this observation, we adapt widely used bioorthogonal chemistry methods to conjugate immunogens to the bacterial cell surface.^9–15^ This approach eliminates the need for genetic engineering, works for a wide variety of antigens, and is cheap, easy, and efficient, and therefore broadly accessible to the research community.

We show that chemical conjugation results in efficient labeling of the bacterial cell surface, and *S. epidermidis* labeled in this way with the immunogen tetanus toxin fragment C (TTFC) elicits a highly potent antibody response. We then use chemical labeling to investigate the microbial requirements for eliciting this response. A wide variety of bacterial species can be labeled with antigen; most of them, regardless of origin, induce an antibody response when conjugated to TTFC, indicating that the inductive mechanism is not restricted to colonists. We find that there are two distinct pathways for sampling: one that operates through the skin and results in a systemic IgG response, and another that functions intranasally and results in an exuberant systemic and mucosal immune response in the nares, lungs, and intestine. Microbiologic requirements differ in each case, suggesting distinct mechanisms of sampling.

Finally, we demonstrate that chemically labeled *S. epidermidis* elicits a potent antibody response in pet shop mice, and we show that commensal vaccination works for a variety of antigens, including whole viral capsids.

### Adapting bioorthogonal chemistry to the *S. epidermidis* cell surface

We set out to determine whether the surface of *S. epidermidis* could be labeled with widely used bioorthogonal chemistries. We sought to test two attachment strategies: direct attachment of a dye-linked electrophile, and a two-step strategy in which a chemical handle is installed on the cell surface and then a payload is attached (**Figs. 1A, 1B**). In each case, we labeled the human skin colonist *S. epidermidis* LM088 *in vitro*, monitoring the extent of labeling using flow cytometry (**Fig. 1C**). We found that a range of techniques worked efficiently: amine-reactive crosslinkers, thiol-reactive crosslinkers, and metabolic labeling facilitated attachment of the fluorescent dye Cy5 to the surface of *S. epidermidis*.^13^ Azide/alkyne cycloaddition^9,16^ and the biotin/streptavidin^14^ interaction were both capable of linking payloads to a handle installed on the cell surface (**Figs. 1C, 1D**). After some optimization, we found that attachment to primary amines via *N-*hydroxysuccinimide (NHS) esters proved especially efficient and versatile^17,18^ (**Figs. S1, S2**); we prioritized this chemistry for the remainder of the study, though we expect other chemistries will prove useful in future efforts.

**Figure 1.**
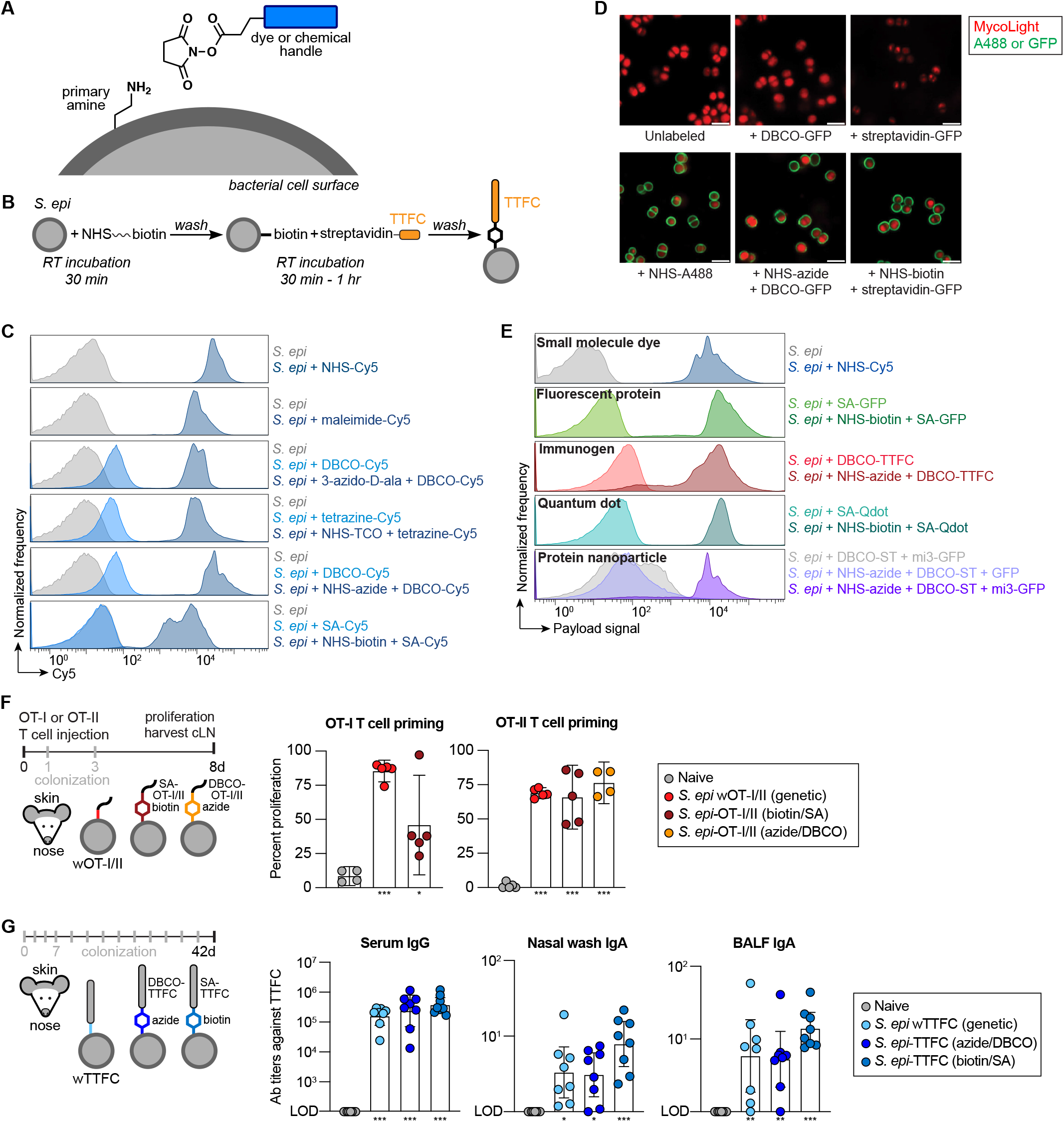
Chemical conjugation of antigens to *S. epidermidis* induces immune responses *in vivo*. (A) Schematic showing chemical conjugation of payloads to the bacterial surface using amine-reactive crosslinkers. (B) Schematic outlining the experimental steps for chemically conjugating antigens to the bacterial cell surface. NHS-PEG_n-_biotin or NHS-PEG_n-_azide were mixed with bacteria and incubated for 30 min. Labeled cells were washed and then incubated with streptavidin (SA) conjugates or dibenzocyclooctyne (DBCO)-derivatized proteins for 30 min-1 h. Functionalized cells were washed and then analyzed *in vitro* or used to colonize mice. (C) Various chemistries facilitate dye attachment to the *S. epidermidis* cell surface. Flow cytometry plots showing *S. epidermidis* conjugated to the small-molecule dye Cy5 using amine-reactive crosslinkers (NHS ester), thiol-reactive crosslinkers (maleimide), and metabolic labeling (3-azido-D-alanine). A two-step labeling procedure can also be used to attach Cy5 to chemical handles on the bacterial cell surface using NHS-biotin + streptavidin (SA) conjugates, NHS-azide + DBCO-tethered molecules, or NHS-TCO + tetrazine-linked molecules. NHS = N-hydroxysuccinimide ester, DBCO = dibenzocyclooctyne, TCO = *trans-*cyclooctene. (D) Confocal microscopy images of *S. epidermidis* conjugated to AlexaFluor488 or GFP (green) using NHS ester chemistry. Bacterial cells identified as MycoLight^+^ (red). Scale bar = 2 µm. (E) A variety of payloads can be attached to the cell surface. Flow cytometry plots showing *S. epidermidis* conjugated to Cy5, GFP, the immunogen TTFC, quantum dots, and mi3 protein-based nanoparticles. (F) Colonization with chemically conjugated *S. epidermidis* induces the proliferation of adoptively transferred OT-I or OT-II T cells. OT-I or OT-II T cells were isolated from CD45.2 *Rag2^-/^*^-^donor mice, stained with CellTrace Violet dye, and injected retro-orbitally into CD45.1 SPF mice. Recipient mice were colonized on days 1 and 3 after transfer with *S. epidermidis* LM087 strains either genetically engineered to express or conjugated to OT-I or OT-II peptides. On day 8, cervical lymph nodes were harvested and analyzed by flow cytometry for T cell proliferation. Data are shown as frequency of Live Tcrb^+^ CD8^+^ CD4^-^CD45.2^+^ CD45^-^Vb5^+^ Va2^+^ for OT-I cells or Live Tcrb^+^ CD4^+^ CD8^-^CD45.2^+^ CD45^-^Vb5^+^ Va2^+^ for OT-II cells; proliferation is quantified by the dilution of CellTrace Violet. (G) *S. epidermidis* bearing chemically linked TTFC elicits a potent antibody response. Antibody endpoint titers from mice colonized with *S. epidermidis* LM088 either genetically engineered to express or chemically conjugated to TTFC. Mice were colonized 13 times over six weeks, as previously described.^1^ Serum, nasal wash, and bronchoalveolar lavage fluid (BALF) were harvested and assayed by ELISA for TTFC-specific antibodies. Data shown are pooled from two independent experiments. Data are from one (**F**), or pooled from two (**G**), or representative of at least three (**C**, **D**, **E**) independent experiments. Graphs show mean (**F**) or geometric mean (**G**) with a 95% confidence interval. *P* values (ns, not significant; * *p* < 0.05; ** *p* < 0.01; *** *p* < 0.001) were calculated using an ordinary one-way ANOVA followed by Tukey’s multiple comparisons test, assuming normal (**F**) or log-normal (**G**) distributions. Asterisks in figures represent pairwise comparisons of the indicated group to naive mice; *p* values for all comparisons are reported in **Table S1**.

Next, we sought to determine what payload types could be attached in this manner. First, we treated *S. epidermidis* with NHS-PEG_4-_biotin (NHS-biotin), and then we added two different payloads: a recombinant streptavidin-GFP fusion protein (SA-GFP) or a streptavidin-linked quantum dot (SA-Qdot). In a similar fashion, we labeled cells with NHS-PEG_4-_azide (NHS-azide) and utilized two additional payloads: TTFC labeled with the electrophilic alkyne NHS-dibenzocyclooctyne (DBCO-TTFC) or DBCO-SpyTag003; to the latter, we added the SpyCatcher003-based protein nanoparticle mi3 displaying GFP. All four of the candidate payloads were efficiently attached (**Figs. 1E, S2**), yielding a set of strains that are equipped to interrogate various aspects of the immune response to topical colonization.

### Chemically labeled *S. epidermidis* elicits a highly potent immune response

Although we observed in the previous study that SpyCatcher-linked antigens were immunogenic on the *S. epidermidis* cell surface, we did not know whether chemically linked antigens—which are attached to different sites on the cell surface, in different orientations, by non-peptidic linkers—would be competent to induce an immune response. To address this question, we performed two immunogenicity experiments: one to test T cell priming, and the other to assess B cell activation.

First, we used an adoptive transfer assay to assess the ability of *S. epidermidis* bearing a chemically linked T cell epitope to prime cognate T cells (**Fig. 1F**). We adoptively transferred ovalbumin (OVA)-specific CD8^+^ and CD4^+^ T cells (OT-I and OT-II, respectively) into specific pathogen-free (SPF) C57BL/6 mice. We then colonized these mice topically with *S. epidermidis* bearing a matched, chemically linked class I or II epitope (*S. epidermidis*-OT-I, *S. epidermidis-*OT-II) by applying the labeled bacterial strains with a cotton swab onto the head and ears of the mice, collectively referred to as the skin, and tapping the entrance of the nostrils to colonize the nose. After eight days, we quantified the extent of T cell proliferation by flow cytometry. OVA-specific CD8^+^ and CD4^+^ T cells proliferated in response to colonization with *S. epi*-OT-I and *S. epi-*OT-II (respectively), indicating that chemically linked T cell epitopes are accessible to the immune system.

Second, we used two conjugation chemistries to link a recombinant B cell immunogen, TTFC, to the cell surface: DBCO-TTFC was linked to azide-bearing *S. epidermidis*, as described above, and a recombinant streptavidin-TTFC (SA-TTFC) fusion protein was linked to biotin-bearing *S. epidermidis* (*S. epidermidis-*TTFC). Both techniques yielded highly efficient cell surface labeling, as evidenced by flow cytometry and microscopy (**Figs. S1G, S2C, S2E**)

We then colonized SPF C57BL/6 mice topically on the skin and nose with these labeled cells following our earlier colonization schedule (**Fig. 1G**).^1^ After six weeks, we measured the titer of IgG against TTFC in systemic circulation (i.e., serum), and the titer of IgA against TTFC in nasal wash and bronchoalveolar lavage (BAL) fluid. Both strains of *S. epidermidis-*TTFC elicited robust antibody responses, at least as potent as the genetically engineered strain developed in our prior study.

### Diverse bacterial species can be labeled

One of the main advantages of attaching antigen chemically instead of encoding it genetically is that it could, in principle, work on a variety of organisms. To test this premise, we assembled a panel of 57 bacterial species that included Gram-positive and Gram-negative organisms; aerobic and anaerobic; and commensals, pathogens, and environmental species. At least a quarter of these strains are considered genetically inaccessible.

Each strain was treated with NHS-PEG_4-_biotin and then incubated with recombinant streptavidin-GFP, and labeling efficiency was assessed by flow cytometry in comparison to a control reaction lacking NHS-PEG_4-b_iotin (**Fig. 2A**). 54/57 strains were successfully labeled, showing that labeling works robustly independent of phylogeny or origin. Notably, all three strains that failed to be labeled are known to produce a surface polysaccharide,^19–22^ suggesting that the only requirement for labeling is a suite of surface-accessible primary amines. We noted differences in labeling efficiency across strains; for example, *Pasteurella multocida* required a longer PEG linker between NHS and biotin. This technique opens a path to manipulating a wide variety of host-adapted and environmental bacterial species to help probe the mechanisms of interspecies interactions.

**Figure 2.**
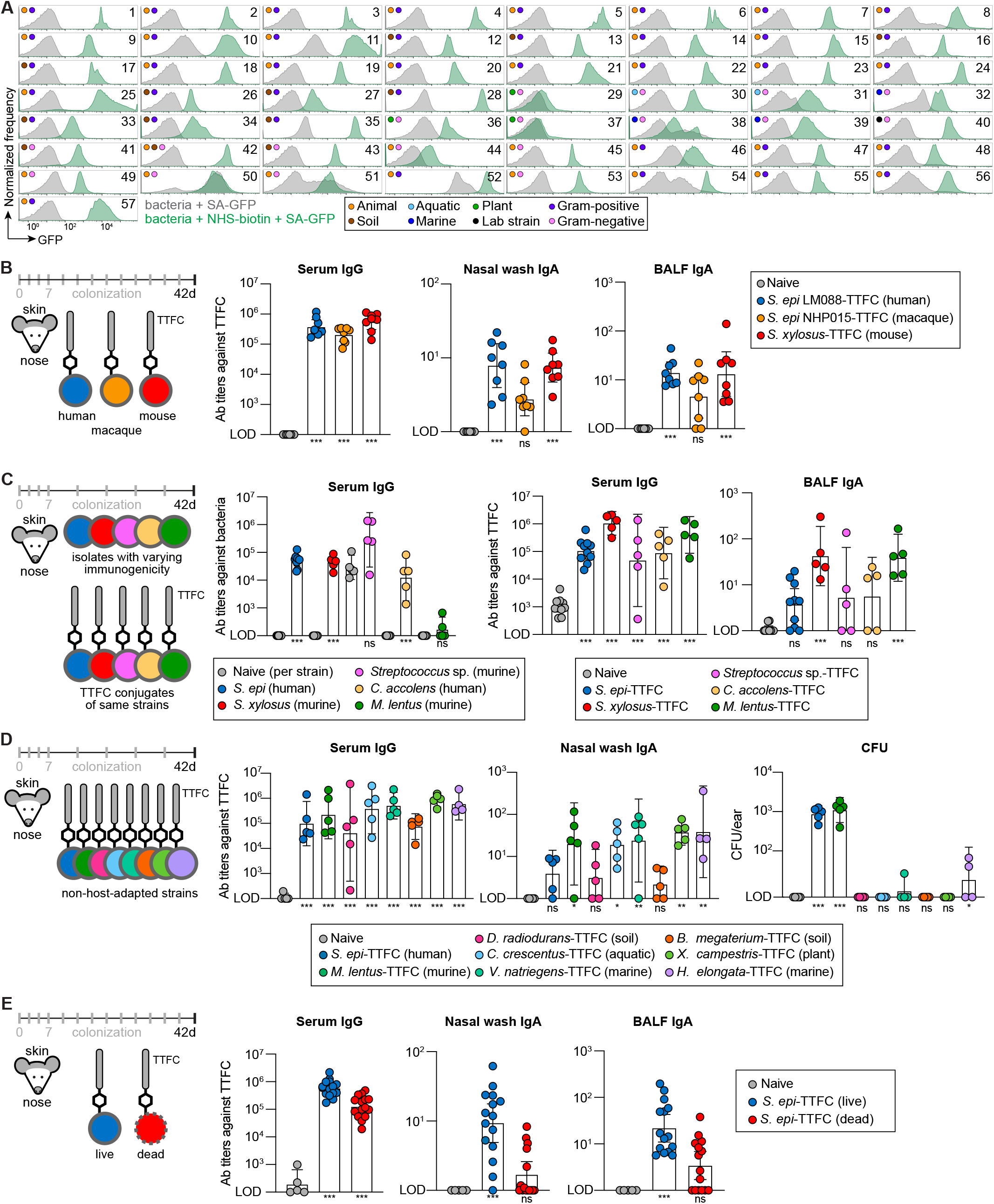
A wide variety of bacterial species elicit an antibody response when applied to mice topically. (A) Flow cytometry plots showing GFP conjugated to a panel of phylogenetically diverse bacteria that are associated with animal or plant hosts or are free-living in soil, marine, and aquatic environments. Numbers refer to strains listed in **Table S2**. (B) *Staphylococcus* isolates from human, macaque, or murine hosts elicit comparably potent antibody responses. Strains were conjugated to TTFC and used to colonize the skin and nose of mice 13 times over six weeks. TTFC-specific antibodies were quantified in serum, nasal wash, and BAL fluid by ELISA and are depicted as antibody endpoint titers. Data shown are pooled from two independent experiments. Data from the naive and *S. epidermidis* LM088-TTFC groups are from the same experiment as Fig. 1F; all mice were colonized and analyzed concurrently. Bacterial CFU recovered from the colonized mice are shown in **Fig. S4**. (C) Regardless of native immunogenicity, colonists elicit a uniformly high response against a heterologous antigen. Human and murine isolates with varying degrees of immunogenicity were assayed in two ways. First, wild-type strains were used to colonize the skin and nose of SPF mice eight times over six weeks; the graph on the left shows antibody titers from the serum of naive (gray) or colonized mice (colored) against the bacterial strain itself. Second, the same strains were conjugated to TTFC and used to colonize mice in the same way. At the experimental endpoint, serum, BAL fluid, and nasal wash (shown in **Fig. S5**) were assayed by ELISA for TTFC-specific antibodies. (D) Non-host-adapted strains are comparably immunogenic to skin colonists. Six non-host-associated bacterial strains (i.e. environmental isolates) were conjugated chemically to TTFC and used to colonize the skin and nose of SPF mice eight times over six weeks. Antibody titers against TTFC were measured by ELISA in serum, nasal wash, and BAL fluid (shown in **Fig. S6**). Bacteria were recovered from mouse ears and quantified by plating on their respective growth media. Naive CFU shown is from BHI media, and all statistical comparisons are to naive (BHI) (additional growth media and associated statistics shown in **Fig. S6** and **Table S1**). LOD = 6.67. (E) Live cells elicit a stronger response than dead cells. Heat-killed or live *S. epidermidis* was attached to TTFC and applied to the skin and nose of mice 13 times over six weeks. At the experimental endpoint, serum, nasal wash, and BAL fluid were collected and assayed for antibodies against TTFC by ELISA. Data shown are from one (**D**), pooled from two (**B**) or representative of two (**A, C, E**) independent experiments. Graphs show geometric mean with a 95% confidence interval. *P* values (ns, not significant; * *p* < 0.05; ** *p* < 0.01; *** *p* < 0.001) were calculated using an ordinary one-way ANOVA followed by Tukey’s multiple comparisons test, assuming log-normal distributions except for bacterial ELISAs in **C** and **D**. For bacterial ELISAs in **C,** log-transformed titers from naive and colonized mice were compared for each strain using unpaired two-tailed *t* tests with the Holm-Šídák correction for multiple comparisons (a = 0.05). Asterisks in figures represent pairwise comparisons of the indicated group to naive mice; *p* values for all comparisons are reported in **Table S1**.

### *Staphylococcus* species elicit an antibody response independent of origin

Next, we sought to determine the mechanistic requirements for eliciting a B cell response on the microbial side. As a starting point, we performed a series of experiments in which the same antigen was displayed on different bacterial species. In other vaccination settings, the host origin of the carrier is known to impact efficacy; for example, human adenoviruses can fail to elicit an antibody response due to pre-existing immunity in humans, so chimpanzee-derived adenoviruses are often used instead.^23,24^ Moreover, there is a considerable degree of host-colonist co-evolution^25–27^ since our previous studies involved colonizing mice with a human isolate of *Staphylococcus*, we sought to determine whether the host of origin impacts the ability of a strain to elicit an antibody response in mice.

To this end, we labeled three strains of *Staphylococcus* with streptavidin-TTFC—*S. epidermidis* LM088 (human), *S. epidermidis* NHP015 (macaque), and *S. xylosus* (mouse)—and used equivalent quantities of the labeled bacteria to colonize mice (**Figs. 2B, S4**). After six weeks, we measured IgG titers in serum and IgA titers in nasal wash and BAL fluid. Antibody titers were comparably potent for all three strains, indicating that—at least among these isolates—the host of origin does not impact the ability to elicit a response. We recovered comparable amounts of each strain from the skin on day 42, suggesting that they colonize with similar efficiency.

### Immune responses against native vs. conjugated antigens

Next, we asked whether the ability of a wild-type (non-engineered) strain to elicit a B-cell response against its native surface antigens is correlated with its ability to direct an immune response against non-native antigens on its surface. From a broader screening effort, we assembled a small panel of skin isolates that vary widely in the degree of antibody response they elicit against their native surface antigens. On one end of the spectrum, *S. epidermidis* LM088 and *Staphylococcus xylosus* (the primary murine *Staphylococcus* colonist) elicit very strong antibody responses against their own antigens; on the other end, the human isolate *Corynebacterium accolens* and the murine isolate *Mammaliicoccus lentus* elicit a weak response or none at all, respectively.^1^ Each strain was labeled with streptavidin-TTFC and equal quantities of each strain were used to colonize mice (**Figs. 2C, S5**).

To our surprise, we found that regardless of their native immunogenicity, every strain elicited a uniformly high response against the heterologous antigen. This was particularly notable in the case of *M. lentus*; this strain colonizes robustly but elicits no antibody response on its own. However, when linked to TTFC, it generates a very strong response against the chemically attached protein, indicating that while it is efficiently sampled, native antigens are absent or hidden. Importantly, *M. lentus-*TTFC does not elicit a response against the endogenous *M. lentus* surface antigens, suggesting that the process of sampling the heterologous antigen does not unmask native antigens.

### Non-host-adapted strains are comparably immunogenic to skin colonists

We had assumed that the ability of a strain to persist on a host barrier surface was inherently tied to its ability to elicit an immune response; as a result, every strain we had assayed was a native skin or nasal colonist. To test this assumption, we conjugated TTFC to a group of phylogenetically diverse non-host-adapted strains isolated from soil, plants, or aquatic ecosystems. Notably, many of these strains are Gram-negative, whereas the predominant colonists of skin are Gram-positive (*Staphylococcus, Corynebacterium,* and *Cutibacterium*).^28,29^ Additionally, several species we tested have strict growth requirements that are not met on a mammalian host.^30,31^

When we colonized the skin and nose of mice with the same quantity of each labeled strain, we were surprised to see that every strain we tested elicited a response against TTFC that was comparable to two colonists included as controls, *S. epidermidis*-TTFC and *M. lentus-*TTFC (**Figs. 2D, S6**). Moreover, the responses elicited by the Gram-negative species *Vibrio natriegens*, *Xanthomonas campestris*, and *Halomonas elongata* were particularly striking, especially for their ability to induce mucosal IgA. This is notable in light of our inability to recover these strains from the skin of colonized mice, consistent with their lack of adaptation to mammalian hosts. Taken together, these findings suggest that the inductive process is not specific to colonists.

### Live cells elicit a stronger response than dead cells

Given that every bacterial strain we tested could elicit an immune response, we next asked whether immunogenicity requires bacterial viability. In prior work, we showed that there is a requirement for *S. epidermidis* to be alive in order to elicit a T cell response,^32^ but this has been difficult to probe in the context of the B cell response because the process of heat killing could denature the immunogen.

To address this question, we began by heat-killing *S. epidermidis* LM088 and then subsequently labeled it with an immunogen. As shown in **Fig. S7**, TTFC can be conjugated efficiently to heat-killed *S. epidermidis* in a manner that maintains the cell-surface disposition of the antigen, enabling us to compare its immunogenicity to live cells labeled with the same immunogen. By using live and dead *S. epidermidis-*TTFC to colonize mice, we find that the live cells elicit a much stronger response than the dead cells both systemically and mucosally, indicating that viability increases the potency of commensal vaccination (**Fig. 2E**).

### Antibody response differs on the skin and in the nostril

Having determined the coarse cellular requirements for topical immunogenicity (the ‘what’), we turned next to the ‘where’: does the site of colonization impact the nature of the antibody response?

Our standard method of colonization consists of running a cotton swab over the head and ears of the mice, including tapping the entrance of the nostril. *S. epidermidis* is a ubiquitous colonist of the skin and nostrils, so it is naturally found in both sites.^28,29^ However, the skin is a keratinized squamous epithelium, whereas the nostril transitions to a columnar respiratory epithelium lined with mucus. Reasoning that the mechanisms of sampling and immune sequelae might differ between these sites, we set out to characterize the immunologic outcomes of skin versus nostril colonization.

We colonized one group of mice with *S. epidermidis*-TTFC on the skin of the head and ears only, avoiding the face and the nostrils; a second group in the nostrils only; and a third group the same way we originally performed colonization, encompassing both sites. As before, concurrent colonization of the skin and nostrils led to a robust induction of IgG against TTFC in circulation and IgA in the nasal and pulmonary mucosa (**Figs. 3A, S8A**).

**Figure 3.**
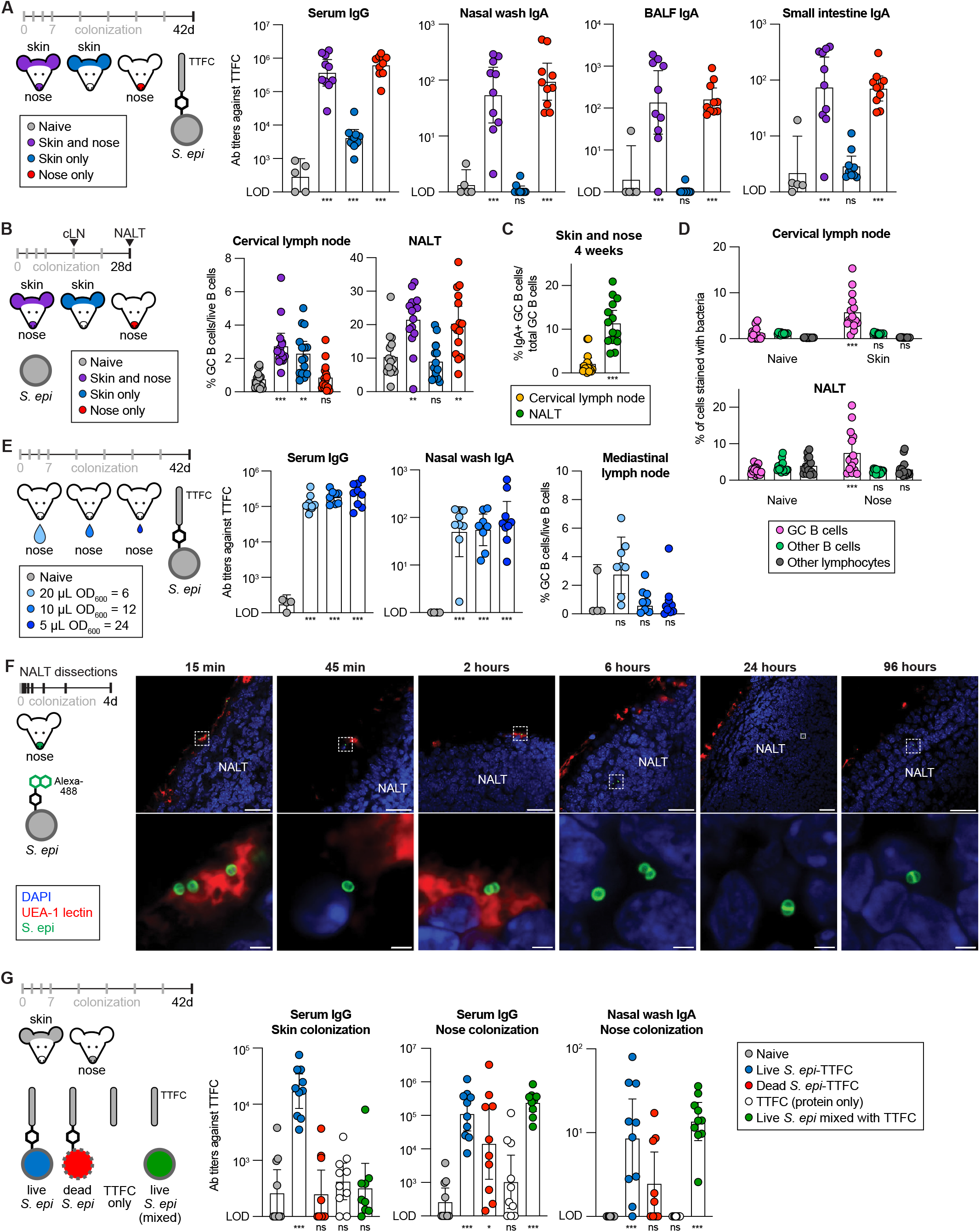
Antibody responses to *S. epidermidis* on the skin and in the nostril. (A) Antibody responses elicited by skin and nostril colonization are markedly different. Mice were colonized in the nose, on the skin, or in both sites with *S. epidermidis*-TTFC. After eight colonizations over six weeks, serum, nasal wash, BAL fluid and small intestinal contents were harvested and analyzed for TTFC-specific antibodies by ELISA. (B) The cervical lymph nodes and NALT serve as the inductive sites for skin and nasal colonization, respectively. Mice were colonized with wild-type *S. epidermidis* on the skin, in the nostril, or in both sites. Cervical lymph nodes were harvested after two weeks and NALT was harvested after four weeks. At each site, the frequency of germinal center B cells (CD38^-^Fas^+^) among live B cells (Live CD45.2^+^ CD19^+^) was assessed by flow cytometry. (C) IgA-expressing B cells derive from the NALT. The percentage of germinal center B cells that were IgA+ (IgA^+^ IgG1^-^) from mice colonized on the skin and nose was quantified by flow cytometry. Animals are from the same experiment shown in Fig. 3B. (D) Germinal center B cells are *S. epidermidis*-specific. Cell suspensions from cervical lymph nodes (naive or skin-colonized mice) and the NALT (naive or nostril-colonized mice) were incubated with Alexa488-conjugated *S. epidermidis* to quantify the fraction of germinal center B cells (Live CD45.2^+^ CD19^+^ CD38^-^Fas^+^) that were bacteria-specific, compared to other B cells (Live CD45.2^+^ CD19^+^ CD38^+^ Fas^-^), or other lymphocytes (Live CD45.2^+^ CD19^-^). Animals are from the same experiment shown in Fig. 3B. (E) Nasal colonization by *S. epidermidis*-TTFC can elicit an immune response without mediastinal involvement. The same number of *S. epidermidis*-TTFC cells was administered intranasally by pipetting three different volumes (as indicated, µL per nostril). After eight colonizations over six weeks, serum and nasal wash were analyzed for TTFC-specific antibodies by ELISA. At the experimental endpoint, mediastinal lymph nodes were collected, and the percentage of germinal center B cells (CD38^-^Fas^+^) among live B cells (Live^+^ CD45.2^+^ CD19^+^) was assessed by flow cytometry. (F) Time course of bacterial localization in the NALT after colonization. Mice were colonized intranasally with fluorescent *S. epidermidis* conjugated to AlexaFluor488. NALT was harvested from groups of mice at various time points after colonization (15 minutes, 45 minutes, 2 hours, 6 hours, 24 hours, and 96 hours) and flash frozen. Tissues were sectioned (100 µm), stained, and subsequently imaged by confocal microscopy (blue = DAPI, red = UEA-1 lectin, green = *S. epidermidis*). The bottom row depicts enlarged images outlined by a white dashed box (top row). Scale bars shown are 25 µm (top) or 2 µm (bottom). More images and time points are shown in **Fig. S10**. (G) Antigen must be attached to a live bacterium to induce an immune response on the skin, whereas antigen conjugated to or mixed with live bacteria can elicit antibodies intranasally. Mice were colonized by swabbing either the skin or nose with one of four different test articles (live *S. epidermidis* conjugated to TTFC, dead *S. epidermidis* conjugated to TTFC, TTFC alone, or *S. epidermidis* mixed with TTFC). The same amount of antigen was compared between each of these formulations, and three different doses were tested at each body site (**Fig. S12**). Mice were colonized eight times over six weeks, after which serum and nasal wash were collected and assessed for TTFC-specific antibody titers by ELISA. Antibody titers shown here are from mice colonized with the high dose of antigen (skin colonization) or medium dose of antigen (nose colonization). Data from both body sites and all dosages are shown in **Fig. S12**. Data shown are representative of at least two independent experiments. Graphs show mean (**B, C, D**) mediastinal lymph node in **E** or geometric mean (**A, E, G**) with a 95% confidence interval. *P* values (ns, not significant; * *p* < 0.05; ** *p* < 0.01; *** *p* < 0.001) were calculated using an ordinary one-way ANOVA (**A, B, C, E, G**) or a two-way ANOVA (**D**) followed by Tukey’s multiple comparisons test, assuming normal (**B, C, D**, mediastinal lymph node in **E**) or log-normal (**A, E, G**) distributions. Two-way ANOVA for **G** comparing test articles across all dose levels and subsequent statistical testing is shown in **Fig. S13**. Asterisks in figures represent pairwise comparisons of the indicated group to naive mice (**A, B, E, G**), to cervical lymph nodes (**C**), or to the corresponding cell type from naive mice (**D**); *p* values for all comparisons are reported in **Table S1**.

Colonization of each site alone gave markedly different responses. Mice that were colonized on the skin only had a systemic IgG response of moderate strength and no IgA at mucosal surfaces. In contrast, colonization of the nostrils led to a highly potent IgG response and robust induction of IgA in the nasal wash and BAL fluid. Having seen a strong IgA response in the nose and lungs, we collected and assayed the contents of the small intestine, revealing a marked induction of IgA against TTFC in this site as well. Notably, intestinal IgA was not observed when we gavaged mice orally with the same strain, consistent with a model in which intestinal IgA derives from mucosal spreading rather than local induction in the intestine (**Fig. S8B**).

### Compartmentalization of the germinal center B cell response

Given the functional differences in the antibody response to skin versus nostril colonization, we set out to identify the lymphoid tissues from which the antibodies derive. We colonized mice in the same way, separating skin from nose, and collected three lymphoid tissues: the cervical lymph node (cLN), which drains the skin of the ears of the mouse; the nasal-associated lymphoid tissue (NALT), which drains the nasal passage^33–36^ and the inguinal lymph node, which drains skin sites that were not colonized (back and groin).^37^ We saw an induction of germinal center (GC) B cells in the cervical lymph node in mice colonized on the skin but not intranasally (**Figs. 3B, S8C**). In contrast, GC B cells were present in the NALT in mice colonized intranasally but not on the skin. There was no induction of GC B cells in the inguinal lymph node in either group of mice. These data suggest that the germinal centers giving rise to the skin and nostril antibody responses are compartmentalized in separate sites: the cLN and NALT, respectively. Moreover, in mice colonized on the skin and nose, IgA^+^ GC B cells were abundant in the NALT but not the cLN (**Figs. 3C, S8D**), indicating that the IgA response derives from the NALT.

Next, we sought to assess whether the germinal center reaction was specific to *S. epidermidis.* Canonically, antigen-specific B cells are detected with dye-labeled protein multimers, which can be onerous to construct.^38–40^ We reasoned that dye-labeled *S. epidermidis* cells*—*which harbor surface antigens at a high valency—might stain B cells expressing an *S. epidermidis-*specific BCR sensitively and specifically. First, we colonized mice with *S. epidermidis* LM088, prepared cell suspensions from harvested lymphoid tissues, and stained them with *S. epidermidis* LM088 that had been labeled with NHS-AlexaFluor488 (**Figs. 3D, S9**). By titrating the number of bacterial and cLN cells in the mixture, we identified a ratio at which we clearly stain germinal center B cells from colonized mice but not naïve mice. Three lines of evidence suggest that this staining is specific: when we block with excess unlabeled bacteria, signal drops (**Fig. S9F**); when we colonize transgenic mice that only express an unrelated BCR, we observe no staining above naïve background (**Fig. S9G**); and when we stain germinal center B cells from mice colonized with LM088 using three other fluorescently labeled strains (LM088 Δ*aap*, which is missing the immunodominant antigen; *S. epidermidis* LM087; or *M. lentus*), we do not observe staining above background (**Fig. S9H**). Using this method to analyze germinal center B cells from the cLN and NALT, we observe *S. epidermidis-*specific B cells when we colonize the skin and nose, respectively, indicating that both pathways lead to an antigen-specific response.

### Most of the antibody response derives from the NALT

Under certain conditions, liquid administered into the nasal passage can drain to the lungs, involving the mediastinal lymph node.^41–43^ To determine whether the mediastinal lymph node is involved in the antibody response we observed following nasal colonization, we carried out three experiments. (*i*) We performed a volumetric titration using a pipet to administer different volumes of *S. epidermidis-*TTFC into the nasal vestibule of awake mice, using our swab-based colonization method as a control (**Fig. S10A**). Tapping and pipetting high volumes (30 µL per nostril and above) led to a modest induction of GC B cells in the mediastinal lymph node. However, a lower volume (20 µL per nostril) led to robust induction of IgG and IgA with little to no mediastinal involvement. (*ii*) We administered a similar volume of dye using the same methods, revealing that 20 µL of dye per nostril in mice that were awake led to no visible staining in the lungs (**Fig. S10C**). (*iii*) Having observed that 20 µL of bacteria per nostril—but not less—elicits a potent antibody response, we wondered whether this effect is due to the volume applied or the antigen dosage (**Figs. 3E, S10B**). To test this, we suspended an equivalent quantity of *S. epidermidis-*TTFC in three different volumes: 20 µL of OD_600=_6 (our standard concentration), 10 µL of OD_600=_12, and 5 µL of OD_600=_24. All three treatments elicited a highly potent IgG and IgA response, consistent with the view that the strength of the response is driven by the quantity of *S. epidermidis-*TTFC and not by the volume of liquid in which it is delivered. Taken together, these data are consistent with the view that most of the antibody response derives from germinal centers in the NALT, though we cannot rule out the involvement of other lymphoid tissues. Moreover, these findings have important practical applications, as they demonstrate that a very small volume of bacterial culture can elicit a highly potent response.

### Features of the immune response to skin colonization

Previous work has implicated Langerhans cells as the primary antigen-presenting cell for the immune response to skin colonization with *S. epidermidis*.^8,44,45^ This response proceeds via inductive sites in tertiary lymphoid organs in the skin as well as in skin-draining lymph nodes.^1,8^ We find that many skin sites on the mouse are competent to induce immunity, regardless of their thickness or hair follicle density (**Fig. S11A**). The responses in skin-draining lymph nodes are compartmentalized: the inguinal lymph node appears to be the inductive site for colonization of the lower back (**Fig. S11B**), whereas the cLN drains the skin on the head and ears of the mouse.

A gentle method of application—spraying bacterial culture onto the back of a mouse using a mucosal atomization device—elicited an equivalent immune response to our standard swab-based application. This suggests that the sampling mechanism is not dependent on any feature of the swabbing process, including potential micro-abrasions. Regardless of site, we did not observe the production of mucosal IgA via skin colonization.

### Features of the immune response to intranasal colonization

The NALT consists of bilateral B cell rich lymphoid aggregates at the floor of the nasal cavity atop the mouse palate (**Fig. S12A**). The analogous anatomical feature in primates is Waldeyer’s ring, a set of lymphoid tissues (tonsils and adenoids) situated in the back of the nasal and oral cavities.^36,46^ The surface of the NALT is covered with a follicle-associated epithelium (FAE) with interspersed microfold (M) cells,^33,36,47^ mirroring Peyer’s patches in the intestines. These M cells, which express GP2 when mature, are thought to play a pivotal role in antigen sampling (**Fig. S12B**).^48–50^ To observe the fate of bacteria delivered intranasally, we labeled *S. epidermidis* with AlexaFluor488 and harvested the NALT for imaging at various time points after colonization. As early as 15 minutes after colonization, we observed bacteria on the surface of the FAE, co-localizing with UEA-1^+^ cells (**Fig. 3F**). Several hours after bacterial administration, we begin to see intact bacteria inside the NALT, persisting through our last time point (4 days post-colonization). Taken together, these observations are consistent with a model in which intact bacteria are transcytosed by M cells into the NALT, where they are processed in order to facilitate an antibody response to *S. epidermidis*.

### Mechanistic requirements for eliciting an antibody response on the skin and in the nose

Our data suggest that there are two mechanisms of antigen sampling: one on the skin that involves trans-epithelial sampling by an antigen-presenting cell, and the other in the nostril in which M cells in the NALT harvest antigen directly. Both represent promising channels for eliciting immunity, so we next sought to answer three questions about bacterial carrier of the antigen eliciting the response at each site: (*i*) Is there a requirement for bacteria at all? (*ii*) If so, do the bacteria need to be alive? (*iii*) Does the antigen need to be attached to the bacterium?

To address these questions, we designed an experiment in which we would compare four samples: live bacteria conjugated to TTFC, dead bacteria linked to TTFC, TTFC alone, and TTFC mixed with live bacteria. To make accurate comparisons among these immunogens, we needed to ensure that the same quantity of TTFC would be administered in each case. By comparing labeled bacteria to quantification beads bearing known quantities of antibody-binding sites, we estimate that our standard conjugation reaction yields ∼60,000 molecules of TTFC per *S. epidermidis* cell (**Fig. S12**). We reasoned that an antigen dose this high might obscure the comparisons we sought to make, so we used three quantities of antigen for each of the four conditions: 60,000 (high), 20,000 (medium), and 3,000 (low) molecules of TTFC per cell. For the protein-only and mixed conditions, we administered an equivalent quantity of antigen. Each of these 12 stimuli (four test articles x three doses) were administered separately to the skin or the nose, for a total of 24 conditions.

On the skin, live *S. epidermidis*-TTFC elicited a systemic IgG response at the highest dose, while none of the other test articles elicited any detectable response above background (**Figs. 3G, S13C**). These data suggest that during skin colonization, antigen needs to be physically attached to live bacteria in order to elicit a response.

In the nose, the clearest comparison was at the medium dose (**Figs. 3G, S13C**). There are three main findings: (*i*) live *S. epidermidis*-TTFC was more potent than dead *S. epidermidis*-TTFC, which was in turn more potent than TTFC alone. Thus, bacteria potentiate the response, live more so than dead. (*ii*) Live *S. epidermidis*-TTFC elicits a potent nasal and pulmonary IgA response, whereas the same quantity of pure TTFC elicits little to no IgA. This pattern, which holds at the high dose, is a striking example of a bifurcation in the IgA response that is dependent on live bacteria. (*iii*) To our surprise, live *S. epidermidis* mixed with TTFC was far more potent than TTFC alone and performs comparably to live *S. epidermidis* conjugated to TTFC. Thus, in contrast to the skin, antigen in the nostril does not need to be physically attached to a bacterium to elicit a strong response but still requires the presence of bacterial cells. This suggests an important difference in the mechanism of sampling on the skin and in the nostril, a finding we consider in more detail in the discussion.

### Reduced colonization schedule coupled with a non-classical adjuvant

Next, we turned from questions of mechanism to a series of experiments focused on the translational potential of an *S. epidermidis-*vectored vaccine. We wanted to test our vaccine candidates under more challenging conditions, where we reasoned that the addition of an adjuvant might augment the immune response. Classically, protein subunit and mRNA vaccines are almost universally adjuvanted, in contrast to live-attenuated and inactivated vaccines like the chickenpox (varicella) and polio vaccines, in which the pathogen serves as its own adjuvant.^51,52^ With this in mind, we tested the inclusion of all-trans retinoic acid (formulated as tretinoin), which is topically active, immune modulatory, and safe, but not considered a classical adjuvant.^53–56^

In most of the experiments up to this point, we administered eight or 13 doses over six weeks, a schedule that traces back to our earliest experiments characterizing the antibody response to *S. epidermidis*.^1^ With the addition of tretinoin, as few as two doses administered intranasally resulted in remarkably high and consistent systemic antibody titers, although four doses were required for robust induction of mucosal IgA (**Figs. 4A, S14A**). On the skin, while two doses were capable of inducing an immune response, more frequent dosing schedules resulted in titers that were similar to intranasal dosing, which we had never previously observed. Additionally, we found that frozen glycerol stocks of *S. epidermidis*-TTFC elicit comparable antibody titers to freshly labeled cultures (**Fig. S14B**), which is of practical importance for translational studies.

**Figure 4.**
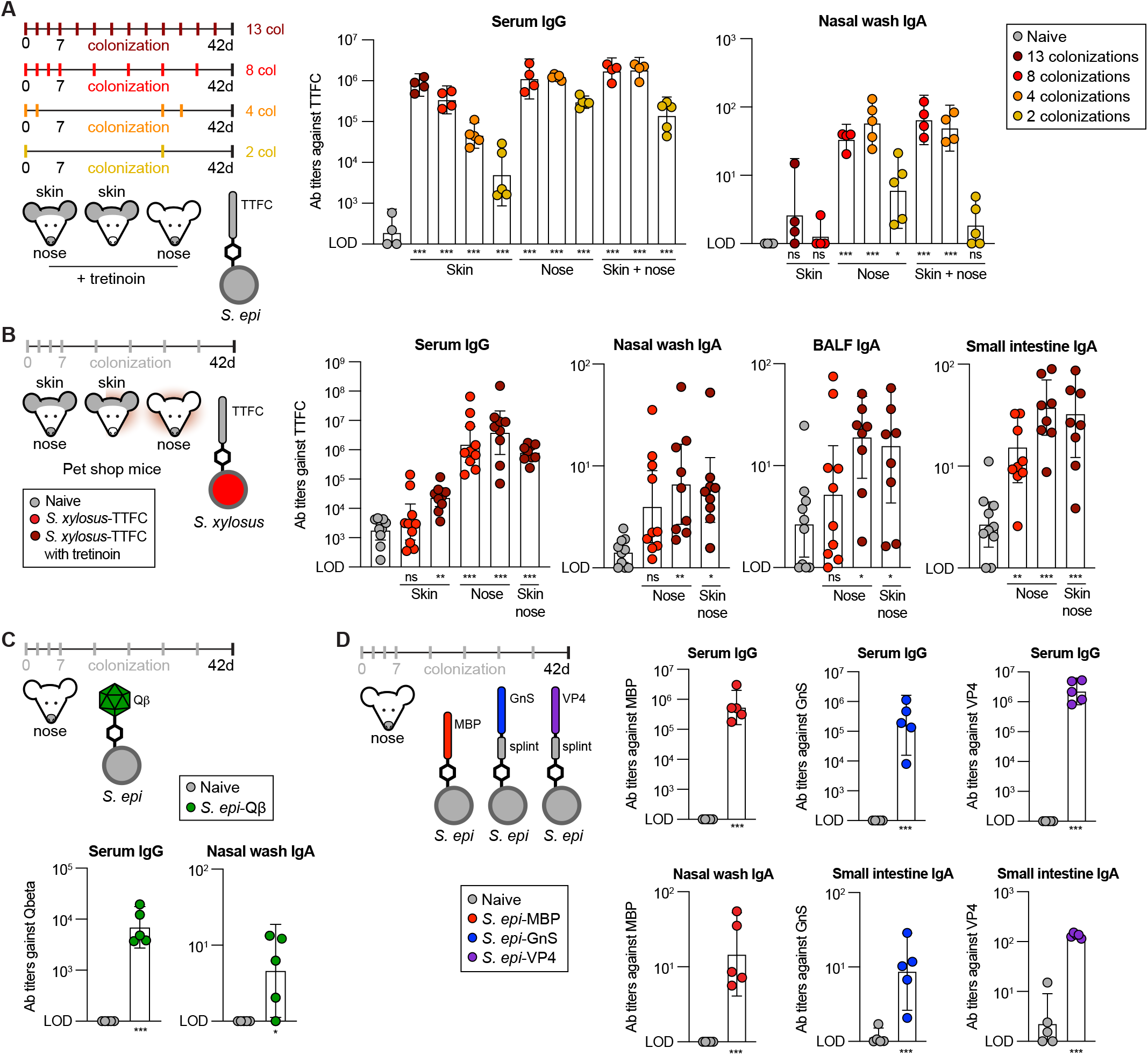
Commensal vaccines are active in pet shop mice and with a variety of antigens. (A) A reduced colonization schedule coupled with a non-classical adjuvant yields robust antibody responses topically and intranasally. Mice were colonized on the skin, in the nostril, or in both sites with *S. epidermidis*-TTFC and tretinoin. Dosing regimens of 2, 4, 8 or 13 colonizations were used. Nostril colonization was performed by pipetting 20 µL of *S. epidermidis-*TTFC (OD_600 =_ 6) into each nostril. After six weeks, serum and nasal wash were collected and analyzed by ELISA for antibody titers against TTFC. (B) Commensal vaccines elicit an antibody response in pet shop mice. Genetically mice of unknown ages from a local pet shop were colonized with *S. xylosus*-TTFC with and without tretinoin. Pet shop mice were colonized on the skin, in the nose (10 µL of *S. xylosus*-TTFC per nostril, OD_600 =_ 18), or at both sites eight times over six weeks. At the experimental endpoint, serum, nasal wash, BAL fluid, and small intestinal contents were harvested and assessed for TTFC-specific antibodies by ELISA. (C) Bacteriophage Qβ is immunogenic when conjugated to *S. epidermidis.* Pure Qβ capsids were derivatized with NHS-DBCO and conjugated to the surface of *S. epidermidis* that had been functionalized with NHS-azide. *S. epidermidis*-Qβ was used to colonize mice intranasally eight times over six weeks (10 µL OD_600 =_ 18 per nostril). Serum and nasal wash were collected, and antibody titers against Qβ were measured by ELISA. (D) Commensal vaccination with a variety of additional antigens. *S. epidermidis* was conjugated to the model antigen maltose binding protein (MBP) from *E. coli* using NHS-biotin and a streptavidin-MBP fusion protein. The clinically relevant antigens VP4 (from rotavirus) and glycoprotein GnS (from Rift Valley fever virus, RVFV) were purified as SpyTag003 fusions and attached to biotinylated *S. epidermidis* using a streptavidin-SpyCatcher003 fusion splint protein. These strains were used to colonize mice intranasally (10 µL of OD_600 =_ 18 per nostril) eight times over six weeks. Serum and mucosal tissues were assayed for antibodies specific for each antigen by ELISA; endpoint titers are shown. Data shown are from one (**A, D**) or representative of two (**B, C**) independent experiments. Graphs show geometric mean with a 95% confidence interval. *P* values (ns, not significant; * *p* < 0.05; ** *p* < 0.01; *** *p* < 0.001) were calculated using Welch’s t tests (**C**, anti-MBP nasal wash IgA in panel **D**), ordinary one-way ANOVA followed by Tukey’s multiple comparisons test (**A, B**, anti-MBP serum IgG in panel **D**), assuming log-normal distributions, or two-way ANOVA on log-transformed data followed by pairwise testing with Šidák’s correction for multiple comparisons (**D**). Asterisks in figures represent pairwise comparisons of the indicated group to naive mice; *p* values for all comparisons are reported in **Table S1**.

### Commensal vaccination works in pet shop mice

SPF C57BL/6 mice have two characteristics that might make them hypersensitive to commensal vaccination: their skin is not as densely colonized as mice living under real-world conditions, and they are less immunologically experienced, which might make them more sensitive to new microbial exposures.^57,58^ Pet shop and wild-caught mice, on the other hand, have more densely colonized skin and an immune system that better mimics that of an adult human.^57,59,60^ This is important translationally: an *S. epidermidis-*vectored vaccine would have to elicit a potent immune response in the presence of a thoroughly colonized skin or nasal mucosal surface. Furthermore, pet shop mice are genetically outbred, better recapitulating the diversity in immune responses found in the human population.

To determine whether vaccination with a topical colonist elicits an immune response in pet shop mice, we conjugated TTFC to two skin commensals that elicited strong antibody titers in our previous experiments—the human colonist *S. epidermidis* and the mouse isolate *S. xylosus*—and colonized mice from a local pet shop on the skin, in the nose, or in both sites.

Although pet shop mice had higher background titers and more variable immune responses, colonization with *S. epidermidis-*TTFC and *S. xylosus*-TTFC elicited a robust antibody response; the murine isolate was more potent than the human isolate (**Figs. 4B, S14C**). The immune response in pet shop mice mirrored our findings from SPF C57BL/6 mice: antibody titers were higher from intranasal colonization than skin colonization, including IgA at mucosal surfaces (**Fig. 4B**). Induction of immunity through the skin required the addition of tretinoin, which also augmented the response to intranasal administration. More broadly, the robust responses elicited by *Staphylococcus* colonization in pet shop mice bode well for the vaccination of large animals in real-world settings.

### Immunogenicity of additional antigens

In principle, chemical conjugation should be capable of linking a wide variety of antigens to the cell surface, including immunogens that cannot be folded, assembled, or post-translationally modified properly in *Staphylococcus.* However, most of our studies have used TTFC as a model antigen. We carried out two experiments to determine whether more complex antigens are immunogenic when displayed on the bacterial cell surface.

First, in light of our observation that the protein-based nanoparticle mi3-GFP can be conjugated efficiently to the surface of *S. epidermidis* (**Fig. 1E**, ∼40 nm), we attempted to attach a whole viral capsid to the surface of *S. epidermidis*. We selected the bacteriophage Qβ, which self-assembles into a ∼28 nm capsid that is amenable to surface functionalization due to a large number of surface-exposed primary amines.^61–63^ We conjugated DBCO to Qβ using amine-reactive crosslinkers and incubated this derivatized capsid with azide-linked *S. epidermidis*, resulting in Qβ capsids bound to the surface of *S. epidermidis* (**Figs. S15C, S15D**). By colonizing mice intranasally with *S. epidermidis-*Qβ, we elicited antibodies against Qβ both systemically and at mucosal surfaces (**Figs. 4C, S15E**). This suggests that commensal vaccination is a viable strategy for immunization using whole viral capsids as antigens.

Second, we conjugated a series of recombinant immunogens including the immunogenic maltose-binding protein (MBP, *E. coli*),^64^ the carrier protein exoprotein A (EPA, *Pseudomonas aeruginosa*),^65,66^ and stabilized forms of the glycoprotein Gn from Rift Valley fever virus and the VP4 spike of human rotavirus (GnS and VP4, to be described elsewhere). GnS and VP4 were purified as SpyTag003 fusion proteins and coupled to the bacterial surface using a streptavidin-SpyCatcher003 splint protein (**Fig. S15F**).^67^ We colonized mice with all four of these strains intranasally (*S. epidermidis*-MBP, *S. epidermidis*-EPA, *S. epidermidis*-GnS, *S. epidermidis*-VP4); in each case, we elicited a robust systemic and mucosal antibody response (**Figs. 4D, S15G**). Particularly striking was the intestinal IgA induced upon colonization with *S. epi*-VP4, which may protect against rotavirus infection.

## DISCUSSION

Chemical conjugation is simple, inexpensive, and easy to use. We anticipate that it will be enabling in four ways: (*i*) it opens the door to studying bacterial species from the microbiome that are difficult to manipulate genetically; (*ii*) it offers precise control over antigen quantity, enabling studies in the physiologic (moderate density) vs. therapeutic (high-density) regimes and comparisons between different species using the same quantity of antigen; (*iii*) it facilitates the attachment of payloads that cannot be attached via genetic engineering or are inefficiently expressed; and (*iv*) protocols are straightforward and will be accessible to non-specialized labs. We have not exhaustively characterized the chemistries that could be used to attach payloads to the cell surface; other surface chemistries will likely work.^68^

Bacterial species could elicit an immune response independent of their ecological origin, colonization efficiency, or native immune modulatory properties. We were surprised to see the potent inductive properties of Gram-negative organisms, given that the predominant colonists of the skin and nostril are Gram-positive.^28,29^ Since the response to *S. epidermidis* is dominated by induction in the NALT, we suspect that the response to environmental strains reflects sampling in the nostril and may not appear when the same strains are applied to the skin alone.

The potency of the immune response in the nostril was surprising. We had assumed we were studying the response to skin colonization; we had not realized that the comparably small volume of liquid getting into the nares had a predominant effect. While our data suggest the NALT as the primary inductive site, we cannot rule out other lymphoid tissues in the airways, either constitutive or induced.^69–73^

Nostril colonization is physiologic; *S. epidermidis* is a universal colonist of the nostril, and the role of this site in immune induction remains largely unexplored.^33–36^ Intranasal administration of *S. epidermidis* with antigen resulted in extremely potent antibody responses, both systemically and along the mucosal chain (the nasal passage, the lungs and the intestine). In contrast, traditional intramuscular vaccination is largely unable to elicit a mucosal response,^74,75^ while intranasal vaccination efforts (e.g. FluMist) have faced challenges eliciting durable systemic immunity.^76–79^ Live *S. epidermidis* might be a ‘natural ligand’ for the immune sampling process in the nostril. It presents a compelling opportunity to explore the mechanisms of sampling and the induction of mucosal immunity from intranasal administration.

Live bacteria, either conjugated to or mixed with antigen, gave a potent systemic and mucosal response via intranasal administration. The immune response to the ‘mixed’ condition (*S. epidermidis* + antigen) was unexpected and could be explained by several mechanistic possibilities. For example, intranasal administration of bacteria may activate a sampling process in the NALT that acts broadly and enables fluid-phase protein sampling. Alternatively, the presence of live bacteria in the NALT may promote a downstream immune response to soluble antigen that is constitutively sampled. In contrast, the requirements on the skin are more stringent; antigen must be physically linked to live bacteria, highlighting that the sampling system can discriminate between live bacteria and other forms of antigen. Defining the sampling processes at each of these barrier sites will ultimately enable rational design of vaccines tailored to each route.

Conjugating a dye to the cell surface allowed us to use bacteria as a staining reagent for antigen-specific B cells *in vivo*. This simple and powerful approach has several important use cases. For example, it could be used to map the targets of commensal-specific B cells, with the goal of discovering patterns that underlie antigenicity. This technique will also enable analysis of the largely uncharacterized human antibody repertoire, allowing us to answer a variety of questions regarding the immune response to commensals and how it can be harnessed for vaccination.^80–82^

Chemical conjugation of antigens to the bacterial cell surface yields a robust systemic and mucosal antibody response to skin and nostril colonization. Commensal vaccines are effective in pet shop mice and can be made with a variety of clinically relevant antigens. The data presented here outline a compelling translational path for a class of vaccines that would be needle-free, self-applied, and straightforward to distribute.^83–85^

## ACKNOWLEDGMENTS

We are deeply indebted to members of the Fischbach laboratory, especially M. McLaughlin for chemical biology discussions; J. Bunker for experimental questions; A. Espinoza and J. Au for laboratory management and administrative support; D. Wattendorf, C. Karp, O. Vandal, C. Somerville, and H. Youngs for ideas and critical feedback; J. Rajagopal and R. Chivukula for helpful discussions; A. Jacobson and M. Levia for coordination with the Stanford Microbiome Therapies Initiative; the Stanford University Veterinary Service Center for animal husbandry; the Stanford Shared FACS Facility (RRID: SCR_017799); the Stanford Cell Sciences Imaging Facility; and Zeiss LSM880 confocal (RRID SCR_017787) at the Beckman Center for confocal microscopy imaging. This work was supported by the Gates Foundation (M.A.F.), Coefficient Giving (M.A.F.), the HS Chau Foundation (M.A.F.), the Stanford Microbiome Therapies Initiative, the Chan Zuckerberg Biohub (M.A.F.), NIH grants R01 AI175642 (M.A.F.), K99 AI180358 (D.B.), the Helen Hay Whitney Foundation (K.D.B.), the Knight-Hennessy Fellowship (P.V.L.), a Department of Defense NDSEG Fellowship (P.V.L.), the Hertz Foundation Fellowship (D.B.L.), the Gates Foundation INV-043758 (N.P.K.), and the National Institute of Allergy and Infectious Diseases U19AI181881 (N.P.K.).

## AUTHOR CONTRIBUTIONS

K.D.B., P.V.L., D.B. and M.A.F. conceived of the study. Most experiments were done by K.D.B., P.V.L., A.J., M.C., and T.N.R., with help from D.B. J.M.S., E.T., D.B.L., A.N., A.T., M.L., A.Z., and M.T.. K.D.B. and P.V.L. developed the methodology and analyzed the data. H.K., F.R., N.J., Z.J., and N.P.K. designed, expressed, and purified rotavirus and RVFV immunogens. The manuscript was drafted by K.D.B., P.V.L., and M.A.F. All authors reviewed and approved the manuscript.

