## Supplement for "Microbial mechanistic requirements for eliciting a topical and intranasal immune response"

### TABLE OF CONTENTS


**Fig. S1.** Optimization of biotin/streptavidin-mediated surface display on *S. epidermidis.* ........ 3-4

**Fig. S2.** Optimization of azide/alkyne (DBCO) chemical conjugation to *S. epidermidis* and attachment of novel payloads to the bacterial surface. ........ 5-6

**Fig. S3.** Chemical conjugation of antigens to the surface of *S. epidermidis* can elicit immune responses *in vivo*. ........ 7-8

**Fig. S4**. *Staphylococcus* strains from diverse sources can be labeled using NHS-ester chemistry ........ 9-10

**Fig. S5.** Bacterial strains with varying native immunogenicity can be conjugated to TTFC and induce antibody responses in mice ........ 11-12

**Fig. S6.** Bacterial strains isolated from a variety of environments can be labeled with TTFC and used to induce antibody responses in mice ........ 13-14

**Fig. S7.** Heat-killed *S. epidermidis* can be efficiently surface labeled with proteins ........ 15-16

**Fig. S8.** Nasal colonization induces an IgA response at mucosal sites. ........ 17-18

**Fig. S9.** Fluorescent *S. epidermidis* can be used as a staining reagent to identify bacteria-specific B cells ........ 19-21

**Fig. S10.** Low-volume intranasal delivery elicits immune responses without draining to the lungs ........ 22-23

**Fig. S11.** Several skin sites are competent to elicit immunity after colonization ........ 24-25

**Fig. S12.** Whole bacteria are present in the NALT after nasal colonization. ........ 26-28

**Fig. S13.** Mechanistic requirements for eliciting antibody responses after topical or intranasal colonization ........ 29-30

**Fig. S14.** Compressed dosing regimens and activity in pet shop mice ........ 31-32

**Fig. S15.** Commensal vaccination with clinically relevant antigens ........ 33-34

**METHODS** ........ 36-46


**OTHER SUPPLEMENTARY MATERIALS**


**Table S1**. All statistical testing and p values

**Table S2**. Bacterial strains and growth conditions

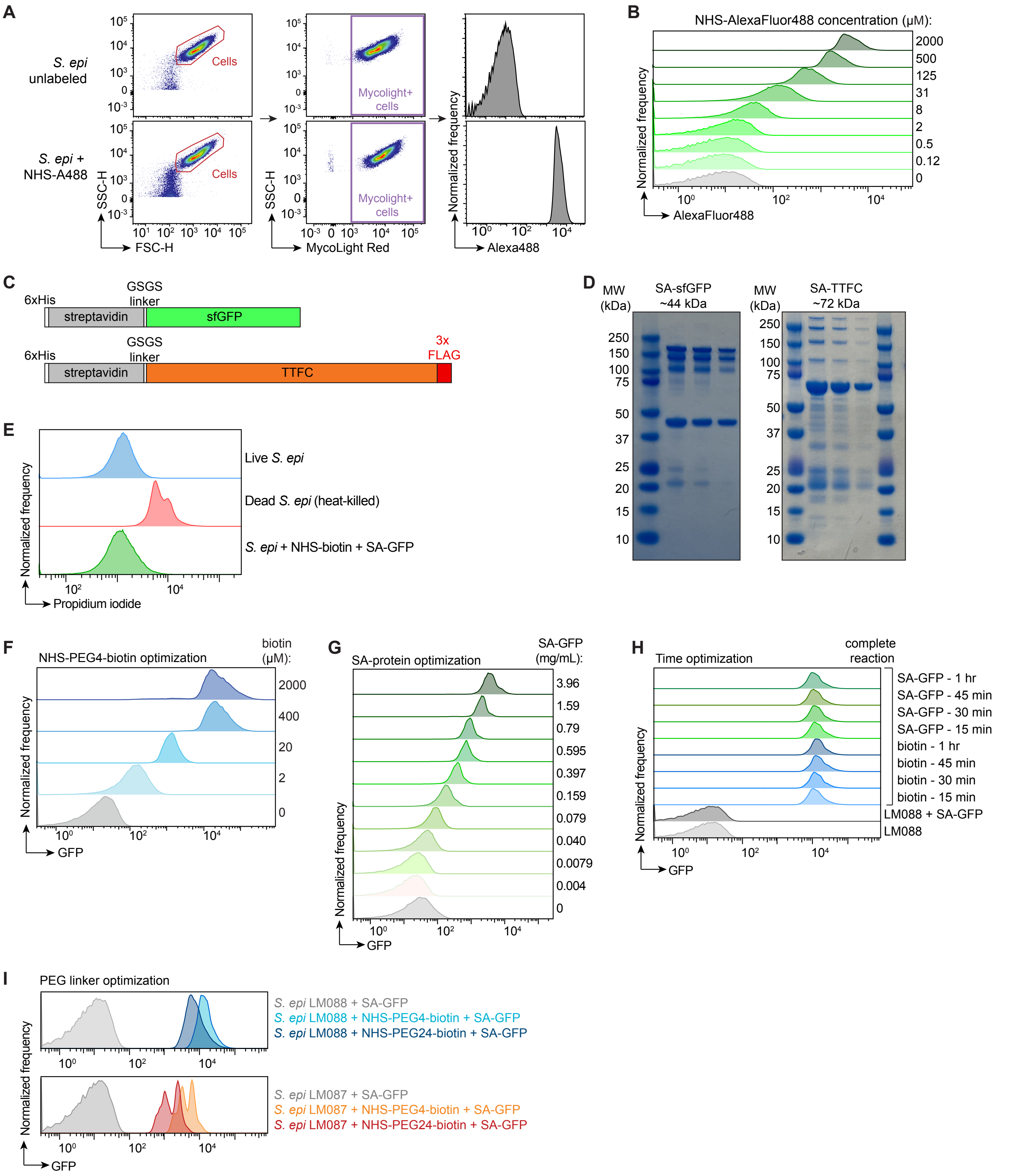

A
S. epi unlabeled
S. epi + NHS-A488
Cells
Mycolight+ cells
SSC-H
FSC-H
MycoLight Red
Normalized frequency
Alexa488
B
NHS-AlexaFluor488 concentration (μM):
2000
500
125
31
8
2
0.5
0.12
0
AlexaFluor488
C
6xHis
GSGS linker
streptavidin
sfGFP
TTFC
3x FLAG
D
MW (kDa)
SA-sfGFP ~44 kDa
SA-TTFC ~72 kDa
E
Live S. epi
Dead S. epi (heat-killed)
S. epi + NHS-biotin + SA-GFP
Propidium iodide
F
NHS-PEG4-biotin optimization
biotin (μM):
2000
400
20
2
0
GFP
G
SA-protein optimization
SA-GFP (mg/mL):
3.96
1.59
0.79
0.595
0.397
0.159
0.079
0.040
0.0079
0.004
0
H
Time optimization
complete reaction
SA-GFP - 1 hr
SA-GFP - 45 min
SA-GFP - 30 min
SA-GFP - 15 min
biotin - 1 hr
biotin - 45 min
biotin - 30 min
biotin - 15 min
LM088 + SA-GFP
LM088
I
PEG linker optimization
S. epi LM088 + SA-GFP
S. epi LM088 + NHS-PEG4-biotin + SA-GFP
S. epi LM088 + NHS-PEG24-biotin + SA-GFP
S. epi LM087 + SA-GFP
S. epi LM087 + NHS-PEG4-biotin + SA-GFP
S. epi LM087 + NHS-PEG24-biotin + SA-GFP

**Figure S1**. Optimization of biotin/streptavidin-mediated surface display on *S. epidermidis.*

(**A**) Gating strategy for bacterial flow cytometry to assess surface conjugation. Bacteria are distinguished by MycoLight Red$^+$ gate and labeling efficiency is determined by Alexa488 fluorescence intensity.

(**B**) Flow cytometry histograms of *S. epidermidis* (MycoLight$^+$) incubated with varying concentrations of NHS-ester AlexaFluor488 to determine labeling efficiency.

(**C**) Construct design for streptavidin fusion proteins (GFP and TTFC) used for bacterial surface attachment with NHS-PEG4-biotin. Based on Addgene_124296.

(**D**) SDS-PAGE gel (4-12% BisTris) with Coomassie staining of purified streptavidin-GFP (monomer ~44 kDa) and streptavidin-TTFC (monomer ~72 kDa) fusion proteins. Three concentrations shown per protein on gel. Higher molecular weight bands due to streptavidin multimerization.

(**E**) Flow cytometry plots of live *S. epidermidis* (blue), heat-killed *S. epidermidis* (red), and *S. epidermidis* conjugated to SA-GFP using NHS-ester chemistry (green), showing that *S. epidermidis* (MycoLight$^+$) is alive after labeling as indicated by propidium iodide staining.

(**F**) Flow cytometry analysis of NHS-PEG4-biotin concentration optimization for bacterial surface attachment. *S. epidermidis* was incubated with different concentrations of NHS-PEG4-biotin followed by SA-GFP (1 mg/mL) and visualized by flow cytometry (MycoLight$^+$). Each histogram represents a different concentration of NHS-PEG4-biotin used in the reaction.

(**G**) Protein conjugation to the *S. epidermidis* surface can be titrated by varying the amount of protein in the reaction. *S. epidermidis* was incubated with NHS-PEG4-biotin (2 mM) followed by varying concentrations of SA-GFP, and labeling efficiency was assessed by flow cytometry (MycoLight$^+$ cells). Each histogram represents a different concentration of SA-GFP used in the reaction.

(**H**) Optimization of reaction times for NHS-PEG4-biotin and SA-GFP reactions on the bacterial cell surface. To determine the optimal timing for the NHS-ester reaction, *S. epidermidis* was incubated with NHS-PEG4-biotin (2 mM) for varying lengths of time followed by SA-GFP (1 mg/mL) for 30 min (blue histograms). To determine optimal timing for the biotin/streptavidin reaction, S. epidermidis was incubated with NHS-PEG4-biotin (2 mM) for 30 min followed by incubation with SA-GFP (1 mg/mL) for varying lengths of time. Labeling efficiency determined by flow cytometry (MycoLight$^+$ cells).

(**I**) Polyethylene glycol (PEG) linker length impacts bacterial conjugation. *S. epidermidis* LM088 (top plot) and *S. epidermidis* LM087 (bottom) were incubated with either NHS-PEG4-biotin or NHS-PEG24-biotin, followed by SA-GFP. No biotin reaction included as a control. Surface labeling was visualized by flow cytometry (MycoLight$^+$ cells).

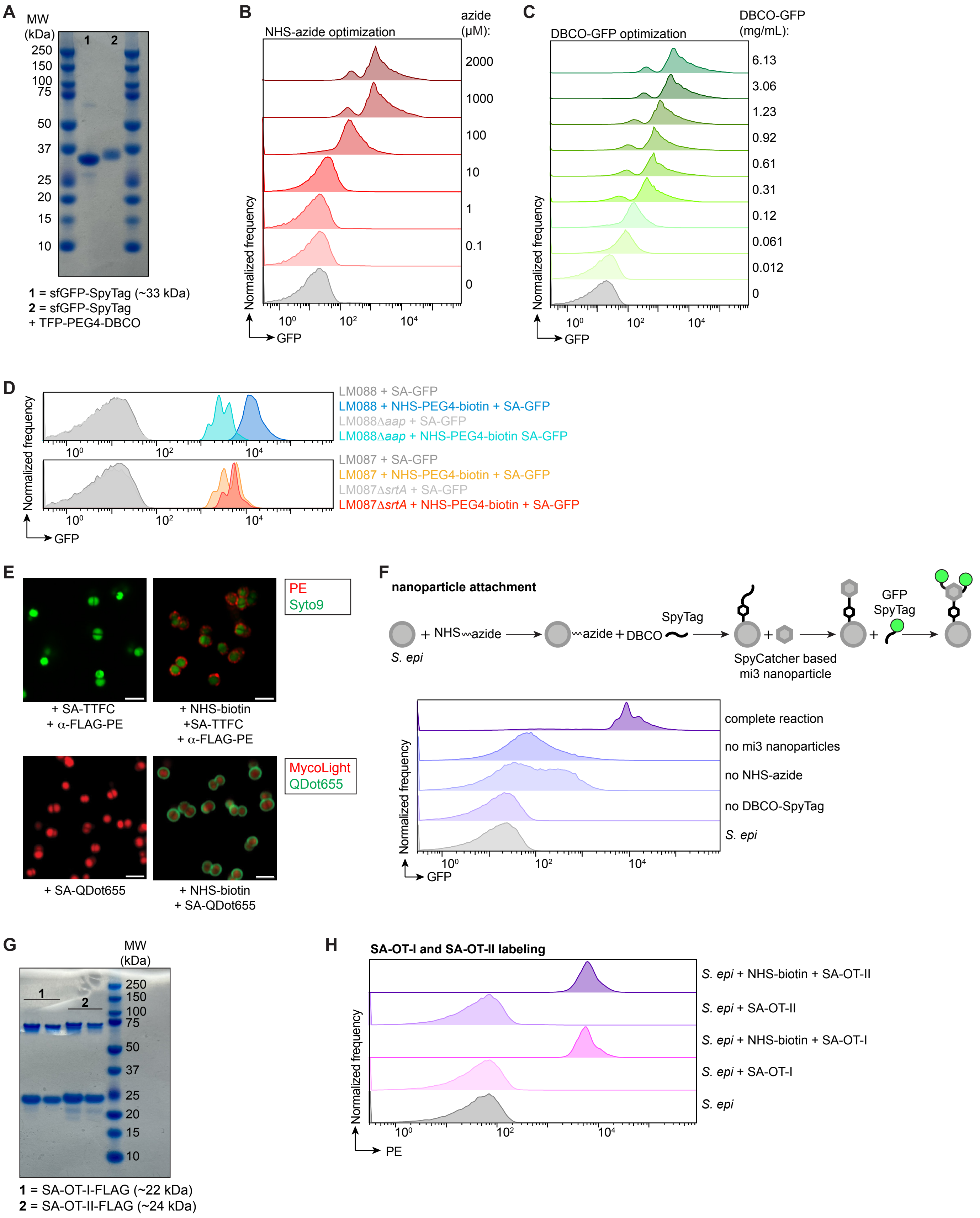
A
MW (kDa)
1 2
250
150
100
75
50
37
25
20
15
10
1 = sfGFP-SpyTag (~33 kDa)
2 = sfGFP-SpyTag + TFP-PEG4-DBCO
B
NHS-azide optimization
azide (µM):
2000
1000
100
10
1
0.1
0
Normalized frequency
GFP
C
DBCO-GFP optimization
DBCO-GFP (mg/mL):
6.13
3.06
1.23
0.92
0.61
0.31
0.12
0.061
0.012
0
Normalized frequency
GFP
D
LM088 + SA-GFP
LM088 + NHS-PEG4-biotin + SA-GFP
LM088Δaap + SA-GFP
LM088Δaap + NHS-PEG4-biotin SA-GFP
LM087 + SA-GFP
LM087 + NHS-PEG4-biotin + SA-GFP
LM087ΔsrtA + SA-GFP
LM087ΔsrtA + NHS-PEG4-biotin + SA-GFP
Normalized frequency
GFP
E
PE
Syto9
+ SA-TTFC + α-FLAG-PE
+ NHS-biotin +SA-TTFC + α-FLAG-PE
MycoLight
QDot655
+ SA-QDot655
+ NHS-biotin + SA-QDot655
F
nanoparticle attachment
S. epi
+ NHS azide
azide + DBCO
SpyTag
SpyCatcher based mi3 nanoparticle
GFP SpyTag
complete reaction
no mi3 nanoparticles
no NHS-azide
no DBCO-SpyTag
S. epi
Normalized frequency
GFP
G
MW (kDa)
1
2
250
150
100
75
50
37
25
20
15
10
1 = SA-OT-I-FLAG (~22 kDa)
2 = SA-OT-II-FLAG (~24 kDa)
H
SA-OT-I and SA-OT-II labeling
S. epi + NHS-biotin + SA-OT-II
S. epi + SA-OT-II
S. epi + NHS-biotin + SA-OT-I
S. epi + SA-OT-I
S. epi
Normalized frequency
PE

**Figure S2**. Optimization of azide/alkyne (DBCO) chemical conjugation to *S. epidermidis* and attachment of novel payloads to the bacterial surface.

(**A**) SDS-PAGE gel (4-12% BisTris) and Coomassie staining of GFP-SpyTag before and after derivatization with TFP-PEG4-DBCO. Lane 1 = GFP-SpyTag (~33 kDa), lane 2 = GFP-SpyTag incubated with TFP-PEG4-DBCO and purified by size exclusion spin column (increase in size observed).

(**B**) Optimization of NHS-PEG4-azide concentrations for surface labeling of *S. epidermidis*. Bacteria were incubated with varying concentrations of NHS-PEG4-azide (indicated on plot) followed by DBCO-GFP (1 mg/mL) and conjugation efficiency was assessed by flow cytometry. Histograms show MycoLight$^+$ cells.

(**C**) Flow cytometry histograms showing *S. epidermidis* (MycoLight$^+$) labeling with varying concentrations of DBCO-GFP. *S. epidermidis* was incubated with NHS-PEG4-azide (1 mM) followed by different concentrations of DBCO-GFP (GFP derivatized immediately prior to use).

(**D**) PEG linker length optimization for azide/alkyne click reactions on the bacterial cell surface. *S. epidermidis* was incubated with TFP-PEG$_n$-azide (n = 4, 12, or 24) followed by DBCO-Cy5. No azide reaction included as a control. Conjugation was assessed by flow cytometry; histograms show GFP fluorescence intensity of MycoLight$^+$ cells.

(**E**) Confocal microscopy images of *S. epidermidis* conjugated to SA-TTFC or quantum dots (Qdot655) using NHS-PEG4-biotin. No biotin samples included as controls. Cells visualized by Syto9 and a-FLAG-PE staining (TTFC only). Scale bar = 2 µm.

(**F**) Schematic demonstrating strategy for the chemical attachment of protein-based mi3 nanoparticles to the bacterial cell surface. Bacteria were incubated with NHS-PEG4-azide followed by DBCO-SpyTag003. SpyCatcher003-based mi3 nanoparticles were then added to the bacteria for a short incubation followed by an excess of GFP-SpyTag to bind the nanoparticle structure. Attachment to the bacterial surface was assessed by flow cytometry; histograms represent GFP fluorescence of MycoLight$^+$ cells, and each histogram represents a reaction where one component was omitted as a control, until the top histogram (labeled complete reaction).

(**G**) SDS-PAGE gel (4-12% BisTris) with Coomassie staining of SA-OT-I (1, ~22 kDa) and SA-OT-II (2, ~24 kDa). Two concentrations of protein shown on gel. Higher molecular weight band indicative of streptavidin multimerization.

(**H**) Flow cytometry analysis of *S. epidermidis* labeled with SA-OT-I or SA-OT-II. Bacteria were incubated with NHS-PEG4-biotin (2 mM) followed by SA-OT-I (4.5 mg/mL) or SA-OT-II (5 mg/mL). Protein attachment was determined by staining with a-FLAG-PE and flow cytometry analysis. No biotin reactions were included as controls. Each histogram represents PE fluorescence of Syto9$^+$ cells.

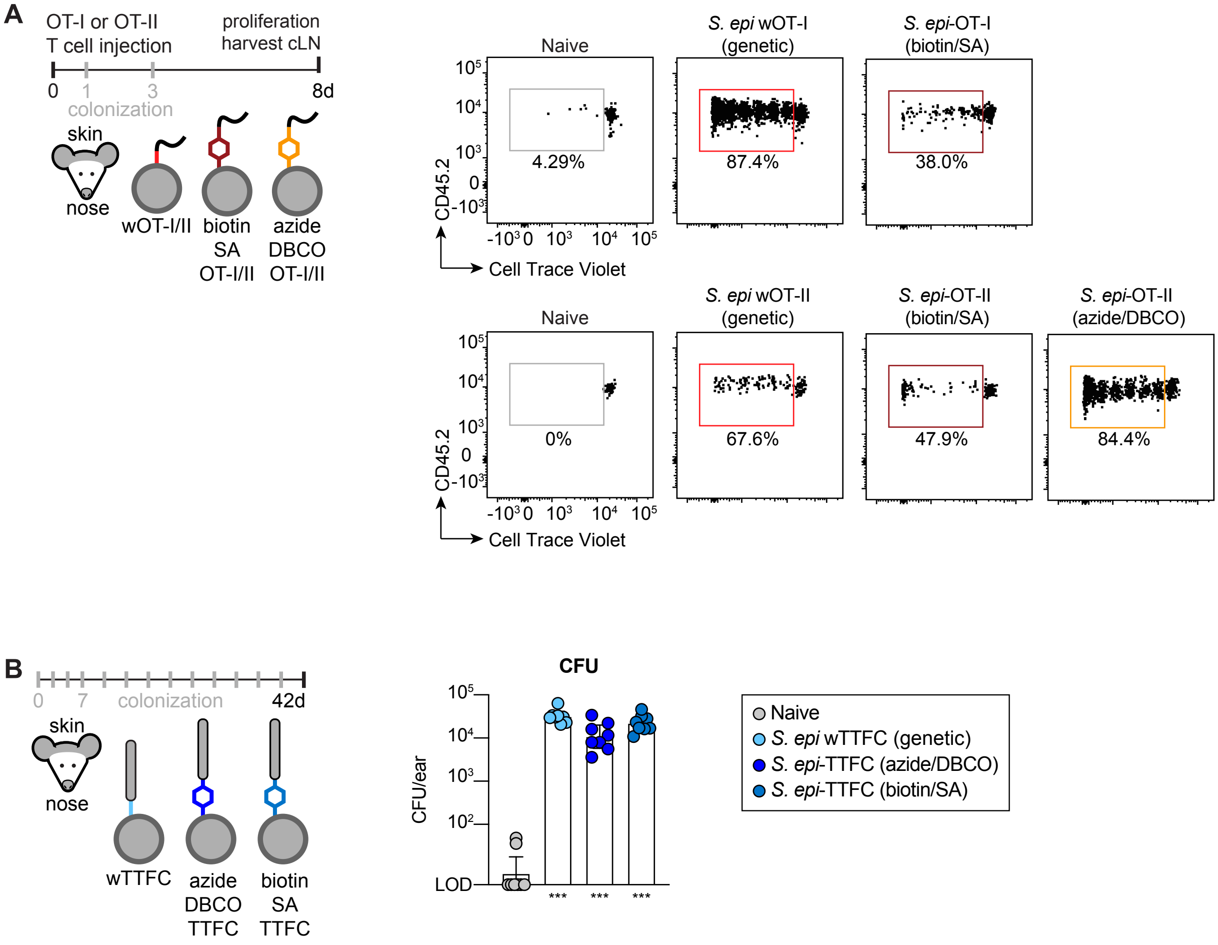
A
OT-I or OT-II
T cell injection
proliferation
harvest cLN
0
1
3
8d
colonization
skin
nose
wOT-I/II
biotin
SA
OT-I/II
azide
DBCO
OT-I/II
Naive
S. epi wOT-I
(genetic)
S. epi-OT-I
(biotin/SA)
4.29%
87.4%
38.0%
CD45.2
Cell Trace Violet
Naive
S. epi wOT-II
(genetic)
S. epi-OT-II
(biotin/SA)
S. epi-OT-II
(azide/DBCO)
0%
67.6%
47.9%
84.4%
CD45.2
Cell Trace Violet
B
0
7
colonization
42d
skin
nose
wTTFC
azide
DBCO
TTFC
biotin
SA
TTFC
CFU
CFU/ear
LOD
***
***
***
Naive
S. epi wTTFC (genetic)
S. epi-TTFC (azide/DBCO)
S. epi-TTFC (biotin/SA)

**Figure S3.** Chemical conjugation of antigens to the surface of *S. epidermidis* can elicit immune responses *in vivo*.

(**A**) Representative flow plots of *in vivo* proliferation (assessed by dilution of CellTrace Violet) of adoptively transferred OT-I and OT-II T cells after colonization of the skin and nose of mice with *S. epidermidis* wOT-I or wOT-II (genetic), *S. epidermidis*-OT-I or -OT-II (biotin/SA attachment), or *S. epidermidis*-OT-I or -OT-II (azide/alkyne click conjugation). Plots represent Live $Tcrb^+$ $CD8^+$ $CD4^-$ $CD45.2^+$ $CD45^-$ $Vb5^+$ $Va2^+$ cells for OT-I cells or Live $Tcrb^+$ $CD4^+$ $CD8^-$ $CD45.2^+$ $CD45^-$ $Vb5^+$ $Va2^+$ for OT-II cells.

(**B**) Data shown here are from the same experiment in **Fig. 1G**. Mice were colonized with *S. epidermidis* LM088 genetically expressing or chemically conjugated to TTFC, either by azide/alkyne cycloaddition or biotin/streptavidin interactions. Mice were colonized 13 times over six weeks, after which ears were harvested, homogenized, and serial dilutions were plated in triplicate. Data are expressed as CFU per ear, and baseline indicates limit of detection (LOD) (4 CFU/ear). Data shown are pooled from two independent experiments.

Data shown are from one (**A**) or two experiments (**B**). *P* values (ns, not significant; * $p < 0.05$; ** $p < 0.01$; *** $p < 0.001$) were calculated using an ordinary one-way ANOVA followed by Tukey's multiple comparisons test, assuming or log-normal distributions. Asterisks in figures represent pairwise comparisons of the indicated group to naive mice; *p* values for all comparisons are reported in **Table S1**.

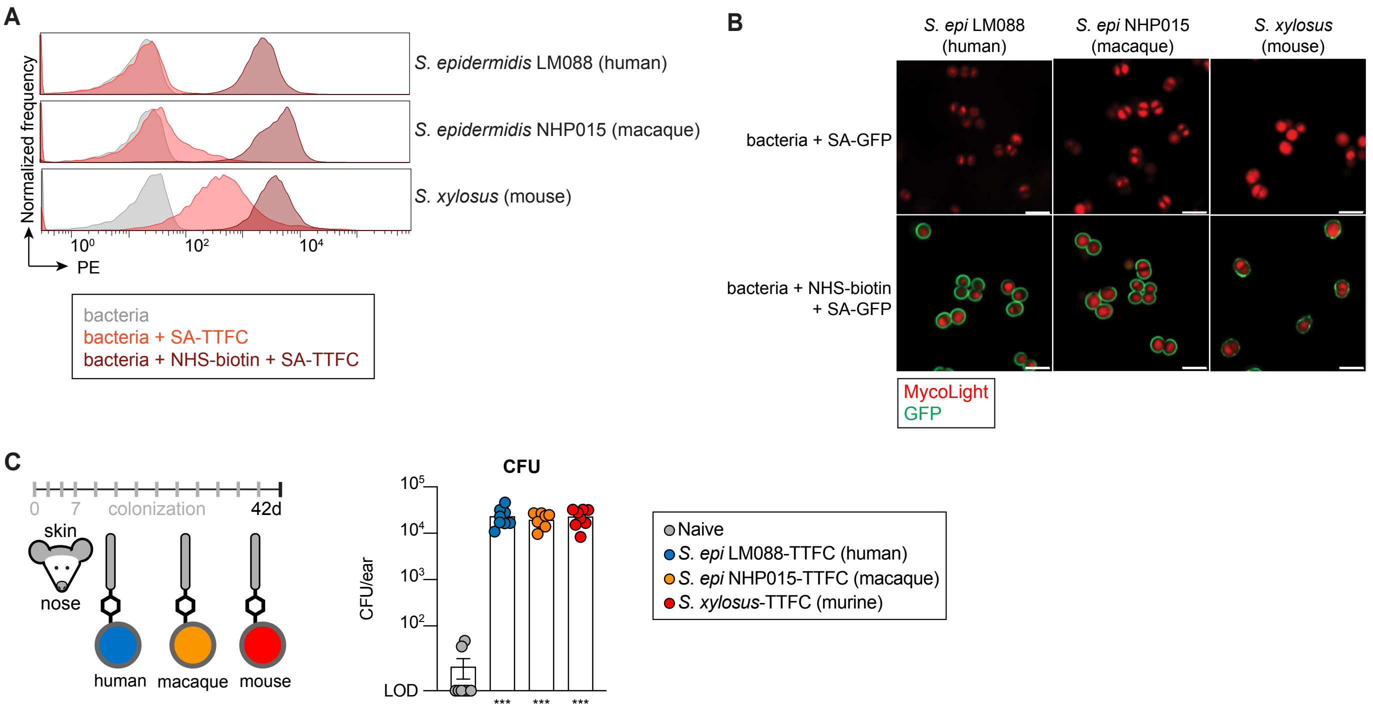
A
Normalized frequency
S. epidermidis LM088 (human)
S. epidermidis NHP015 (macaque)
S. xylosus (mouse)
10⁰
10²
10⁴
PE
bacteria
bacteria + SA-TTFC
bacteria + NHS-biotin + SA-TTFC
B
S. epi LM088 (human)
S. epi NHP015 (macaque)
S. xylosus (mouse)
bacteria + SA-GFP
bacteria + NHS-biotin + SA-GFP
MycoLight
GFP
C
0
7
colonization
42d
skin
nose
human
macaque
mouse
CFU
10⁵
10⁴
10³
10²
LOD
CFU/ear
***
***
***
Naive
S. epi LM088-TTFC (human)
S. epi NHP015-TTFC (macaque)
S. xylosus-TTFC (murine)

**Figure S4**. *Staphylococcus* strains from diverse sources can be labeled using NHS-ester chemistry.

(**A**) Flow cytometry histograms showing *S. epidermidis* LM088 (human), *S. epidermidis* NHP015 (macaque), and *S. xylosus* (mouse) can be labeled with the model immunogen TTFC using NHS-PEG4-biotin. Labeled bacteria were stained with anti-FLAG-PE to detect attachment of TTFC-FLAG, and histograms show PE fluorescence of Syto9$^{+}$ cells for each strain. Gray histograms show bacteria + anti-FLAG-PE, light red histograms indicate no biotin control reactions, and dark red histograms are complete reactions.

(**B**) Confocal microscopy of *S. epidermidis* LM088, *S. epidermidis* NHP015, and *S. xylosus* surface labeled with NHS-biotin and SA-GFP. Cells identified as MycoLight$^{+}$. No biotin reaction included as a control. Scale bar = 2 μm.

(**C**) Data shown are from the same experiment as **Fig. 2B**. Mice were colonized on the skin and nose with different *Staphylococcus* strains from human, macaque, or mouse hosts labeled with TTFC. After 13 colonizations in six weeks, ears were harvested, and bacterial burden was assessed by plating for CFUs. Data are shown as CFU per ear, and represent two pooled independent experiments. Baseline indicates limit of detection (LOD = 4 CFU/ear).

Data shown are from at least two independent experiments. Graphs show geometric mean with a 95% confidence interval. *P* values (ns, not significant; * $p < 0.05$; ** $p < 0.01$; *** $p < 0.001$) were calculated using an ordinary one-way ANOVA followed by Tukey's multiple comparisons test, assuming log-normal distributions. Asterisks in figures represent pairwise comparisons of the indicated group to naive mice; *p* values for all comparisons are reported in **Table S1**.

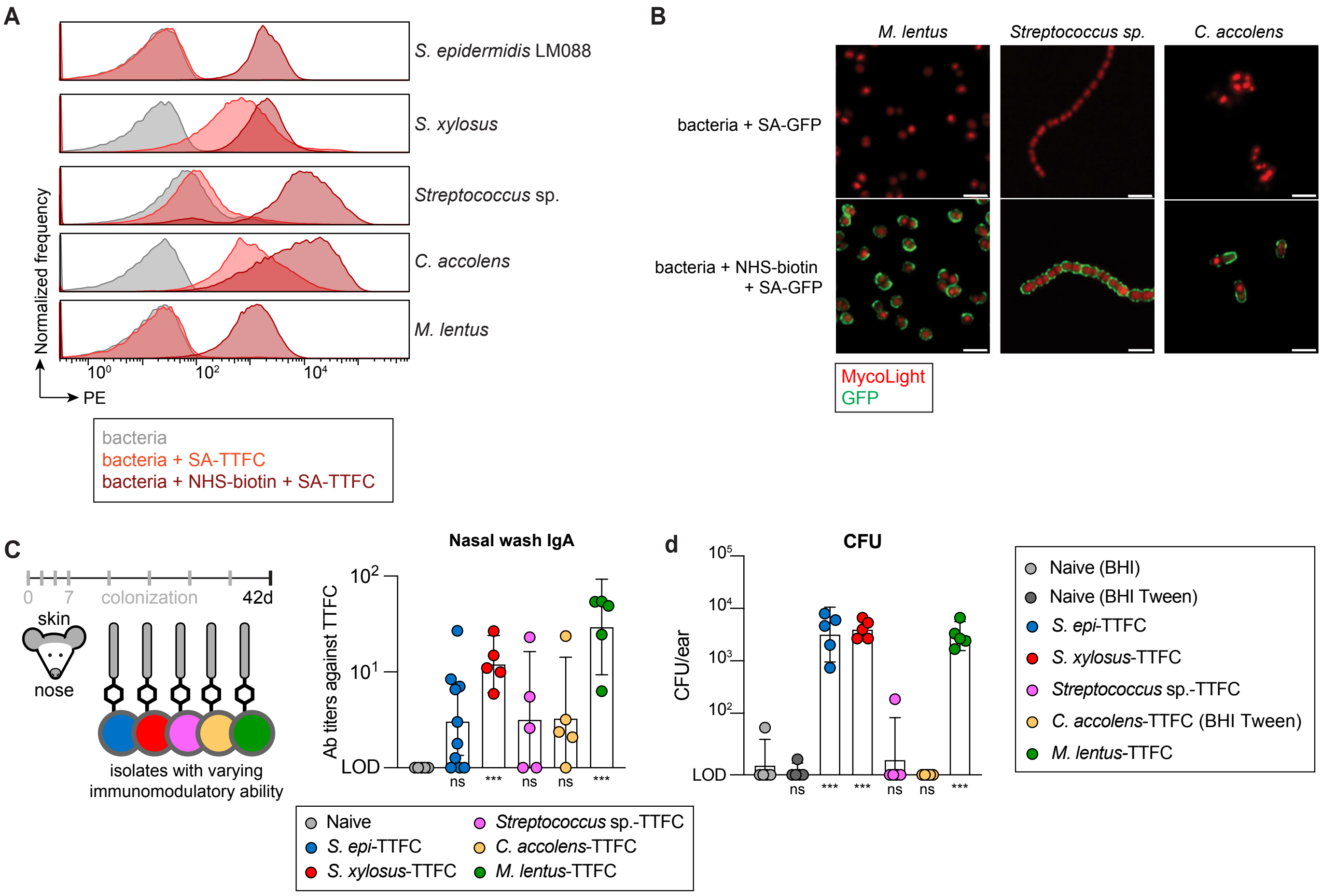
A
S. epidermidis LM088
S. xylosus
Streptococcus sp.
C. accolens
M. lentus
Normalized frequency
$10^0$
$10^2$
$10^4$
PE
bacteria
bacteria + SA-TTFC
bacteria + NHS-biotin + SA-TTFC
B
M. lentus
Streptococcus sp.
C. accolens
bacteria + SA-GFP
bacteria + NHS-biotin + SA-GFP
MycoLight
GFP
C
0
7
colonization
42d
skin
nose
isolates with varying immunomodulatory ability
Nasal wash IgA
Ab titers against TTFC
$10^2$
$10^1$
LOD
ns
***
ns
ns
***
Naive
S. epi-TTFC
S. xylosus-TTFC
Streptococcus sp.-TTFC
C. accolens-TTFC
M. lentus-TTFC
d
CFU
CFU/ear
$10^5$
$10^4$
$10^3$
$10^2$
LOD
ns
***
***
ns
ns
***
Naive (BHI)
Naive (BHI Tween)
S. epi-TTFC
S. xylosus-TTFC
Streptococcus sp.-TTFC
C. accolens-TTFC (BHI Tween)
M. lentus-TTFC

**Figure S5.** Bacterial strains with varying native immunogenicity can be conjugated to TTFC and induce antibody responses in mice.

(**A**) Flow cytometry histograms showing strains from human and mouse hosts with different immunomodulatory activity can be labeled with the model immunogen TTFC using NHS-PEG4-biotin. Labeled bacteria were stained with anti-FLAG-PE to detect attachment, and histograms show PE fluorescence of Syto9$^{+}$ cells for each strain. Gray histograms show bacteria + anti-FLAG-PE, light red histograms indicate no biotin control reactions, and dark red histograms are complete reactions.

(**B**) Confocal microscopy of *M. lentus*, *Streptococcus sp.*, and *C. accolens* labeled with SA-GFP. No biotin reactions included as controls and bacterial cells identified with MycoLight$^{+}$ staining. Scale bar = 2 µm.

(**C**) Data shown are from the same experiment as **Fig. 2C**. Strains were conjugated to TTFC and used to colonize the skin and nose of mice eight times over six weeks, after which serum, nasal wash, and BAL fluid were collected and assessed for TTFC-specific antibodies. Data here show IgA antibodies in the nasal wash specific for TTFC.

(**D**) Data shown here are from the same experiment as **Fig. 2C** and represent CFU/ear from the colonized mice. Ears were harvested after the six-week colonization period, homogenized in PBS, and serial dilutions were plated on BHI or BHI Tween (naive and *C. accolens*) overnight.

Data shown are representative of two (**C, D**) or three (**A, B**) independent experiments. Graphs show geometric mean with a 95% confidence interval. *P* values (ns, not significant; * $p < 0.05$; ** $p < 0.01$; *** $p < 0.001$) were calculated using an ordinary one-way ANOVA followed by Tukey's multiple comparisons test, assuming log-normal distributions. Asterisks in figures represent pairwise comparisons of the indicated group to naive mice (BHI naive for CFU data); *p* values for all comparisons are reported in **Table S1**.

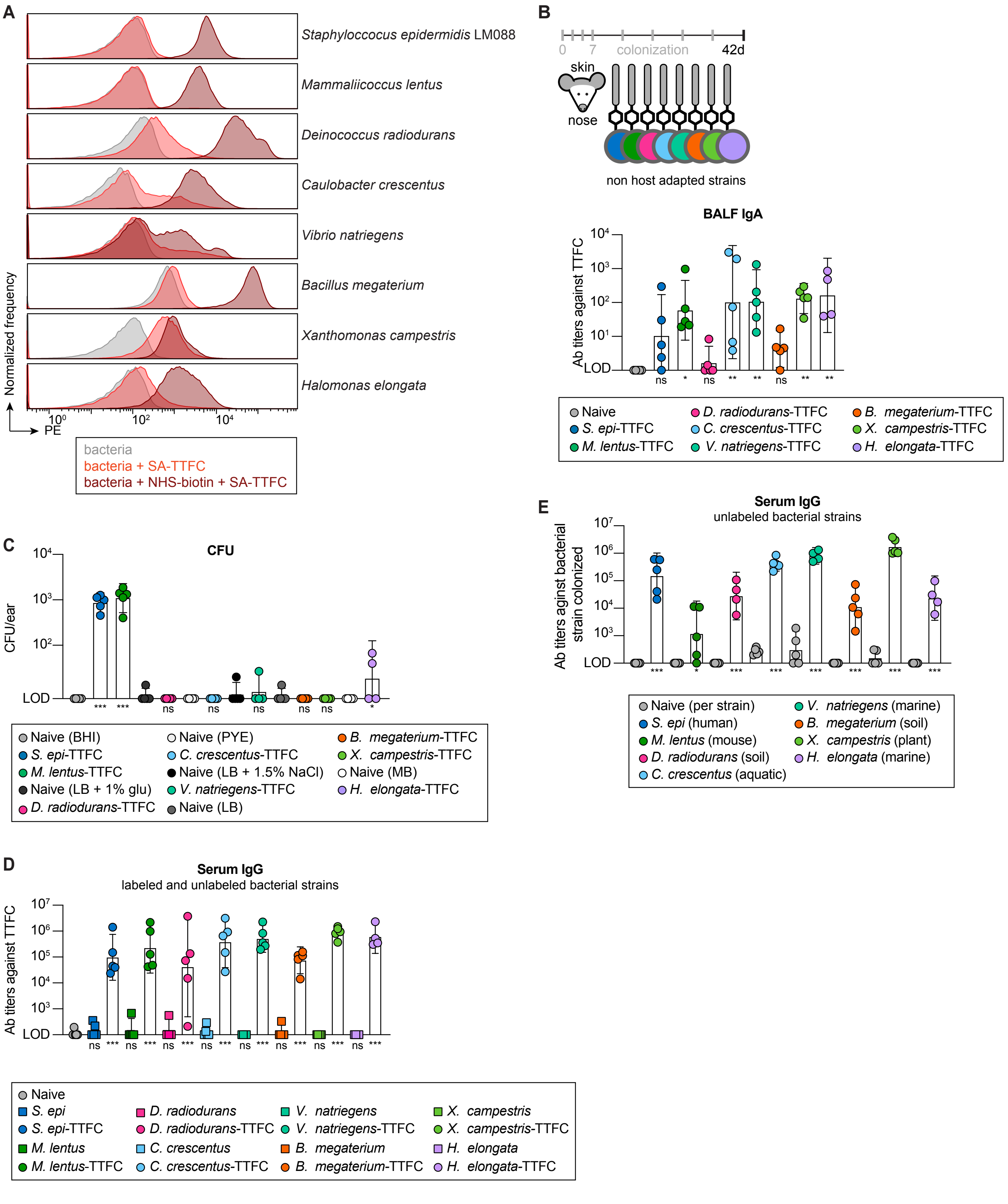
A
Staphyloccocus epidermidis LM088
Mammaliicoccus lentus
Deinococcus radiodurans
Caulobacter crescentus
Vibrio natriegens
Bacillus megaterium
Xanthomonas campestris
Halomonas elongata
Normalized frequency
PE
bacteria
bacteria + SA-TTFC
bacteria + NHS-biotin + SA-TTFC
B
0 7 colonization 42d
skin
nose
non host adapted strains
BALF IgA
Ab titers against TTFC
Naive
S. epi-TTFC
M. lentus-TTFC
D. radiodurans-TTFC
C. crescentus-TTFC
V. natriegens-TTFC
B. megaterium-TTFC
X. campestris-TTFC
H. elongata-TTFC
C
CFU
CFU/ear
Naive (BHI)
S. epi-TTFC
M. lentus-TTFC
Naive (LB + 1% glu)
D. radiodurans-TTFC
Naive (PYE)
C. crescentus-TTFC
Naive (LB + 1.5% NaCl)
V. natriegens-TTFC
Naive (LB)
B. megaterium-TTFC
X. campestris-TTFC
Naive (MB)
H. elongata-TTFC
D
Serum IgG
labeled and unlabeled bacterial strains
Ab titers against TTFC
Naive
S. epi
S. epi-TTFC
M. lentus
M. lentus-TTFC
D. radiodurans
D. radiodurans-TTFC
C. crescentus
C. crescentus-TTFC
V. natriegens
V. natriegens-TTFC
B. megaterium
B. megaterium-TTFC
X. campestris
X. campestris-TTFC
H. elongata
H. elongata-TTFC
E
Serum IgG
unlabeled bacterial strains
Ab titers aginst bacterial strain colonized
Naive (per strain)
S. epi (human)
M. lentus (mouse)
D. radiodurans (soil)
C. crescentus (aquatic)
V. natriegens (marine)
B. megaterium (soil)
X. campestris (plant)
H. elongata (marine)

**Figure S6.** Bacterial strains isolated from a variety of environments can be labeled with TTFC and used to induce antibody responses in mice.

(**A**) Flow cytometry histograms showing diverse strains from the environment can be labeled with the model immunogen TTFC using NHS-PEG4-biotin. Labeled bacteria were stained with anti-FLAG-PE to detect attachment, and histograms show PE fluorescence of Syto9$^+$ cells for each strain. Gray histograms show bacteria + anti-FLAG-PE, light red histograms indicate no biotin control reactions, and dark red histograms are complete reactions.

(**B**) Data are from the same experiment shown in **Fig. 2D.** A panel of environmental isolates, along with *S. epidermidis* and *M. lentus* for comparison, were individually conjugated to TTFC and used to colonize the skin and nose of mice eight times over six weeks. Serum, nasal wash, and BAL fluid were subsequently harvested and assessed by ELISA for antibody titers.

(**C**) Expanded CFU/ear data shown in **Fig. 2D**. Ears from naive mice were harvested, homogenized, and plated in triplicate on each growth media for each strain used in this experiment. Ears from colonized mice were plated on their appropriate growth media. LOD = 6.67.

(**D**) Anti-TTFC antibody titers in serum from mice colonized with TTFC-conjugated bacterial strains and unconjugated strains (i.e. wild type bacteria). Serum was collected after six weeks of colonization and assessed for titers against TTFC purified from mammalian cells by ELISA. Data from labeled strains is the same as shown in **Fig. 2D.** Unlabeled, wild-type strains elicit no detectable anti-TTFC response, indicating no background cross-reactivity to the bacterial strain itself.

(**E**) Antibody titers against the bacterial strain used to colonize mice to examine the native immunomodulatory activity of environmental isolates. Mice were colonized with unlabeled (wild-type) bacteria from a variety of different environments eight times over six weeks. Serum was harvested and assessed by ELISA for antibody titers against the bacterial strain itself in comparison to serum from naive mice. Naive titers were evaluated individually against each strain depicted to its right.

Data shown are from one (**B, C, D**) or representative of three (**A**) independent experiments. Graphs show geometric mean with a 95% confidence interval. *P* values (ns, not significant; * $p < 0.05$; ** $p < 0.01$; *** $p < 0.001$) were calculated using an ordinary one-way ANOVA followed by Tukey's multiple comparisons test, assuming log-normal distributions except for bacterial ELISAs in **E.** Asterisks in **B** and **D** represent pairwise comparisons of the indicated group to naive mice, while asterisks shown in **C** represent comparison of naive and colonized mice per growth medium. For bacterial ELISAs in **E,** log-transformed titers from naive and colonized mice were compared for each strain using unpaired two-tailed *t* tests with the Holm-Šídák correction for multiple comparisons ($\alpha = 0.05$), and asterisks represent these internal comparisons. All *p* values are reported in **Table S1**.

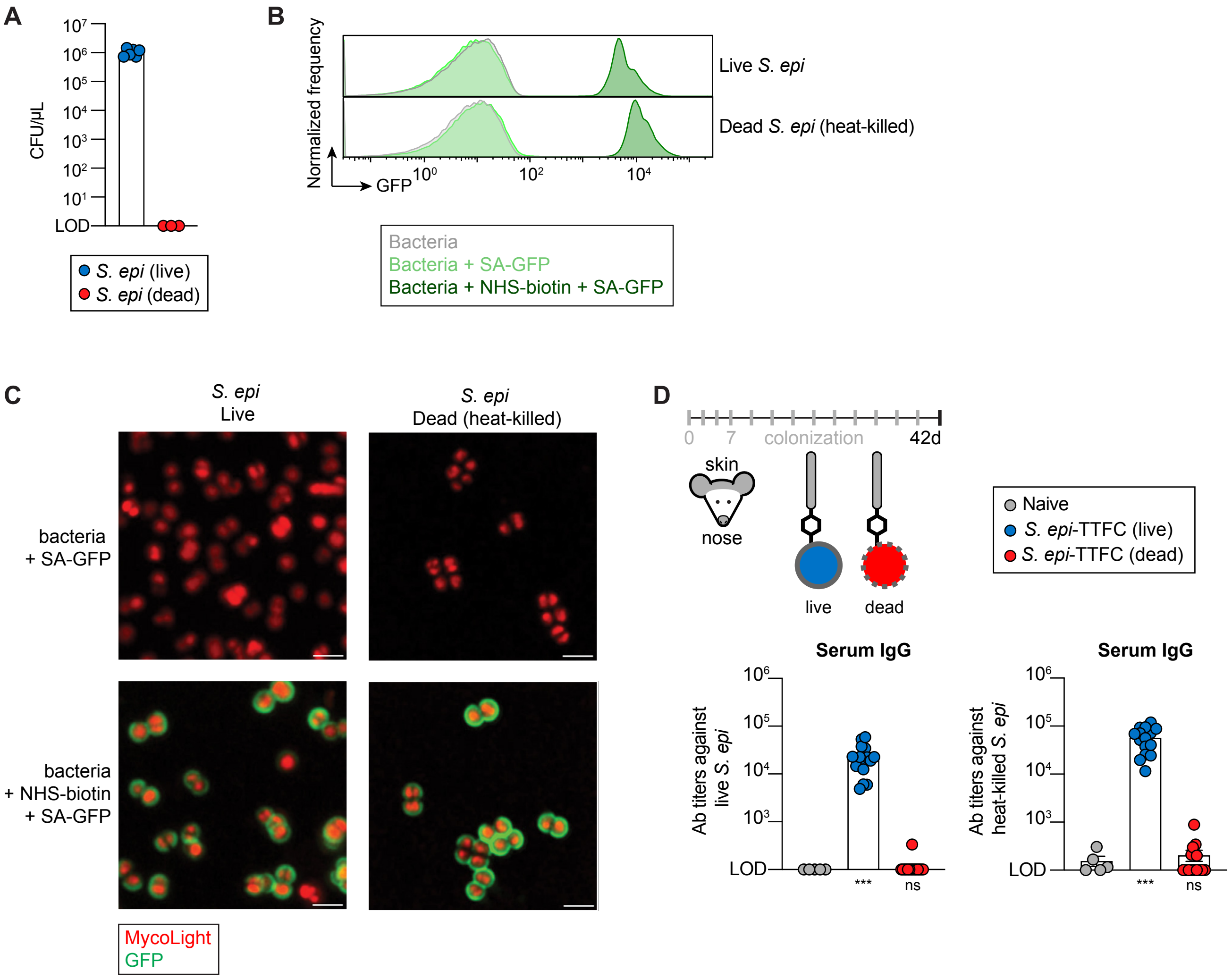
A
CFU/μL
$10^7$
$10^6$
$10^5$
$10^4$
$10^3$
$10^2$
$10^1$
LOD
S. epi (live)
S. epi (dead)
B
Normalized frequency
GFP
$10^0$
$10^2$
$10^4$
Live S. epi
Dead S. epi (heat-killed)
Bacteria
Bacteria + SA-GFP
Bacteria + NHS-biotin + SA-GFP
C
S. epi
Live
S. epi
Dead (heat-killed)
bacteria
+ SA-GFP
bacteria
+ NHS-biotin
+ SA-GFP
MycoLight
GFP
D
0
7
colonization
42d
skin
nose
live
dead
Naive
S. epi-TTFC (live)
S. epi-TTFC (dead)
Serum IgG
Ab titers against live S. epi
$10^6$
$10^5$
$10^4$
$10^3$
LOD
***
ns
Serum IgG
Ab titers against heat-killed S. epi
$10^6$
$10^5$
$10^4$
$10^3$
LOD
***
ns

**Figure S7.** Heat-killed *S. epidermidis* can be efficiently surface labeled with proteins.

(**A**) Bacterial recovery of *S. epidermidis* before and after heat-killing. Cells plated on BHI media. Data depicted as CFU/µL.

(**B**) Flow cytometry histograms showing live and dead *S. epidermidis* can be labeled with GFP using NHS-PEG4-biotin. *S. epidermidis* was heat-killed prior to labeling. Histograms show GFP fluorescence of MycoLight$^{+}$ cells for both strains. Gray histograms show bacteria, light green histograms indicate no biotin control reactions, and dark green histograms are complete reactions.

(**C**) Confocal microscopy of live and dead *S. epidermidis* labeled with SA-GFP. Bacterial cells detected with MycoLight. Scale bar = 2 µm.

(**D**) Antibody titers in serum from mice colonized with live or dead *S. epidermidis*- TTFC against live (left) and dead (right) *S. epidermidis*. Data from the same experiment shown in **Fig. 2E.** Mice colonized with live *S. epidermidis* have antibody titers against both live and dead *S. epidermidis*, while mice colonized with dead *S. epidermidis* have no antibody titers against the bacterial strain.

Data shown are from or representative of two (**D**) or three (**A, B, C**) independent experiments. Graphs show geometric mean with a 95% confidence interval. *P* values (ns, not significant; * $p < 0.05$; ** $p < 0.01$; *** $p < 0.001$) were calculated using an ordinary one-way ANOVA followed by Tukey's multiple comparisons test, assuming log-normal distributions**.** Asterisks in **D** represent pairwise comparisons of the indicated group to naive mice; *p* values for all comparisons are reported in **Table S1**.

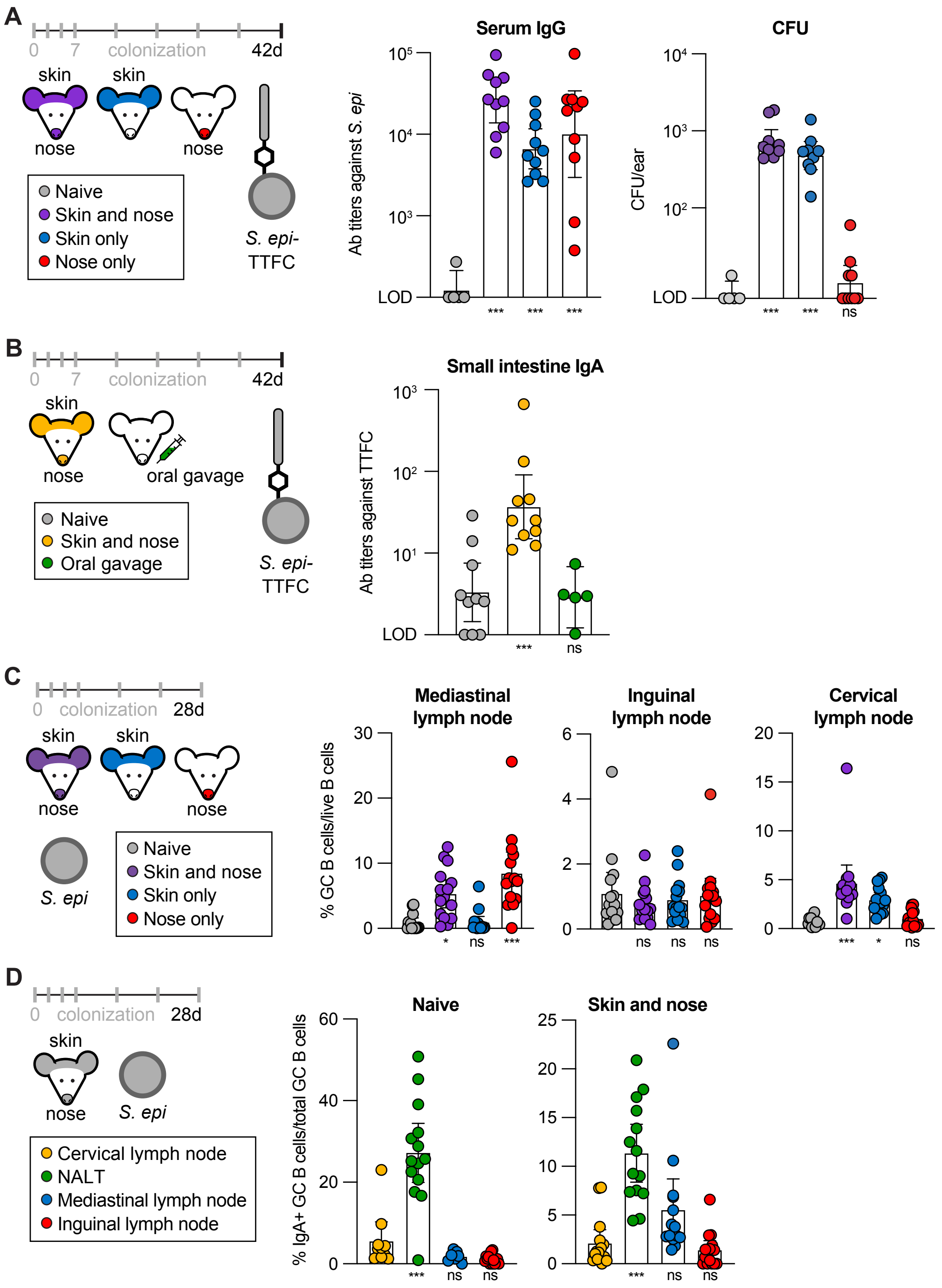
A
0
7
colonization
42d
skin
skin
nose
nose
Naive
Skin and nose
Skin only
Nose only
S. epi-
TTFC
Serum IgG
Ab titers against S. epi
LOD
CFU
CFU/ear
B
skin
nose
oral gavage
Naive
Skin and nose
Oral gavage
Small intestine IgA
Ab titers against TTFC
C
28d
S. epi
Mediastinal
lymph node
Inguinal
lymph node
Cervical
lymph node
% GC B cells/live B cells
D
Cervical lymph node
NALT
Mediastinal lymph node
Inguinal lymph node
Naive
Skin and nose
% IgA+ GC B cells/total GC B cells

**Figure S8.** Nasal colonization induces an IgA response at mucosal sites.

(**A**) Serum IgG endpoint titers against *S. epidermidis*, measured using ELISA, and CFU recovered per mouse ear, plated on BHI media, for the experiment shown in **Fig. 3A**.

(**B**) Mice were either colonized on the skin and nose with S. epi-TTFC or fed the TTFC-conjugated bacteria by oral gavage, eight times over six weeks. Small intestinal contents were then harvested from these mice and analyzed for TTFC-specific IgA antibodies by ELISA.

(**C**) Data depicted are from the same experiment as **Fig. 3B**. All flow cytometry data shown are from the day 28 time point.

(**D**) Data displayed are from the same experiment as **Fig. 3C**. Mice were either naive or colonized with wild-type *S. epidermidis* on the skin and nose using a swab, six times over four weeks.

Data shown are representative of at least two independent experiments. Graphs show mean (**C, D**) or geometric mean (**A, B**) with a 95% confidence interval. *P* values (ns, not significant; * $p < 0.05$; ** $p < 0.01$; *** $p < 0.001$) were calculated using an ordinary one-way ANOVA followed by Tukey's multiple comparisons test, assuming normal (**C, D**) or log-normal (**A, B**) distributions. Asterisks in figures represent pairwise comparisons of the indicated group to naive mice (**A**, **B**, **C**) or cervical lymph node samples (**D**); *p* values for all comparisons are reported in **Table S1**.

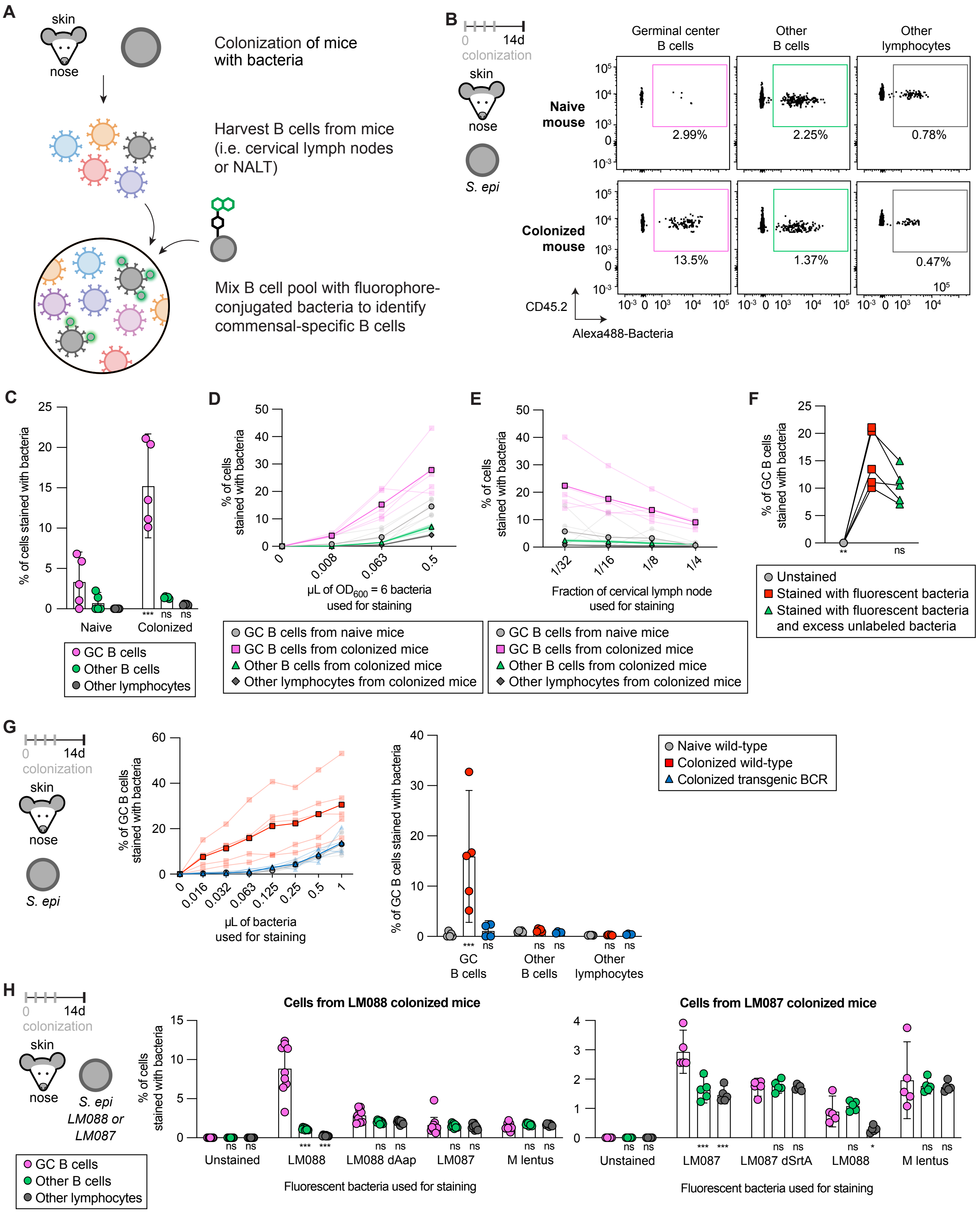

A
skin
nose
Colonization of mice with bacteria
Harvest B cells from mice (i.e. cervical lymph nodes or NALT)
Mix B cell pool with fluorophore-conjugated bacteria to identify commensal-specific B cells
B
0 14d colonization
S. epi
Germinal center B cells
Other B cells
Other lymphocytes
Naive mouse
Colonized mouse
2.99%
2.25%
0.78%
13.5%
1.37%
0.47%
CD45.2
Alexa488-Bacteria
C
% of cells stained with bacteria
Naive
Colonized
GC B cells
Other B cells
Other lymphocytes
D
% of cells stained with bacteria
μL of OD600 = 6 bacteria used for staining
GC B cells from naive mice
GC B cells from colonized mice
Other B cells from colonized mice
Other lymphocytes from colonized mice
E
% of cells stained with bacteria
Fraction of cervical lymph node used for staining
GC B cells from naive mice
GC B cells from colonized mice
Other B cells from colonized mice
Other lymphocytes from colonized mice
F
% of GC B cells stained with bacteria
Unstained
Stained with fluorescent bacteria
Stained with fluorescent bacteria and excess unlabeled bacteria
G
0 14d colonization
S. epi
% of GC B cells stained with bacteria
μL of bacteria used for staining
% of GC B cells stained with bacteria
GC B cells
Other B cells
Other lymphocytes
Naive wild-type
Colonized wild-type
Colonized transgenic BCR
H
0 14d colonization
S. epi LM088 or LM087
GC B cells
Other B cells
Other lymphocytes
Cells from LM088 colonized mice
% of cells stained with bacteria
Unstained
LM088
LM088 dAap
LM087
M lentus
Fluorescent bacteria used for staining
Cells from LM087 colonized mice
Unstained
LM087
LM087 dSrtA
LM088
M lentus
Fluorescent bacteria used for staining

**Figure S9.** Fluorescent *S. epidermidis* can be used as a staining reagent to identify bacteria-specific B cells.

(**A**) Schematic depicting strategy for identifying commensal-specific B cells using fluorophore-conjugated bacteria as a staining reagent.

(**B**) Representative flow cytometry data demonstrating that fluorescently labeled bacteria specifically identify germinal center B cells reactive to *S. epidermidis*. Mice were colonized with wild-type *S. epidermidis* four times by swabbing the skin and nose. After 14 days, cervical lymph nodes were harvested and stained. The percentage of germinal center B cells (Live CD45.2$^{+}$ CD19$^{+}$ CD38$^{-}$ Fas$^{+}$), other B cells (Live CD45.2$^{+}$ CD19$^{+}$ CD38$^{+}$ Fas$^{-}$), and other lymphocytes (Live CD45.2$^{+}$ CD19$^{-}$) that were bacteria-positive was quantified by flow cytometry.

(**C**) Quantification of data from the same experiment as **B**.

(**D**) Data from the same experiment as **B**, where mouse cells (consisting of ⅛ of a cervical lymph node preparation) were stained with different volumes of bacteria, as indicated.

(**E**) Data from the same experiment as **B**, where different fractions of a cervical lymph node sample, as indicated, were each stained with 0.0625 µL of $OD_{600}$ = 6 of fluorescent *S. epidermidis*.

(**F**) Data from the same experiment as **B**, where cervical lymph node cells were stained with 0.0625 µL fluorescent *S. epidermidis*, with or without 2 µL of $OD_{600}$ = 6 unlabeled *S. epidermidis*.

(**G**) Over two weeks, wild-type or transgenic mice engineered to solely express a foreign B cell receptor targeting hen egg lysozyme were colonized four times with wild-type *S. epidermidis* using a swab to apply culture to the skin and nose. One group of mice was kept naive. At the experimental endpoint, cells from cervical lymph nodes were harvested, stained with the indicated volume of AlexaFluor488-conjugated *S. epidermidis*, and analyzed by flow cytometry to quantify the percentage of bacteria-stained germinal center B cells (Live CD45.2$^{+}$ CD19$^{+}$ CD38$^{-}$ Fas$^{+}$), other B cells (Live CD45.2$^{+}$ CD19$^{+}$ CD38$^{+}$ Fas$^{-}$), or other lymphocytes (Live CD45.2$^{+}$ CD19$^{-}$).

(**H**) Mice were either colonized with *S. epidermidis* LM088 (left) or LM087 (right) by swabbing their skin and nose four times over the course of 14 days. At the experimental endpoint, cells from lymph nodes were harvested and stained with the indicated strains of bacteria conjugated to AlexaFluor488. The percentage of cells that stained with bacteria was quantified for bacteria-stained germinal center B cells (Live CD45.2$^{+}$ CD19$^{+}$ CD38$^{-}$ Fas$^{+}$), other B cells (Live CD45.2$^{+}$ CD19$^{+}$ CD38$^{+}$ Fas$^{-}$), or other lymphocytes (Live CD45.2$^{+}$ CD19$^{-}$).

Data shown are from one (**F**, **G**) or representative of two (**B**, **C**, **D**, **E**, **H**) independent experiments. Graphs show mean with a 95% confidence interval. *P* values (ns, not significant; * $p < 0.05$; ** $p < 0.01$; *** $p < 0.001$) were calculated using a two-way ANOVA (**C**, **G**, **H**) or repeated measures one-way ANOVA (**f**) followed by Tukey's multiple comparisons test. Asterisks in figures represent pairwise comparisons of the indicated group to naive mice within the corresponding cell type (**C**, **G**), to the "stained with fluorescent bacteria” group (**F**), or to the “germinal center B cells” group within each fluorescent strain (**H**); *p* values for all comparisons are reported in **Table S1**.

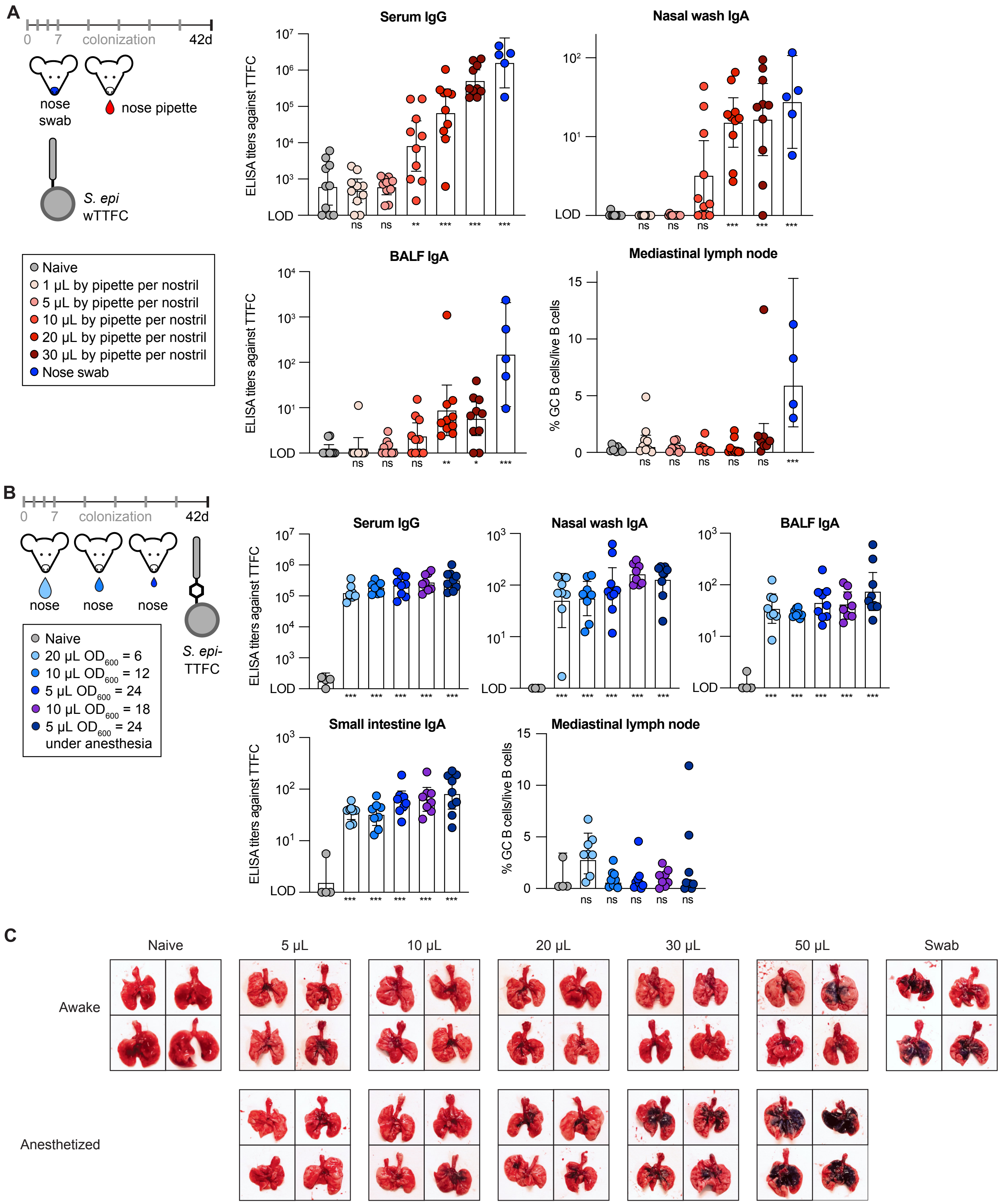
A
0
7
colonization
42d
nose swab
nose pipette
S. epi wTTFC
Naive
1 μL by pipette per nostril
5 μL by pipette per nostril
10 μL by pipette per nostril
20 μL by pipette per nostril
30 μL by pipette per nostril
Nose swab
Serum IgG
Nasal wash IgA
BALF IgA
Mediastinal lymph node
ELISA titers against TTFC
% GC B cells/live B cells
LOD
ns
B
0
7
colonization
42d
nose
nose
nose
S. epi-TTFC
Naive
20 μL OD600 = 6
10 μL OD600 = 12
5 μL OD600 = 24
10 μL OD600 = 18
5 μL OD600 = 24 under anesthesia
Serum IgG
Nasal wash IgA
BALF IgA
Small intestine IgA
Mediastinal lymph node
ELISA titers against TTFC
% GC B cells/live B cells
LOD
C
Naive
5 μL
10 μL
20 μL
30 μL
50 μL
Swab
Awake
Anesthetized

**Figure S10.** Low-volume intranasal delivery elicits immune responses without draining to the lungs. (**A**) *S. epidermidis* genetically engineered to express TTFC was used to colonize mice either by nose swab or by pipetting a range of volumes into the nostrils. After eight colonizations administered during the course of six weeks, mice were euthanized, and serum, nasal wash, and BAL fluid were harvested and analyzed for the presence of TTFC-specific antibodies by ELISA. Additionally, mediastinal lymph nodes (which drain the lungs) were dissected, and the percentage of germinal center B cells ($CD38^-$ $Fas^+$) among total live B cells (Live $CD45.2^+$ $CD19^+$) was quantified using flow cytometry. Data shown are pooled from two independent experiments.

(**B**) Data are from the same experiment as **Fig. 3E**. The indicated volumes of *S. epi*-TTFC were pipetted into each nostril of mice. After eight colonizations in six weeks, serum, nasal wash, BAL fluid, and small intestinal contents were collected, and TTFC-specific antibody endpoint titers were determined by ELISA. As above, the mediastinal lymph nodes of the mice were harvested, and the percentage of total live B cells (Live $CD45.2^+$ $CD19^+$) that were germinal center B cells ($CD38^-$ $Fas^+$) was calculated by flow cytometry.

(**C**) Evans blue dye was mixed with *S. epidermidis* culture and intranasally administered to mice by either pipetting (doses shown are volume per nostril) or by swab. Groups of mice were either awake or anesthetized during colonization, as indicated. Mice were euthanized, and the dissected trachea and lungs are shown.

Data shown are representative of at least two independent experiments. Graphs show mean (mediastinal lymph node in **A** and **B**) or geometric mean (**A**, **B**) with a 95% confidence interval. *P* values (ns, not significant; * $p < 0.05$; ** $p < 0.01$; *** $p < 0.001$) were calculated using an ordinary one-way ANOVA followed by Tukey's multiple comparisons test, assuming normal (mediastinal lymph node in **A** and **B**) or log-normal distributions (**A**, **B**). Asterisks in figures represent pairwise comparisons of the indicated group to naive mice; *p* values for all comparisons are reported in **Table S1**.

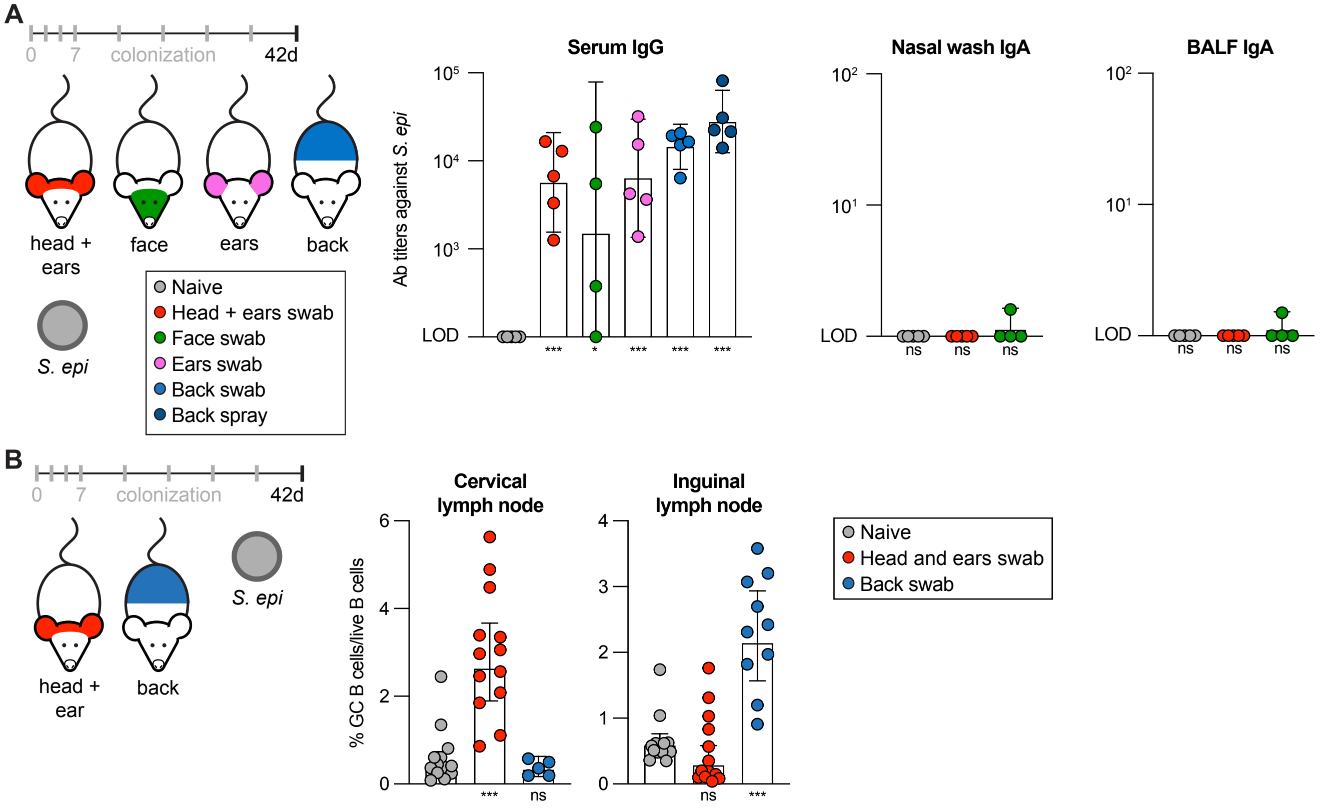
A
0
7
colonization
42d
head + ears
face
ears
back
S. epi
Naive
Head + ears swab
Face swab
Ears swab
Back swab
Back spray
Serum IgG
Ab titers against S. epi
LOD
Nasal wash IgA
BALF IgA
ns
B
head + ear
back
Cervical lymph node
Inguinal lymph node
% GC B cells/live B cells
Naive
Head and ears swab
Back swab

**Figure S11.** Several skin sites are competent to elicit immunity after colonization.

(**A**) Mice were either colonized with *S. epidermidis* using a swab to apply bacteria on the head and ears, face, ears, or back. An additional group of mice was colonized by spraying culture onto the lower back using a mucosal atomization device (MAD). A total of eight colonizations were administered over six weeks, after which serum, nasal wash, and BAL fluid were harvested and analyzed for the presence of antibodies recognizing *S. epidermidis* by ELISA.

(**B**) *S. epidermidis* was applied to the head and ears or back of mice with a cotton swab, eight total times. After six weeks, the cervical and inguinal lymph nodes were harvested, and analyzed by flow cytometry to quantify the percentage of live B cells (Live $CD45.2^+$ $CD19^+$) that were germinal center B cells ($CD38^-$ $Fas^+$). Data shown are pooled from two independent experiments.

(**C**) Wild-type or MHC-II knockout mice were colonized with *S. epidermidis* genetically engineered to express TTFC, on the skin and nose by means of a swab. After eight colonizations to the mice administered over six weeks, levels of TTFC- or bacteria-specific antibodies in the serum, nasal wash, and BAL fluid were analyzed by ELISA. Additionally, bacteria recovered from the ears of each mouse were plated on media for CFU quantification.

Data shown are from one (**C**) or representative of two (**A**, **B**) independent experiments. Graphs show mean (**B**) or geometric mean (**A, C**) with a 95% confidence interval. *P* values (ns, not significant; * $p < 0.05$; ** $p < 0.01$; *** $p < 0.001$) were calculated using an ordinary one-way ANOVA followed by Tukey's multiple comparisons test (**A**, **B**) or two-way ANOVA with Fisher's LSD test (**C**), assuming normal (**B**) or log-normal (**A, C**) distributions. Asterisks in figures represent pairwise comparisons of the indicated group to naive mice within the corresponding group (**A**, **B**, **C**); *p* values for all comparisons are reported in **Table S1**.

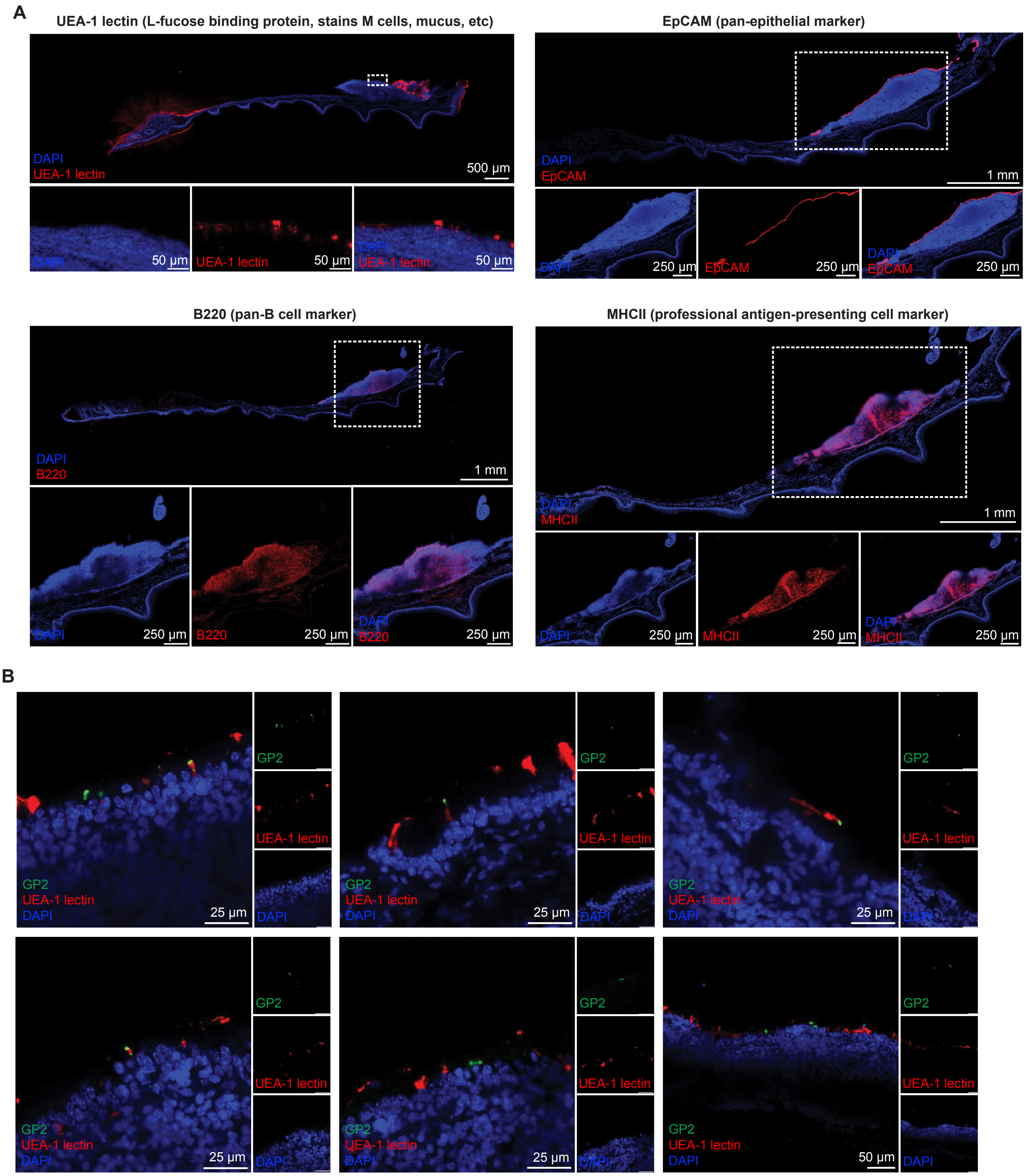
A
UEA-1 lectin (L-fucose binding protein, stains M cells, mucus, etc)
DAPI
UEA-1 lectin
500 µm
50 µm
EpCAM (pan-epithelial marker)
DAPI
EpCAM
1 mm
250 µm
B220 (pan-B cell marker)
DAPI
B220
1 mm
250 µm
MHCII (professional antigen-presenting cell marker)
DAPI
MHCII
1 mm
250 µm
B
GP2
UEA-1 lectin
DAPI
25 µm
50 µm

C

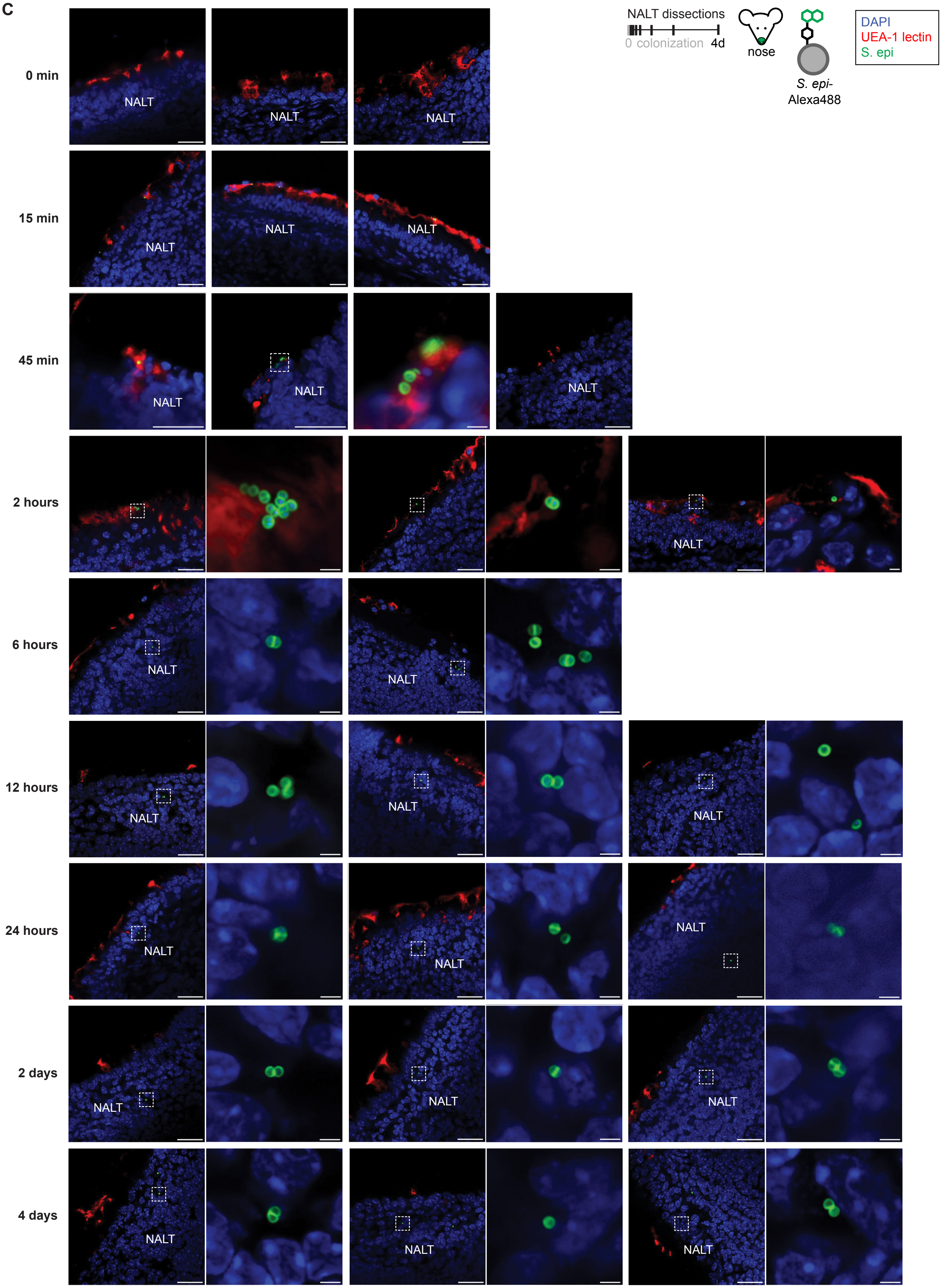

NALT dissections
0 colonization 4d
nose
S. epi-
Alexa488
DAPI
UEA-1 lectin
S. epi
0 min
15 min
45 min
2 hours
6 hours
12 hours
24 hours
2 days
4 days
NALT

**Figure S12.** Whole bacteria are present in the NALT after nasal colonization.

(**A**) Confocal microscopy of the whole palate, including the NALT, stained with different markers by immunofluorescence performed on 100 μm sections. Sections are stained in blue with DAPI, and in red with UEA-1 lectin (top left), EpCAM (top right), B220 (bottom left), or MHC-II (bottom right). Boxed regions are shown with higher magnification below each overview. Z-stack maximum projection images were acquired by tile scan and stitched together. Scale bars are the indicated sizes.

(**B**) Confocal microscopy images of the follicle-associated epithelium lining the NALT surface, stained for DAPI (blue), the mature M cell marker GP2 (green), and UEA-1 lectin (red). Each panel shows the merged image alongside the separated single-color channels. Scale bars are the indicated sizes.

(**C**) Imaging data from the same experiment shown in **Fig. 3F**. Mice were colonized with *S. epi* conjugated to AlexaFluor488 (green) by nasal swab, and the NALT was harvested from each mouse at the indicated times after a single colonization. 100 μm sections of tissue were stained with DAPI (blue) and UEA-1 lectin (red) before being imaged by confocal microscopy. Insets show magnified views of the boxed regions, where indicated, in which individual bacteria are resolved with clear morphology. Scale bars are 25 μm (overviews) or 2 μm (insets).

Data shown are representative of images from at least n = 3 mice per group, from one (**B**) or at least two (**A**, **C**) independent experiments.

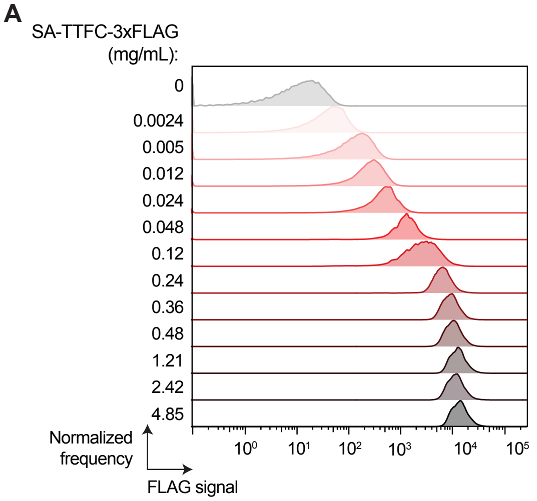


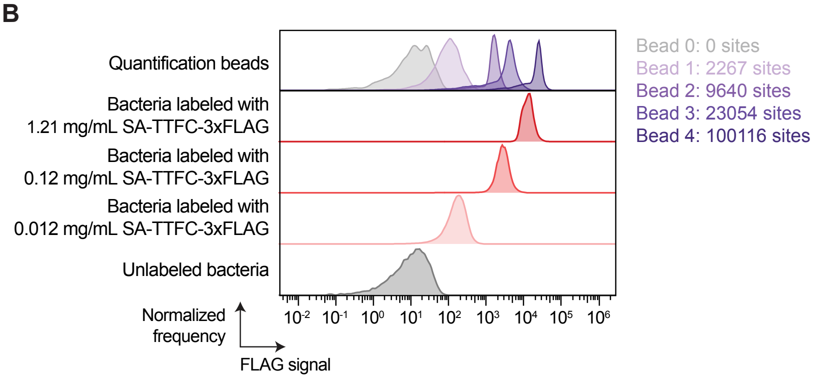


| Dose | mg/mL SA-TTFC used in reaction | μg/mL SA-TTFC on cell surface after reaction | Molecules of SA-TTFC per bacterial cell after reaction |
|---|---|---|---|
| High | 1.21 | 7.07 | 56395 |
| Medium | 0.12 | 2.56 | 19359 |
| Low | 0.012 | 0.34 | 2813 |

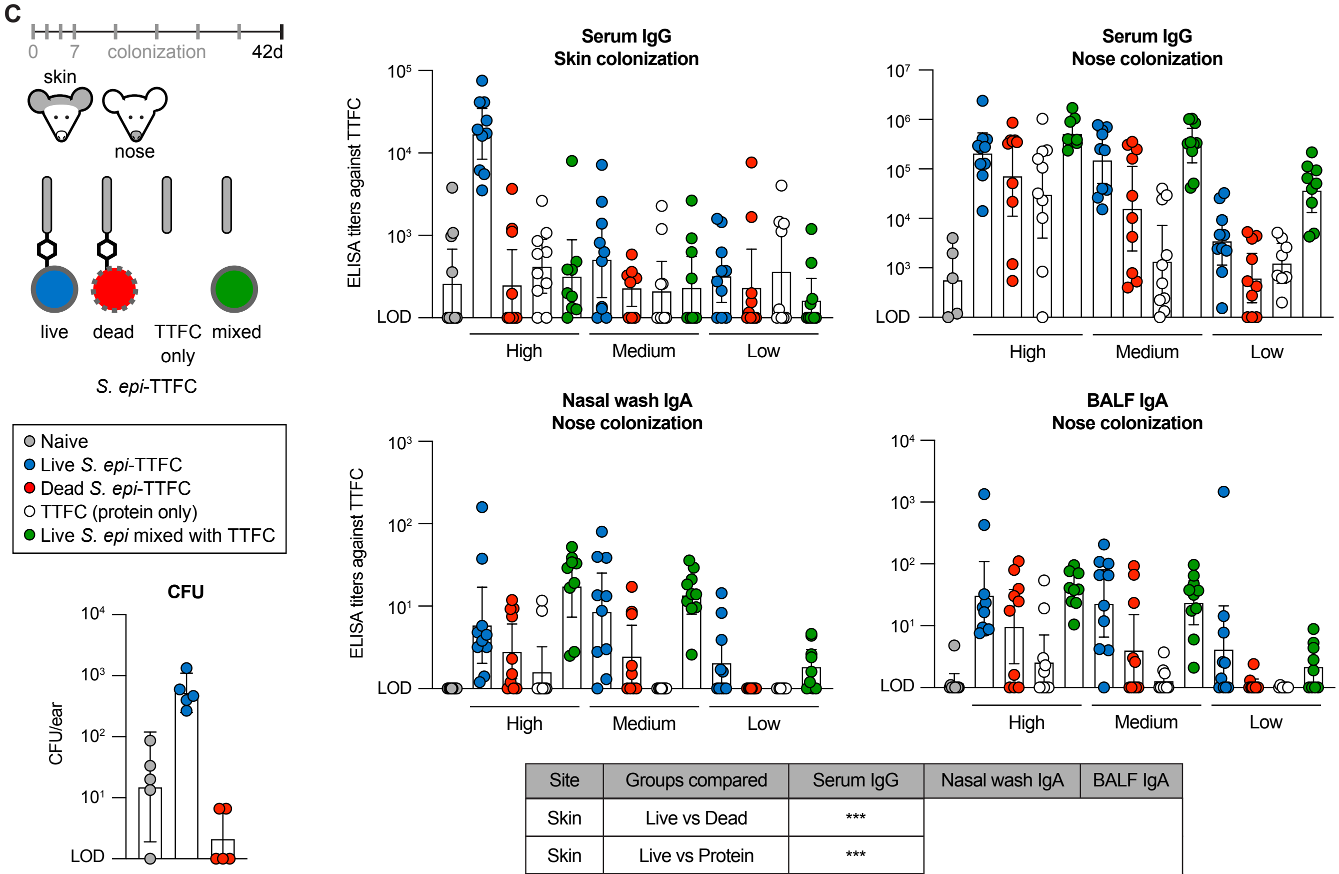


| Site | Groups compared | Serum IgG | Nasal wash IgA | BALF IgA |
|---|---|---|---|---|
| Skin | Live vs Dead | *** | | |
| Skin | Live vs Protein | *** | | |
| Skin | Live vs Mixed | *** | | |
| Skin | Protein vs Dead | ns | | |
| Skin | Protein vs Mixed | ns | | |
| Nose | Live vs Dead | ** | ** | ** |
| Nose | Live vs Protein | *** | *** | *** |
| Nose | Live vs Mixed | * | ns | ns |
| Nose | Protein vs Dead | ns | ns | ns |
| Nose | Protein vs Mixed | *** | *** | *** |

**Figure S13.** Mechanistic requirements for eliciting antibody responses after topical or intranasal colonization.

(**A**) Flow cytometry histograms of *S. epidermidis* conjugated to TTFC at a range of concentrations, detected by anti-FLAG staining. The displayed values indicate the concentrations of SA-TTFC added to the reaction.

(**B**) Absolute quantification of antigen number on the surface of *S. epidermidis* after conjugation. Bacteria attached to three different doses of antigen or unlabeled controls were stained with anti-FLAG antibody and analyzed by flow cytometry alongside calibration beads bearing defined numbers of antibody-binding sites. Median fluorescence intensity for each peak was calculated; the beads were used to create a standard curve, against which the bacterial samples were interpolated to quantify the number of molecules per cell.

(**C**) Data shown are from the same experiment as **Fig 3G**. The three doses of antigen used here correspond to the doses quantified in **B**. At the experimental endpoint, serum was harvested from mice colonized on the skin, and serum, nasal wash, and BAL fluid were harvested from mice colonized on the nose. The levels of TTFC-specific antibodies were measured in each analyte by ELISA. CFU recovered per mouse ear are shown for naive, live *S. epi*-TTFC, and dead *S. epi*-TTFC groups at the high dose.

Data shown are representative of at least two independent experiments. Graphs show geometric mean with a 95% confidence interval. *P* values (ns, not significant; * $p < 0.05$; ** $p < 0.01$; *** $p < 0.001$) were calculated using a two-way ANOVA followed by Tukey's multiple comparisons test performed between each test article, assuming log-normal distributions; *p* values for all comparisons are reported in **Table S1**.

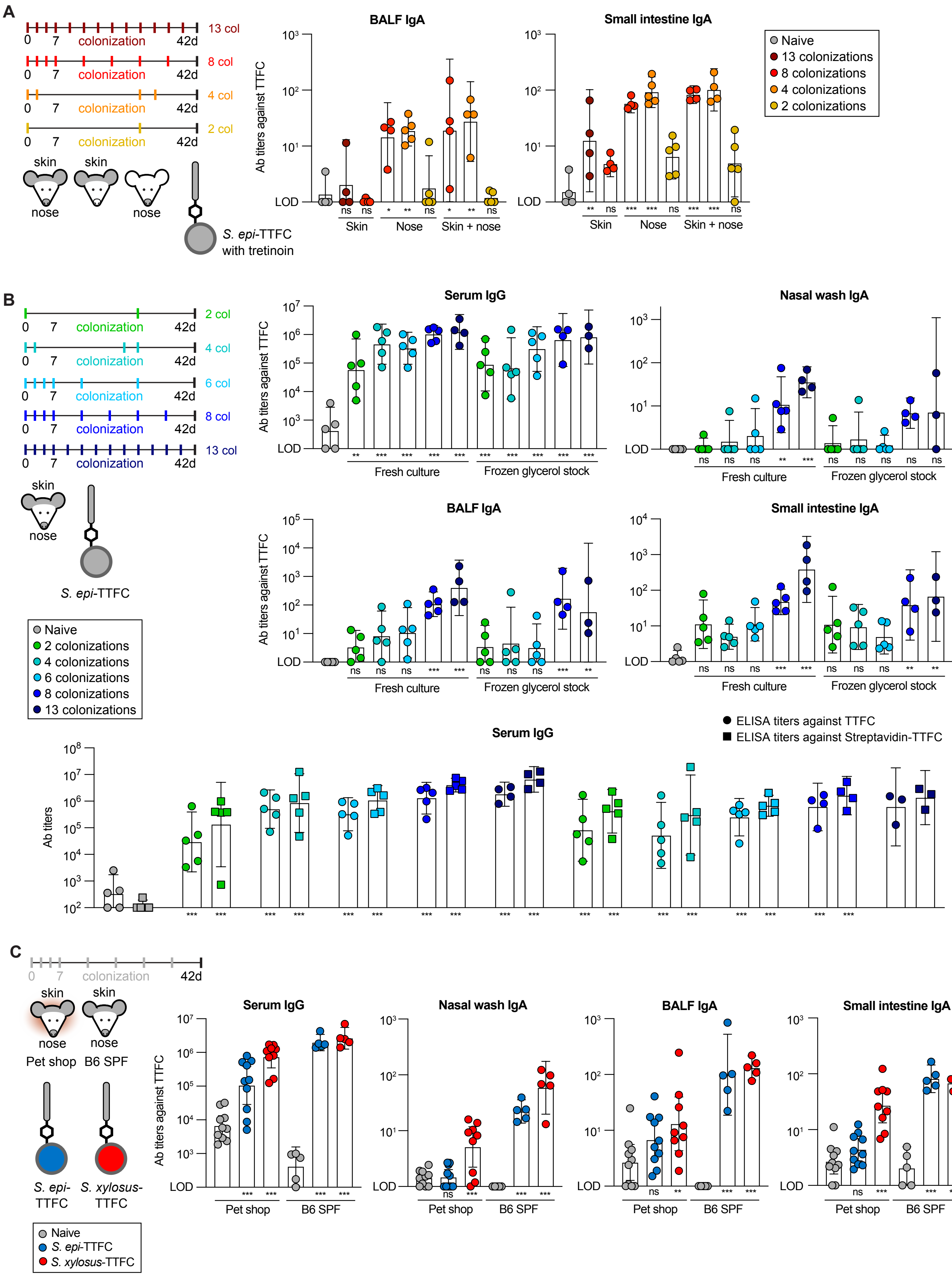
A
13 col
8 col
4 col
2 col
0 7 colonization 42d
skin
nose
S. epi-TTFC with tretinoin
BALF IgA
Small intestine IgA
Ab titers against TTFC
LOD
Skin
Nose
Skin + nose
Naive
13 colonizations
8 colonizations
4 colonizations
2 colonizations
B
2 col
4 col
6 col
8 col
13 col
S. epi-TTFC
Serum IgG
Nasal wash IgA
BALF IgA
Small intestine IgA
Fresh culture
Frozen glycerol stock
Naive
2 colonizations
4 colonizations
6 colonizations
8 colonizations
13 colonizations
ELISA titers against TTFC
ELISA titers against Streptavidin-TTFC
Ab titers
C
Pet shop
B6 SPF
S. epi-TTFC
S. xylosus-TTFC
Naive
S. epi-TTFC
S. xylosus-TTFC

**Figure S14.** Compressed dosing regimens and activity in pet shop mice.

(**A**) Data shown are from the same experiment as **Fig. 4A**, in which mice were colonized with *S. epi*-TTFC plus tretinoin on the skin, nose, or both, across 2 to 13 colonizations. The levels of TTFC-specific antibodies in BAL fluid and small intestinal contents were measured using ELISA.

(**B**) Mice were colonized according to several different colonization schedules by applying either fresh culture or frozen glycerol stocks of *S. epi*-TTFC to the skin and nose using a swab. The various colonization timelines used are: two colonizations on days 0 and 28; four colonizations on days 0, 2, 28, 30; six colonizations on days 0, 2, 4, 7, 14, and 28; 8 or 13 colonizations following the previously described schedules. After six weeks, serum, nasal wash, BAL fluid, and small intestinal contents were harvested and analyzed by ELISA for antibody titers against TTFC (circles) or SA-TTFC (squares), as indicated.

(**C**) Same experiment as **Fig 4B**. Pet shop mice or SPF mice were colonized on either the skin or nose using TTFC conjugated to either *S. epidermidis* or *S. xylosus*. Nasal colonization was performed by pipetting 20 µL of bacterial culture normalized to $OD_{600} = 6$ into each nostril. After eight colonizations in six weeks, serum, nasal wash, BAL fluid, and small intestinal contents were collected and analyzed for the presence of TTFC-specific antibodies.

Data shown are from one (**A**) or are representative of at least two (**B**, **C**) independent experiments. Graphs show geometric mean with a 95% confidence interval. *P* values (ns, not significant; * $p < 0.05$; ** $p < 0.01$; *** $p < 0.001$) were calculated using a one-way ANOVA (**A**, **B**) or two-way ANOVA (TTFC vs. SA-TTFC comparisons in **B**, **C**) followed by Tukey's multiple comparisons test, assuming log-normal distributions. Asterisks in figures represent pairwise comparisons of the indicated group to naive mice of the corresponding group; *p* values for all comparisons are reported in **Table S1**.

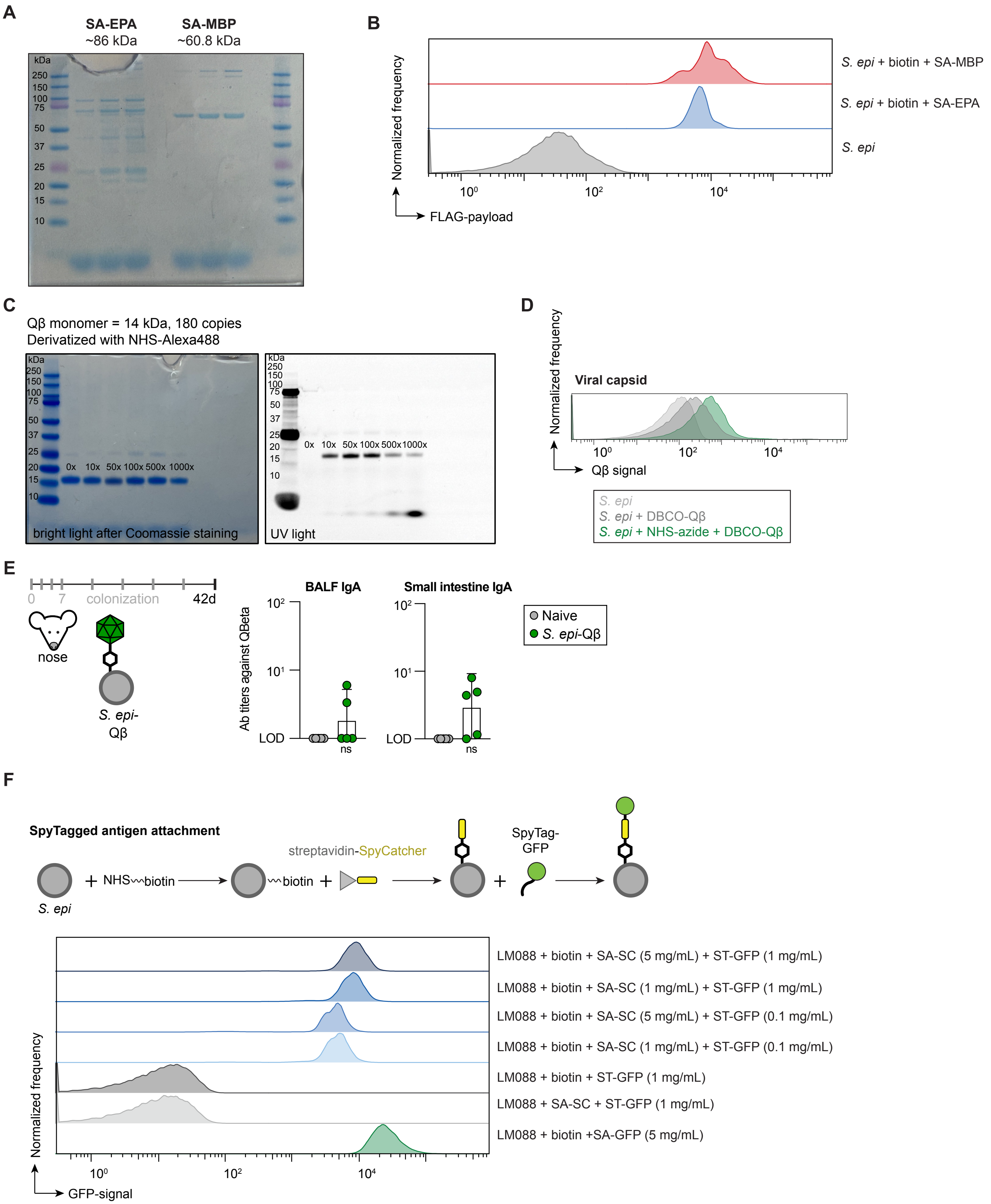
A
SA-EPA
~86 kDa
SA-MBP
~60.8 kDa
kDa
250
150
100
75
50
37
25
20
15
10
B
S. epi + biotin + SA-MBP
S. epi + biotin + SA-EPA
S. epi
Normalized frequency
FLAG-payload
C
Qβ monomer = 14 kDa, 180 copies
Derivatized with NHS-Alexa488
0x 10x 50x 100x 500x 1000x
bright light after Coomassie staining
UV light
D
Viral capsid
Normalized frequency
Qβ signal
S. epi
S. epi + DBCO-Qβ
S. epi + NHS-azide + DBCO-Qβ
E
0 7 colonization 42d
nose
S. epi-Qβ
BALF IgA
Small intestine IgA
Ab titers against QBeta
LOD
ns
Naive
S. epi-Qβ
F
SpyTagged antigen attachment
S. epi
NHS〰biotin
〰biotin
streptavidin-SpyCatcher
SpyTag-GFP
LM088 + biotin + SA-SC (5 mg/mL) + ST-GFP (1 mg/mL)
LM088 + biotin + SA-SC (1 mg/mL) + ST-GFP (1 mg/mL)
LM088 + biotin + SA-SC (5 mg/mL) + ST-GFP (0.1 mg/mL)
LM088 + biotin + SA-SC (1 mg/mL) + ST-GFP (0.1 mg/mL)
LM088 + biotin + ST-GFP (1 mg/mL)
LM088 + SA-SC + ST-GFP (1 mg/mL)
LM088 + biotin +SA-GFP (5 mg/mL)
Normalized frequency
GFP-signal

G

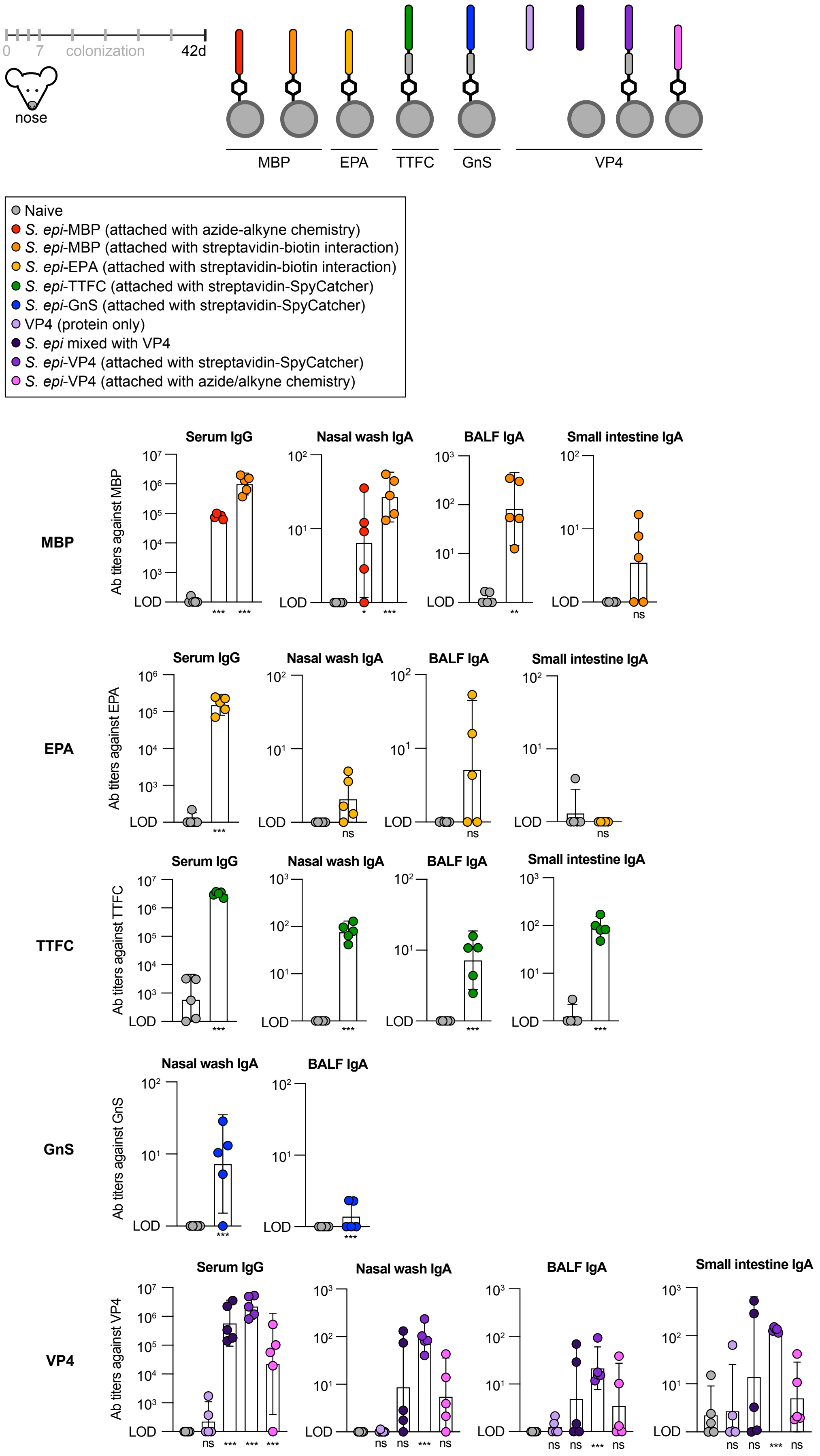

0
7
colonization
42d
nose
MBP
EPA
TTFC
GnS
VP4
Naive
S. epi-MBP (attached with azide-alkyne chemistry)
S. epi-MBP (attached with streptavidin-biotin interaction)
S. epi-EPA (attached with streptavidin-biotin interaction)
S. epi-TTFC (attached with streptavidin-SpyCatcher)
S. epi-GnS (attached with streptavidin-SpyCatcher)
VP4 (protein only)
S. epi mixed with VP4
S. epi-VP4 (attached with streptavidin-SpyCatcher)
S. epi-VP4 (attached with azide/alkyne chemistry)
Serum IgG
Nasal wash IgA
BALF IgA
Small intestine IgA
Ab titers against MBP
Ab titers against EPA
Ab titers against TTFC
Ab titers against GnS
Ab titers against VP4
LOD
ns

**Figure S15.** Commensal vaccination with clinically relevant antigens.

(**A**) Coomassie-stained gel of SA-EPA and SA-MBP proteins used for chemical conjugation.

(**B**) Flow cytometry data showing labeling of *S. epidermidis* (Syto9$^+$) with EPA and MBP *in vitro*. Antigens are detected using staining with anti-FLAG antibody.

(**c**) Coomassie-stained (left) and UV-illuminated (right) gel showing *in vitro* derivatization derivatization of Qβ viral capsid using amine-reactive chemistry *in vitro*, at indicated molar ratios of ester to Qβ subunit.

(**D**) Flow cytometry data demonstrating conjugation of whole Qβ viral capsid to *S. epidermidis in vitro*, using amine-reactive chemistry. Qβ is detected on the bacterial surface using a bacteriophage-specific antibody.

(**E**) Data shown are from the same experiment as **Fig. 4C**. At the experimental endpoint, BAL fluid and small intestinal contents were harvested, and the endpoint titers of capsid-specific antibodies in each analyte were measured by ELISA.

(**F**) Schematic and accompanying flow cytometry data validating the three-step approach for attaching SpyTag003-linked proteins to the surface of *S. epidermidis*, using a streptavidin-SpyCatcher003 splint protein intermediate.

(**G**) Data shown are from the same experiment as **Fig. 4D**. *S. epidermidis* was conjugated to a set of antigens: maltose binding protein (MBP) from *E. coli*, *Pseudomonas aeruginosa* exoprotein A (EPA), TTFC, GnS (from Rift Valley fever virus) and VP4 (from rotavirus) using the attachment strategies (or relevant protein only and mixed conditions) indicated in the legend. These conditions where then used to colonize the nose of mice. For each colonization, 10 μL of $OD_{600}$ = 18 of bacterial culture was pipetted into each nostril. After eight colonizations administered over six weeks, serum, nasal wash, BAL fluid, and small intestinal contents were collected, and the levels of antibodies specific to the indicated antigens were measured by ELISA.

Data shown are from one (**G**) or representative of at least two (**B**, **D**, **E**, **F**) independent experiments. Graphs show geometric mean with a 95% confidence interval. *P* values (ns, not significant; * $p < 0.05$; ** $p < 0.01$; *** $p < 0.001$) were calculated using Welch's t tests (**E**, anti-MBP BALF and Small intestine IgA, anti-EPA in panel **G**), ordinary one-way ANOVA followed by Tukey's multiple comparisons test (anti-MBP Serum IgG and Nasal wash IgA in panel **G**), assuming log-normal distributions, or two-way ANOVA on log-transformed data followed by pairwise testing with Šidák's correction for multiple comparisons (**G**). Asterisks in figures represent pairwise comparisons of the indicated group to naive mice; *p* values for all comparisons are reported in **Table S1**.

### METHODS

#### Bacterial growth

All *Staphylococcus* strains (including *S. epidermidis*) and *Mammaliicoccus lentus* were grown on Brain Heart Infusion (BHI) agar plates (BD Difco 241810) at 37°C overnight. Individual colonies were picked and grown overnight (16-18 hrs) at 37°C in BHI liquid media (BD Difco 237200) with shaking at 220 rpm. Genetically engineered *S. epidermidis* strains were grown under the same conditions with the addition of 10 µg/mL chloramphenicol (Sigma C1919) to all media. *Corynebacteria* strains were grown in BHI supplemented with 1% Tween80 at 37°C overnight with shaking at 220 rpm. *Deinococcus radioduran*s was grown at 30°C on either Luria-Bertani (LB, BD Difco 244510) agar plates supplemented with 1% glucose or in liquid LB (BD DIFCO 244610) + 1% glucose with shaking at 220 rpm. Growth took approximately 2-3 days. *Caulobacter crescentus* was grown for 48 hours on peptone (Sigma P5905), yeast extract (Sigma Y1625), agar plates (referred to as PYE) supplemented with 0.2 g/L $MgSO_4x7H_2O$ (Sigma 230391). A single colony was picked into liquid PYE + $MgSO_4$ and grown at 30°C with shaking at 220 rpm for an additional two days. *Vibrio natriegens* was grown on LB + 1.5% NaCl agar plates or in liquid media overnight at 30°C with shaking at 220 rpm. *Bacillus megaterium* and *Xanthomonas campestris* were grown at 30°C on LB agar or in LB liquid media with shaking at 220 rpm for 18-20 hrs. *Halomonas elongata* was grown on Difco Marine Broth 2216 (279110) agar plates or in Marine Broth at 30°C with shaking at 220 rpm. All other bacterial strains were grown under conditions described in **Table S2.**

#### Mice

Mice were housed at Stanford University in specific pathogen-free (SPF) conditions according to the guidelines established by the Stanford University Institutional Animal Care and Use Committee (IACUC). C57BL/6 (Taconic Biosciences B6-F) and B6.129S2-$H2^{dlAb1\text{-}Ea}$/J mice (The Jackson Laboratory 003584) were purchased commercially. For experiments involving microbially experienced animals (i.e. pet shop mice), mice were purchased from a local pet shop (East Bay Vivarium), screened for the absence of pathogens, and housed separately in BSL2+ rooms in accordance with the Stanford Veterinary Center and IACUC requirements. Experiments were not performed blindly; the same person handling the mice was also performing the readouts. Mice were housed in a 12/12 light/dark cycle, at room temperature (20°C to 26°C) and a humidity level of 40–65%.

#### Protein purification

*E. coli* BL21(DE3) were transformed with pET28 expression plasmids for streptavidin fusion proteins (SA-GFP, SA-TTFC, SA-MBP, or SA-SpyCatcher003) or pET30b expression plasmids for GFP-SpyTag003 or TTFC-SpyTag003. Single colonies were grown overnight shaking at 220 rpm at 37°C in 5 mL LB containing kanamycin (50 µg/mL). The next day, these cultures were used to inoculate 1 L Terrific

Broth (Thermo Scientific AAJ75856A) in a 2.8 L baffled flask with kanamycin. Cultures were grown at 37°C with shaking at 220 rpm until an optical density at 600 nM ($OD_{600}$) of ~0.4-0.6 was observed. Flasks were cooled and then induced with 1 M isopropyl-β-d-thiogalactopyranoside (IPTG) to a final concentration of 0.5 mM. Flasks were then incubated overnight at 18°C, except TTFC-SpyTag003 which was cultured overnight at 30 °C, with shaking at 220 rpm and collected by centrifugation at 4°C (3,800*g*) for 20 min. All subsequent steps were performed at 4°C and/or on ice. The cell pellet was resuspended in lysis buffer (50 mM Tris-HCl pH 7.5, 150 mM NaCl, 10 mM imidazole) plus DNase (Sigma 10104159001), lysozyme (Sigma L6876) and protease inhibitor (Sigma 11836170001). Cells were lysed by sonication with a Thermo Fisher sonicator (FB120) with a 0.63 cm probe at 40% amplitude for 5 min of 15 seconds pulse on, 45 seconds pulse off. Sonication steps were repeated 2-3 times as needed for complete lysis. Lysate was clarified by centrifuging for 1 h at 20,000*g* at 4°C and then added to 5 mL washed Ni-NTA agarose (Qiagen 30210). Protein was left to bind resin for 1 hr on a rotary shaker at 4°C, and then loaded onto a polypropylene column (Qiagen 34964). Resin was washed with 50 mL of lysis buffer and then eluted in 5-10 mL elution buffer (50 mM Tris-HCl pH 7.5, 150 mM NaCl, 50 mM imidazole). Proteins were concentrated to a volume of 2.5 mL using Amicon Ultra centrifugal filters with an appropriate molecular weight cutoff (Millipore UFC9030). Concentrated protein was then buffer exchanged using PD-10 desalting columns (Cytiva 17085101) into storage buffer (100 mM sodium phosphate, 150 mM NaCl, 10% glycerol, pH 7.5). Protein concentration was determined by nanodrop before samples were aliquoted, flash frozen, and stored at -80°C. Protein purity was assessed by SDS-PAGE gel (Invitrogen NP0336BOX) with Coomassie staining (Invitrogen LC6060 and Bio-Rad 1610374EDU).

For purification of recombinant protein from mammalian cells, Expi293 cells (Gibco A14635) were cultured in suspension in Expi293 Expression Medium (Thermo Fisher 1435102) at 37C with 8% CO2 at >80% humidity at 125 rpm on a 19mm orbital shaker. On the day of transfection, cells were split to a density of 2.5E6 cells/mL. To transfect cells, clean maxiprepped expression plasmid DNA and ExpiFectamine reagent (Gibco A14635) were separately diluted in optiMEM, quickly combined, and added to cells after a 5 minute incubation period. 16-18 hours post-transfection, ExpiFectamine 293 Transfection Enhancer reagents 1 and 2 (Gibco A14635) were combined and added to the cells. For a 100 mL transfection, cells were grown in 250 mL flasks (Corning 431144), 100 µg DNA was diluted in 5 mL optiMEM and 270 µL ExpiFectamine was diluted in 4.7 mL optiMEM; for this scale, 500 µL Enhancer reagent 1 was combined with 5 mL Enhancer reagent 2. Four days after transfection, cells were pelleted by spinning at 3500*g* for 15 min at 4°C before being resuspended in hypotonic lysis buffer (10 mM Tris, 1.5 mM $MgCl_2$, 10 mM KCl, pH 7.4), supplemented with protease inhibitor tablets (Sigma 11836170001) on ice for 30-60 minutes. Cells were lysed using a sonicator (Thermo Fisher FB505) with a 0.63 cm probe at 40% amplitude for 5 min of 15 seconds pulse on, 45 seconds pulse off. Lysate was clarified by spinning at 20,000*g* for 30 minutes at 4°C, before being passed through a 0.22 µm filter (Millipore Sigma

S2GPU05RE). Ni-NTA agarose beads (GoldBio H-350-500) was washed and added to lysate for 1 hour at 4°C to purify His-tagged proteins. Resin was washed (20 mM sodium phosphate, 500 mM NaCl, 20 mM imidazole, pH 7.4) on polypropylene columns and eluted (20 mM sodium phosphate, 300 mM NaCl, 250 mM imidazole, pH 7.4) before being concentrated using Amicon centrifugal filters and buffer exchanged using PD-10 columns, as above. Protein was assayed for purity by nanodrop measurement, Coomassie-staining of SDS-PAGE gel, and ELISA, and was flash frozen into aliquots stored at -80°C.

**RVFV GnS antigen production**

The amino acid sequence of stabilized RVFV Gn (GnS) was fused via a glycine-serine linker to a 3xFLAG, SpyTag003, and polyhistidine sequence. The nucleotide sequence was codon optimized by GenScript for mammalian expression. Expi293 cells were transiently transfected using PEI-MAX and cultured for three days at 37°C. Cell supernatant was harvested, centrifuged, and filtered. Clarified supernatants were treated with 5M NaCl (5 mL per 100 mL supernatant), 1M Tris pH 8.0 (7 mL per 100 mL supernatant), and batch bound with Nickel Sepharose Resin (Cytiva 17371202) overnight at 4°C. Next, resin was collected in a gravity column, washed with four column volumes of wash buffer containing 25 mM Tris, 150 mM NaCl, and 30 mM imidazole, pH 8.0, and then eluted with three column volumes of elution buffer comprising 25 mM Tris, 150 mM NaCl, and 300 mM imidazole, pH 8.0. Eluted proteins were subsequently purified by size-exclusion chromatography using a Superdex S200 Increase 10/300 gel filtration column into a final buffer containing 25mM Tris, 150mM NaCl, 5% Glycerol, pH 8.0. Antigenicity of the manufactured protein was evaluated by biolayer interferometry using monoclonal antibodies.

**Rotavirus VP4 expression and purification**

Stabilized rotavirus VP4 was genetically fused via a glycine-serine linker to a 3×FLAG tag, SpyTag003, and a 6×His tag. The nucleotide sequence was codon-optimized by GenScript for bacterial expression. The plasmid was transformed into BL21(DE3) competent E. coli (NEB C2527) and cultured at 37°C in Terrific Broth II containing 50 µg/mL kanamycin until an OD600 of 0.6 was reached. Protein expression was induced with 0.1 mM IPTG (isopropyl β-D-1-thiogalactopyranoside), followed by overnight incubation at 18°C. Cells were harvested by centrifugation, resuspended in lysis buffer (50 mM Tris, 250 mM NaCl, 20 mM imidazole, 100 mM arginine, 250 mM mannitol, 0.5 mg/mL lysozyme, 1 U/mL benzonase, protease inhibitor cocktail, pH 8.0), and lysed by sonication on ice. After removal of cell debris by centrifugation at 14,000 × g for 30 min, the clarified lysate was applied to Ni-NTA agarose resin (Qiagen 30210) equilibrated with wash buffer (50 mM Tris, 250 mM NaCl, 20 mM imidazole, 100 mM L-arginine, 250 mM mannitol). The resin was washed twice with 10 column volumes of wash buffer. Protein was eluted with 2.4 column volumes of elution buffer (50 mM Tris, 250 mM NaCl, 500 mM imidazole, pH

8.0) and further purified by size-exclusion chromatography using a Superdex 200 Increase 10/300 GL column (Cytiva) equilibrated with sizing buffer (25 mM Tris-HCl, 150 mM NaCl, 5% glycerol, pH 8.0). Antigenicity of the purified protein was assessed by biolayer interferometry using monoclonal antibodies.

**Protein derivatization**

Proteins used for azide/alkyne cycloaddition reactions (TTFC-SpyTag003, GFP-SpyTag003, MBP-SpyTag003, and QB sourced from Fina Biosolutions) were derivatized using EZ-link TFP-PEG4-DBCO (dibenzocyclooctyne) following manufacturer's instructions (Thermo Fisher C20043). Briefly, proteins were buffer exchanged using Cytiva PD-10 desalting columns into sodium phosphate buffer if not already stored in an amine-free buffer. 5-50 times molar excess TFP-PEG4-DBCO was added to the protein and incubated for 1 hour at room temperature. Derivatized protein was purified away from excess crosslinker using an appropriately sized Zeba dye and biotin removal column for the amount of sample (Thermo Fisher AA44296 for analytical scale, AA44298 or A44300 for *in vivo* scale). Degree of labeling was determined by measuring absorbance at 280 nm and 309 nm on a Nanodrop spectrophotometer (Thermo Fisher). Proteins were stored on ice until use.

**Bacterial labeling for *in vitro* analysis using amine or thiol reactive crosslinkers**

Overnight cultures of *S. epidermidis* were OD normalized (500 µL of $OD_{600}$ = 6) and harvested in a 1.5 mL microcentrifuge tube by centrifuging for 2 min at 9,000*g* at room temperature. Cells were washed three times with 1 mL phosphate buffered saline (PBS) to remove media components. Bacteria were resuspended in PBS and N-hydroxysuccinimide ester (NHS) reagents were used at 1 mM or 2 mM final concentration. Stock solutions were prepared immediately prior to usage.

For NHS-PEG4-biotin (Vector Laboratories QBD-10200) or NHS-PEG24-biotin (Vector Laboratories QBD-10774) labeling, bacteria were resuspended in 90 µL of PBS and 10 µL of 20 mM NHS-PEGn-biotin were added to reach a 2 mM final concentration. For NHS-PEG4-azide (Vector Laboratories QBD-10501) labeling, bacteria were resuspended in 98 µL of PBS and 2 µL of 50 mM NHS-PEG4-azide (stock solution in DMSO) were added for a final concentration of 1 mM. For direct fluorophore attachment, NHS-AlexaFluor488 (Thermo Scientific A20100) NHS-Cy5 (Vector Laboratories FP-1321-1) or maleimide-Cy5 (Vector Laboratories FP-1322-1) were used at a 1 mM final concentration. For trans-cyclooctene/tetrazine labeling, NHS-PEG4-TCO (CCT-A137) was added to reach a 1 mM final concentration. For crosslinker titration experiments, bacteria were resuspended in an appropriate volume to give the indicated final concentration of crosslinker.

Bacteria and crosslinker were vigorously mixed and incubated for 30-45 minutes at room temperature. After the incubation, bacteria were washed three times with 1 mL PBS, and then resuspended in 50 µL PBS. Streptavidin fusion proteins including SA-Cy5 (Thermo Scientific SA1011), SA-GFP, SA-

TTFC, SA-EPA, SA-MBP, SA-Qdot655 (Thermo Scientific Q10121MP), or DBCO-derivatized proteins including TTFC, MBP, and QB (Fina Biosolutions) were diluted to a 2x concentration stock solution and 50 µL were added to bacterial culture (typically 1 mg/mL final concentration). DBCO-Cy5 (Vector Laboratories CCT-A130) and tetrazine-Cy5 (CCT-1189) were used at a 50 µM final concentration. Bacteria and protein were gently mixed and then incubated for 30-45 min standing at room temperature. Protein linked bacteria were washed three times, each with 1 mL PBS and prepared for downstream applications. For detection of TTFC, MBP, and EPA, protein labeled bacteria were incubated in 100 µL of 1:100 dilution α-FLAG-PE antibody (BioLegend 637310) for 30 min at room temperature. For detection of QB, *Escherichia* phage Qbeta capsid protein antibody (Abbexa abx345659) was used at a 1:50 dilution followed by a secondary antibody, anti-rabbit IgG (H+L) polyclonal antibody conjugated to APC (Thermo Scientific A-10931), at 1:100 dilution. Following staining, bacteria were analyzed by confocal microscopy or flow cytometry.

For heat-killed *S. epidermidis*, bacteria were harvested, OD normalized, washed with PBS and then boiled at 95°C for 45 minutes. Bacterial labeling with the dead cultures then proceeded as described above.

**Bacterial labeling via noncanonical amino acid incorporation**

Single colonies of S. epidermidis were inoculated overnight in BHI supplemented with 1 mM 3-azido-D-alanine (Sigma 909823) for amino acid incorporation. After 16-18 hours of growth, cells were OD normalized (500 µL $OD_{600}$ = 6) and harvested by centrifuging 2 min at 9,000*g* at room temperature. Bacteria were washed three times with 1 mL PBS and then resuspended in 100 µL PBS. DBCO-Cy5 was added to reach a final concentration of 50 µM (Vector Laboratories CCT-A130). Labeled cells were washed an additional three times with 1 mL PBS and then used for in vitro analysis by flow cytometry.

**Nanoparticle labeling**

*S. epidermidis* overnight cultures were harvested (500 µL $OD_{600}$ = 6), washed three times with PBS and then incubated with 1 mM NHS-PEG4-azide as previously described. Surface functionalized bacteria were washed three times and then reacted with 50 µM DBCO-SpyTag003 (ordered from GenScript). After a 1 hour incubation, bacteria were washed with an excess of PBS and then mixed with SpyCatcher003-mi3 nanoparticles (0.5 mg/mL), purified as previously described[86]. The SpyTag/SpyCatcher reaction was allowed to proceed for 10 minutes, followed by immediate addition of GFP-SpyTag003. After 10 minutes, the bacterial pellet was washed three times and labeling efficiency was assessed by flow cytometry.

**Bacterial labeling for *in vivo* colonization**

Bacterial cultures (overnight growth for *S. epidermidis*, up to 48 hours for non-colonist strains) were OD normalized to 5 mL of $OD_{600}$ = 6 culture and washed with 25 mL PBS by centrifuging 3,500*g* for 8 minutes at room temperature. Cells were transferred to a 1.5 mL microcentrifuge tube with 1 mL PBS, and then washed an additional three times, 1 mL PBS each. Cells were resuspended in 900 µL PBS and NHS-PEG4-biotin was added to reach 2 mM final concentration or cells were resuspended in 980 µL PBS and NHS-PEG4-azide was added to reach 1 mM final concentration. Cells were vortexed briefly and incubated for 30-45 minutes, mixing by inversion at the halfway point. Bacteria were subsequently washed five times with 1 mL PBS each by centrifuging for 2 min at 9,000*g* at room temperature. Finally, bacteria were washed an additional five times with PBS and then resuspended in 5 mL BHI media for colonization of the skin and nose together. For experiments where a concentrated dose of labeled bacteria was applied intranasally, bacteria were resuspended in smaller volumes to reach higher OD equivalents.

For the dose matched experiment in **Figs. 3G** and **S13,** live and dead (heat-killed) *S. epidermidis* were labeled using NHS-PEG4-biotin as described, followed by three different concentrations of SA-TTFC for three doses. For the high dose condition, 1.21 mg/mL SA-TTFC was used in the labeling reaction. For the medium dose, 0.12 mg/mL SA-TTFC was used in the reaction, and at the lowest dose, 0.012 mg/mL SA-TTFC was used to label biotinylated *S. epidermidis*. For the TTFC only and *S. epidermidis* mixed with TTFC conditions, a high dose condition of 7.5 µg/mL, a medium dose of 2.5 µg/mL, and a low dose of 0.38 µg/mL were prepared in either 5 mL of BHI media or 5 mL of S. epidermidis ($OD_{600}$ = 6)

**Bacterial surface labeling with streptavidin-SpyCatcher003 splint protein**

*S. epidermidis* overnight cultures (5 mL $OD_{600}$ = 6) were labeled using NHS-PEG4-biotin as described above. The “splint protein” streptavidin-SpyCatcher003 (SA-SC) was added to biotinylated bacteria at a 1 mg/mL final concentration and incubated for 30 minutes, mixing halfway through. After washing five times with 1 mL PBS, SpyTag003 linked antigens (TTFC, GnS and VP4) were added, mixed well, and incubated for 15 minutes at room temperature, mixing halfway through. SpyTag003 linked GnS and VP4 were supplied from the laboratory of Dr. Neil King (unpublished). Labeled bacteria were washed five times with PBS and then resuspended in 1.67 mL ($OD_{600}$ = 18) and used for intranasal administration.

**Confocal microscopy of bacteria**

For experiments where labeled bacteria were imaged by confocal microscopy, cover slips were submerged in 0.1% poly-lysine (Sigma-Aldrich P8920-500ML) at room temperature for 15 minutes, before being washed twice with water and dried. 500 µL of $OD_{600}$=6 of bacterial culture was labeled before being stained in 100 µL with either MycoLight Red (AAT Bioquest 24006) or Syto9 (Invitrogen S34854), each used 1:100, at room temperature for 10 minutes. After two more washes, cells were fixed in 4% PFA in PBS for 10 minutes at room temperature while rocking. Cell pellets were then washed thrice in 1 mL PBS before being resuspended in 800 µL of PBS. Various amounts of bacterial cells were added to cover slips and bacteria were allowed to adhere for 10 minutes before being washed with PBS. Then, cover slips were mounted using ProLong Gold Antifade Mountant (Thermo Scientific P36930) and sealed with nail polish. After curing overnight at 4°C, images were acquired using a Zeiss LSM 880 confocal microscope with AiryScan detector.

**Bacterial flow cytometry**

To assess bacterial conjugation by flow cytometry, labeled bacteria were stained with either MycoLight Red JJ94 (AAT Bioquest 24006) or Syto9 (Invitrogen S34854), each used at a 1:100 dilution, for 10 minutes at room temperature before being washed twice with PBS. Bacteria were then diluted and run on a FACSymphony A5 analyzer (BD Biosciences) after passing samples through a 35 µm filter (Corning 352235). SSC threshold was lowered to detect bacteria, SSC voltage was increased, and bacteria were separated from debris by gating on MycoLight$^{+}$ or Syto9$^{+}$ events. Data were analyzed using FlowJo v.10.10.0 (BD Biosciences).

**Adoptive priming of T cells *in vivo***

On day 0, Rag2$^{-/-}$ OT-I CD45.2 (Taconic Biosciences 11490-F) or Rag2$^{-/-}$ OT-II CD45.2 donor mice (Taconic Biosciences 2334-F) were euthanized, and their spleens were harvested and smashed with the back of a syringe against a 100 µm filter. After red blood cell lysis with ACK lysis buffer (Gibco A1049201), splenocytes were washed, counted, and stained at a concentration of 2E6 cells/mL in warm PBS containing 5 µM CellTrace Violet (Invitrogen C34557) at 37°C for 20 minutes. Cells were washed after staining and then 250,000 cells were injected retro-orbitally into CD45.1 recipient SPF mice (The Jackson Laboratory 002014). On days 1 and 3, mice were colonized by swab on the head, ears, and nose using *S. epidermidis* with OT-I or OT-II epitopes either genetically expressed on or chemically conjugated to the surface of the bacteria. For streptavidin-biotin reactions, 4.5 mg/mL streptavidin-OT-I and 5.0 mg/mL streptavidin-OT-II was used, and for azide/alkyne reactions, 500 µM of each DBCO-linked peptide was used. On day 8, cervical lymph nodes were dissected from mice and homogenized by physical disruption against a 100 µm filter. The percentage of transferred cells that proliferated was analyzed using flow cytometry after surface staining for 30 minutes at 4°C with a cocktail of antibodies (all used at 1:200 final concentration except Viability Dye, which was 1:300): Fc Block (BD Biosciences 553142, clone 2.4G2), Vβ5 FITC (BioLegend 139514, clone MR9-4), Tcrb PerCP-Cy5.5 (eBioscience 45-5961-82, clone H57-597), Vα2 APC (BioLegend 127810, clone B20.1), CD45.2 AlexaFluor700 (BioLegend 109822, clone 104), eBioscience Fixable Viability Dye eFluor 780 (Invitrogen 65-0865-14), CD8a BV711 (BioLegend 100759, clone 53-6.7), CD45.1 PE (BD Biosciences 553776, clone A20), CD4 PE-Cy7 (eBioscience 25-0042-82, clone RM4-5).

**Studying antibody responses in vivo**

Six to ten week old female B6 SPF mice were used to assess the antibody response to commensals *in vivo*. For experiments involving colonizations using swabs, bacteria were normalized to $OD_{600}$=6 unless noted and resuspended in BHI media (or BHI + chloramphenicol 10 µg/mL where appropriate). Roughly 1 mL of bacteria was applied to the head, nose, and ears of each mouse using sterile swabs (Puritan 25-8062PD) dipped in culture. For experiments testing colonization at other skin sites, swabs were used to apply culture in the same way to the skin of the face or back of the mice. Where back spray is indicated, 1 mL of bacterial culture was sprayed onto the lower back of the mice using a mucosal atomization device (Amazon Business B073SV42DC), and then gently dabbed into the fur with the foam tip of the device. For intranasal administration by pipette, culture was administered slowly dropwise into each nostril. To dose culture by oral gavage, 200 µL of normalized bacterial culture was fed to each mouse using syringes attached to feeding tubes (Instech Laboratories FTP2038). When frozen stocks of bacteria were used for colonization, cultures were frozen in BHI containing 25% glycerol (Sigma-Aldrich G5516-1L) at -80°C for long-term storage and were thawed immediately before administration. When tretinoin was used as an adjuvant, tretinoin cream (Padagis or Taro) was thoroughly resuspended in BHI media at a final concentration of 100 mg/mL before being mixed with bacterial cultures and administered to mice as above. Experiments involved 8 or 13 colonizations (every other day on the first week, and then once or twice a week thereafter) administered over the course of six total weeks, unless otherwise indicated. At the experimental endpoint, mice were euthanized by $CO_2$ inhalation and blood was obtained by cardiac puncture before being spun at 3000*g* for 3 minutes in blood collection tubes (BD Biosciences 365967) and collecting serum from the supernatant. Nasal washes were acquired by first inserting a 22G catheter (Exel International 26746) affixed to a 1 mL syringe upwards into the trachea, before flushing the nasal cavity with 1 mL of PBS and collecting the resultant fluid using an Eppendorf tube positioned at the entrance of the nose. Bronchoalveolar lavage (BAL) samples were collected by inserting a cut feeding tube (Instech Laboratories FTP2038) down the trachea, tying a string around the tube and trachea to secure it, and flushing the lungs twice with 1 mL of PBS each time. Small intestinal contents were harvested by running forceps along the length of the intestinal tissue, pooled, and placed into a 2 mL microcentrifuge tube on ice. Cold PBS with protease inhibitor was added (100 mg/mL) and samples were homogenized by vortexing for 5 minutes. Samples were then centrifuged for 5 min at 8000g at 4°C to pellet debris and the supernatant was transferred to a new tube for downstream analysis. To quantify CFUs recovered from mouse skin after colonization, one ear of a mouse was excised and placed into a sterile 2 mL Eppendorf tube with 500 µL sterile room temperature PBS before being vortexed for 1 minute. Supernatant was filtered through a strainer (Stellar Scientific TC70-SWM-100) placed on top of a sterile 1.5 mL Eppendorf tube. 500 µL PBS was then added to the remaining ear pieces, and they were vortexed once more for 1 minute. The remainder of the supernatant was then added to the strainer, and bacteria were isolated by centrifugation at 14000*g* for 3 minutes at room temperature. The cell pellet was resuspended in 100 µL of

PBS and serially diluted 10-fold down a column of a 96-well plate. Finally, 5 µL from each well was applied in triplicate to agar plates containing the appropriate media, spots were allowed to dry, and resultant CFUs were then enumerated after bacterial growth.

**ELISAs**

For ELISAs run against protein antigen, high-binding plates (Corning 9018) were coated with 0.15 or 0.25 µg of recombinant protein per well diluted in 50 µL of PBS, and left to bind overnight at 4°C. For ELISAs run against whole bacteria, ELISA plates were coated with 100 µL of 0.01% poly-L-lysine (Sigma-Aldrich P4707-50ML) for 15 minutes at room temperature before being washed three times with PBS and dried. Overnight cultures of bacteria were washed two times with PBS. Then, 50 µL of $OD_{600}$=6 normalized bacterial cultures diluted in PBS were used to coat each well of the plate. The next day, plates were washed five times with PBS containing 0.05% Tween20 (Sigma-Aldrich P1379) (PBST), and each well was blocked with 200 µL of 2% BSA (Sigma-Aldrich A7030-500G) in PBST for 2 hours at room temperature. Plates were then incubated with the analyte to be assayed, with each sample diluted serially down a single column of the plate in 2% BSA in PBST. Sera were diluted 1:100 to start, with serial dilutions of either 1:4, 1:8, or 1:12 to ensure complete dilution to background levels, depending on the experiment. Nasal wash, BAL fluid, and small intestinal samples were used undiluted to start, with serial dilutions either 1:3 or 1:4 down each column of the plate. After 1 hour incubation at room temperature, plates were washed five times with PBST and incubated with either anti-mouse IgG (SouthernBiotech 1030-05) or anti-mouse IgA (SouthernBiotech 1040-05) secondary antibody for 1 hour at room temperature. Finally, after five more washes with PBST, plates were revealed using 100 µL total of TMB substrate reagents (BD Biosciences 555214) and stopped after 30 minutes with 50 µL of stop solution (BioLegend 432001). Absorbance values at 450 nm were immediately recorded. The antibody titers for each sample were calculated by logarithmic interpolation of the dilutions after normalizing to a blank, using an absolute threshold of 0.2, as previously described[87].

**Intranasal dosing of Evans blue dye**

A mixture of 0.02% Evans blue dye was added to $OD_{600}$=6 normalized S. epidermidis culture in BHI media. Indicated volumes of dye were pipetted into each nostril of mice that were either awake or anesthetized with isoflurane. After a few minutes, mice were euthanized by $CO_2$ inhalation and the trachea and lungs were dissected and photographed.

**Antigen quantification with beads**

To quantify the amount of antigen attached per bacterial cell, *S. epidermidis* LM088 was conjugated to streptavidin-TTFC at the scale described above for *in vivo* experiments, with antigen amounts corresponding to each of the three doses to be used *in vivo*. Then 50 µL of $OD_{600}$=6 labeled culture was aliquoted into each well of a 96-well plate, where it was washed with PBS, with spins performed at 3400g for 8 minutes at room temperature. Bacteria were stained in 50 µL final volume with 1:10 anti-FLAG PE antibody (BioLegend 637310, clone L5) and 1:1000 Syto9 (Invitrogen S34854) for 30 minutes at room temperature; these conditions were determined based on an experiment where we titrated the concentration of antibody and cells to ensure that staining occurred to saturation. After two washes with PBS, bacteria were analyzed by flow cytometry as described above. The median fluorescence intensity from each bacterial peak was compared to the fluorescence of beads from a commercial kit, which bear known numbers of antibody-binding sites (Bangs Laboratories 817). To prepare the beads for flow cytometry, one drop from each tube provided (which includes one unstained bead sample and four bead samples each with different antibody-binding capacity) was mixed with 50 µL of PBS and 1:20 of anti-FLAG PE antibody. After 2 washes with 1 mL PBS (spins performed at 2500g for 5 minutes at room temperature), beads were resuspended in PBS and flow cytometry was performed as above. Interpolation to determine the number of FLAG-containing streptavidin-TTFC molecules per bacterial cell was performed using the calculator provided with the kit from the manufacturer.

**Analysis of germinal center B cells at lymphoid sites**

For experiments involving analysis of germinal center B cells, various secondary lymphoid organs were dissected at the appropriate experimental endpoint. Cervical lymph nodes harvested were identified as the most lateral skin-draining node of the cervical chain. NALT was dissected by first disarticulating the lower jaw of mice, then using a scalpel to cut around the perimeter of the palate on the roof of the mouth before gently peeling off the entire intact palate. All organs were immediately placed into ice-cold PBS with BSA and EDTA (PBE) prepared by filter sterilizing autoMACS Rinsing Solution (Miltenyi Biotech 130-091-222) supplemented with 0.5% BSA (Miltenyi Biotech 130-091-376). Then, organs were smashed using the back of a syringe on a 100 µm cell strainer (Miltenyi Biotec 130-110-917) and transferred to 96-well plates (Corning 3799) for staining. Dissociated cells were washed and incubated for 30 minutes at 4°C with a cocktail of antibodies for surface staining (all 1:200 final concentration except Viability dye, which was 1:300): Fc Block (BD Biosciences 553142, clone 2.4G2), CD19 PerCP-Cy5.5 (BioLegend 115533, clone 6D5), CD38 APC (BioLegend 102711, clone 90), CD45.2 AlexaFluor700 (BioLegend 109822, clone 104), eBioscience Fixable Viability Dye eFluor 780 (Invitrogen 65-0865-14), IgG1 BV605 (BD Biosciences 742477, clone X56), IgA PE (Invitrogen 12-4204-83, clone mA-6E1), Fas PE-Cy7 (BD Biosciences 557653, clone Jo2). Stained cells were washed in cold PBE and prepared for flow cytometry by passing them through a 35 µm filter (Corning 352235). When flow cytometry could not be run on the same day, cells were fixed in BD Cytofix/Cytoperm (BD Biosciences 554722) according to the manufacturer's instructions.

Flow cytometry data were acquired on a FACSymphony A5 analyzer (BD Biosciences), and data were analyzed using FlowJo v.10.10.0 (BD Biosciences).

**Using fluorescent bacteria as a staining reagent**

Overnight cultures of the indicated bacterial species were normalized to 500 µL $OD_{600}$=6, washed thrice with 1 mL PBS, and then stained for 30 minutes at room temperature using 1 mM final concentration of NHS-AlexaFluor488 (Invitrogen A20100). Cells were then washed five times with 1 mL PBS before being resuspended in 500 µL PBS. During the surface staining step of cell suspensions for 30 minutes at 4°C, as indicated above, the indicated volumes of fluorescent bacteria were added to the staining mix, before washing mammalian cells. 1/8 of a cervical lymph node prep stained with 0.0625 µL of $OD_{600}$=6 bacteria per well was found to give appropriate signal above background, which corresponds to approximately a 4:1 ratio of mammalian cells to bacterial cells. For experiment testing blocking of fluorescent bacterial staining with excess unlabeled bacteria, 1.25 µL of $OD_{600}$=6 unlabeled bacteria were used per well. Mice expressing an unrelated BCR carried the IghelMD4 transgene and were purchased commercially (The Jackson Laboratory 002595). Flow cytometry data acquisition and analysis was performed as described above.

**Imaging of mouse NALT**

NALT was dissected as above by peeling the entire intact palate off the roof of the mouth. Tissues were flash frozen in Optimal Cutting Temperature compound (OCT) and then sectioned using a Leica cryostat to produce sections of 100 µm thickness on slides. Slides were then washed thrice with PBS at room temperature for 5 minutes each, before being outlined using an ImmEdge pen and dried. Sections were fixed with 2% PFA at room temperature for 10 minutes. Tissues were washed three times in PBS for 5 minutes each at room temperature and then permeabilized with 0.1% Triton X-100 in PBS for 10 minutes at room temperature. After an additional set of washes with PBS for 5 minutes, slides were blocked (blocking buffer contains 10% FBS, 1% BSA, 0.1% Triton X-100, and 0.01% $NaN_3$ in PBS) for 1 hour at room temperature. Then, sections were stained overnight at 4°C in blocking buffer with either UEA-1 lectin DyLight 594 (1:200; Vector Laboratories DL-1067-1), anti-EpCAM (1:100, Thermo Scientific 14-5791-81, clone G8.8), anti-B220 (1:100, BD Biosciences 553084, clone RA3-6B2), anti-MHCII AlexaFluor594 (1:100, Biolegend 107650, clone M5/114.15.2), or anti-GP2 AlexaFluor488 (1:200, MBL Life Science D278-A48, clone 2F11-C3). The next day, slides were washed three times at room temperature in blocking buffer before being incubated with DAPI (1 µg/mL final concentration) and, if primary antibodies were unlabeled, donkey anti-rat IgG AlexaFluor594 secondary antibody (1:250; Invitrogen A-21209) for 1 hour at room temperature. One last set of three washes in blocking buffer for 5 minutes was performed before slides were dried and mounted in ProLong Gold Antifade Mountant (Thermo Scientific P36930) and sealed with nail polish. After curing overnight at 4°C, images of sections were acquired using a Zeiss LSM 880 confocal microscope.
